# A GenoChemetic strategy for derivatization of the violacein natural product scaffold

**DOI:** 10.1101/202523

**Authors:** Hung-En Lai, Alan M. C. Obled, Soo Mei Chee, Rhodri M. Morgan, Rosemary Lynch, Sunil V. Sharma, Simon J. Moore, Karen M. Polizzi, Rebecca J. M. Goss, Paul S. Freemont

## Abstract

Natural products and their analogues are often challenging to synthesise due to their complex scaffolds and embedded functional groups. Solely relying on engineering the biosynthesis of natural products may lead to limited compound diversity. Integrating synthetic biology with synthetic chemistry allows rapid access to much more diverse portfolios of xenobiotic compounds which may accelerate the discovery of new therapeutics. As a proof-of-concept, by supplementing an *Escherichia coli* strain expressing the violacein biosynthesis pathway with 5-bromo-tryptophan *in vitro* or tryptophan 7-halogenase RebH *in vivo*, 6 halogenated analogues of violacein or deoxyviolacein were generated, demonstrating promiscuity of the violacein biosynthesis pathway. Furthermore, 20 new derivatives were generated from 5-brominated violacein analogues via Suzuki-Miyaura cross-coupling reaction directly using the crude extract without prior purification. Herein, we demonstrate a flexible and rapid approach to access diverse chemical space that can be applied to a wide range of natural product scaffolds.

## Introduction

Total synthesis of natural products and their analogues has often been challenging and costly due to their structural complexities (Kirschning and Hahn, 2012). However, greater chemical space may be accessed through utilisation of synthetic chemistry methodologies and reagents compared to employing enzyme-catalysed biosynthesis alone. Combining biosynthesis with chemical synthesis represents a powerful approach for rapidly generating new analogues of complex natural products and libraries of chemical derivatives suitable for screening and structure-activity relationship (SAR) assays (Eichner et al., 2012; Goss et al., 2012), leading to discovery of new and more potent compounds. The generation of such analogues can enable improvement in bioactivity and bioavailability, as illustrated, for example, by analogues of the ‘last-resort’ antibiotic vancomycin, with >200-fold improvement of potency compared to vancomycin in resistant *Enterococci* strains (Okano et al., 2017). In another study, the addition of a sterically unhindered primary amine group to a Gram-positive antibiotic deoxynybomycin expanded its antimicrobial activity to multi-drug-resistant Gram-negative pathogenic strains (Richter et al., 2017).

As a proof-of-concept, we have focused on the violacein biosynthetic pathway. Violacein (**1**), a violet pigment first isolated from the bacterium *Chromobacterium violaceum*, is part of the bisindole biosynthetic family that utilises l-tryptophan as the starting substrate. The violacein pathway is encoded within a conserved operon of five genes (*vioABCDE*), whose gene products catalyse a 14-electron oxidative biosynthesis pathway (Balibar and Walsh, 2006). The bisindole biosynthetic pathways have attracted considerable interest because of their therapeutic potential for medical applications, including antimicrobial, antiviral, trypanocidal and antitumorigenic properties (Durán et al., 2012). The violacein biosynthetic pathway also produces deoxyviolacein (**2**) as a by-product, and the coloured properties of violacein and deoxyviolacein make them interesting targets for natural product pathway engineering, such as promoter library screening (Jones et al., 2015), CRISPR-based multiplex transcriptional regulation (Zalatan et al., 2015) or diverting pathway flux via RBS engineering (Jeschek et al., 2016).

A study on oxyviolacein, a hydroxylated analogue of violacein generated by feeding exogenous 5-hydroxy-l-tryptophan (Hoshino and Ogasawara, 1990), showed bioactivity against a collection of pathogenic bacteria and fungi strains (Wang et al., 2012) suggesting that violacein analogues may be a good starting point for developing more potent antibiotics. A similar rationally designed precursor-directed biosynthetic strategy has also showed considerable success in generating analogues of flavonoids (Kufs et al., 2020). Other examples of using altered biosynthetic pathways for analogue generation, include using various enzyme homologues from bisindole pathways (Du and Ryan, 2015; Sánchez et al., 2005), interconversion of the biosynthetically related novobiocin and clorobiocin aminocoumarin antibiotics (Eustáquio et al., 2004), introduction of the halogenase from hormaomycin pathway to generate clorobiocin analogues (Heide et al., 2008), the generation of chlorinated monoterpene indole alkaloids (Runguphan et al., 2010), and chlorinated and brominated analogues of the antibiotic pacidamycin (Deb Roy et al., 2010). Although total chemical synthesis of violacein (Petersen and Nielsen, 2013; Wille and Steglich, 2001) and some substituted analogues (Haun et al., 1992; McLaughlin et al., 2014) have been reported, these syntheses are challenging and low yielding. Pathway manipulation could represent a more flexible, sustainable and rapid approach for generating violacein analogues. In this study, we have applied such approaches to generate 26 new violacein or deoxyviolacein analogues via a combination of pathway engineering enabling enzymatic halogenation of the starting substrate tryptophan, feeding brominated tryptophan, and further derivatization using Suzuki-Miyaura cross-coupling directly in crude extracts (**Figure 1**).

**Figure 1.**
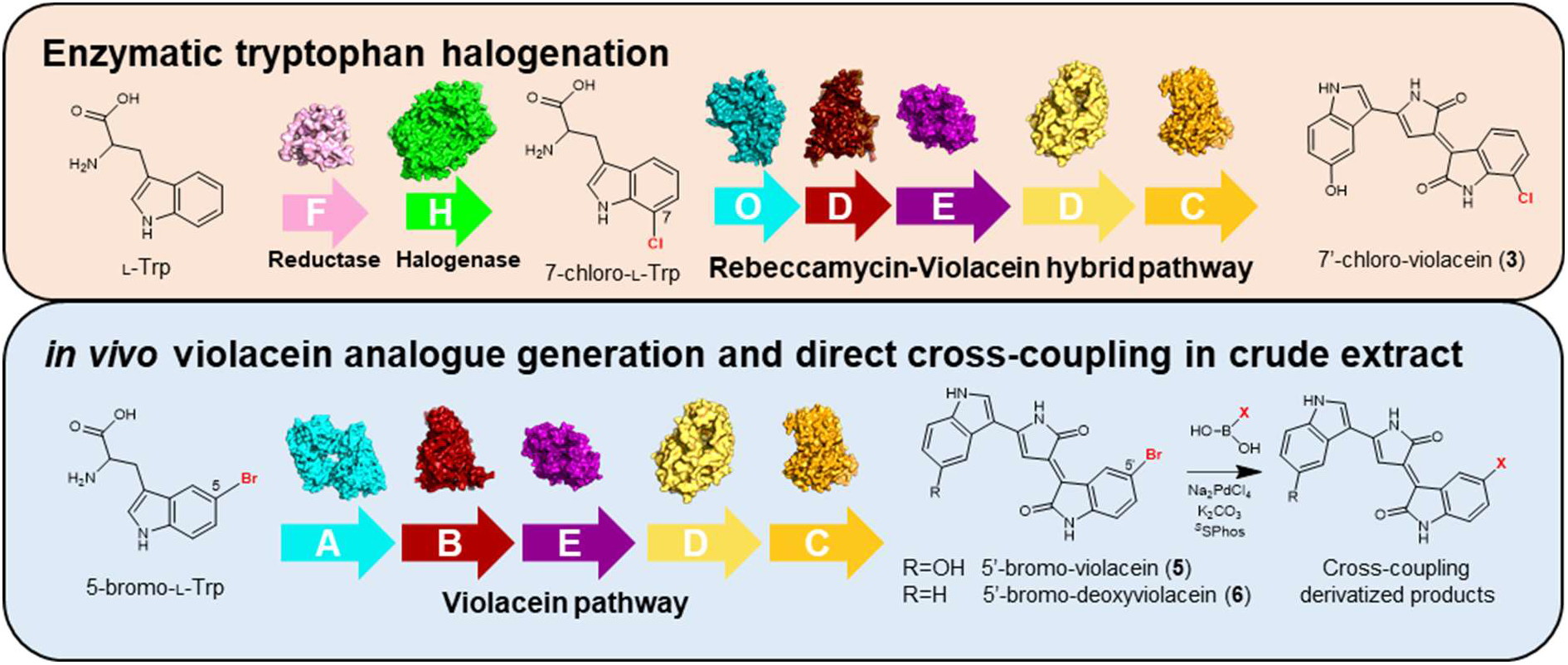
Schematics of GenoChemetic approach used in this study. Analogues of violacein (**1**) or deoxyviolacein (**2**) can be generated via *in vivo* halogenation of tryptophan by expressing flavin reductase RebF and tryptophan halogenase RebH with the rest of violacein pathway. Further derivatization via Suzuki-Miyaura cross-coupling lead to new synthetic analogues. Available enzyme structures are shown and correspond to VioA, (PDB <u>6G2P</u>); VioD (PDB <u>3C4A</u>); VioE (PDB <u>3BMZ</u>); RebH (PDB <u>2E4G</u>) as well as models generated *in silico* by Phyre2 (http://www.sbg.bio.ic.ac.uk/phyre2) for RebO, RebD, VioB, VioC and RebF.

## Results and Discussion

### VioA tryptophan oxidase accepts various tryptophan substrate analogues

VioA, the first enzyme in the violacein biosynthesis pathway, oxidises the substrate l-tryptophan to form the first pathway intermediate indole-3-pyruvic acid imine (IPA imine). In order to determine whether VioA would accept different tryptophan analogues as substrates, we purified VioA (**Figure S1**) and carried out VioA enzyme kinetic assays against various substrate analogues including l-tryptophan (TRP), 4-fluoro-dl-tryptophan (4FT), 5-methyl-dl-tryptophan (5MeT), 6-fluoro-dl-tryptophan (6FT) and 7-methyl-dl-tryptophan (7MeT). The kinetics of VioA against l-tryptophan (k_cat_ = 3.04 s^−1^, 95% CI [2.17 to 5.33]; K_M_ = 447 *μ*M, 95% CI [255 to 977], k_cat_/K_M_ = 6.80 s^−1^ mM^−1^) have been previously determined with both UV-monitored substrate depletion and coupled peroxidase assays, although only the kinetic data generated from the substrate depletion assay was reported (k_cat_ = 3.38 ± 0.32 s^−1^, K_M_ = 31 ± 11 *μ*M) (Balibar and Walsh, 2006). The discrepancy between the reported values and our data could be due to the difference in assay conditions, kinetic models chosen and the multiple reaction steps involved in coupled peroxidase assay. Among the substituted tryptophan substrates, 6FT exhibited the highest k_cat_/K_M_ value (13.9 s^−1^ mM^−1^), followed by 7MeT (11.2 s^−1^ mM^−1^), 5MeT (3.18 s^−1^ mM^−1^) and 4FT (less than 0.0658 s^−1^ mM^−1^) (**Figure 2A-C**). We cannot estimate the k_cat_/K_M_ value of VioA against 4FT accurately because the best-fit value of K_M_, 8.307 mM, is beyond the range of substrate concentration tested (up to 5 mM). Nonetheless, this shows that VioA exhibit much higher activity against 6FT compared to other substituted tryptophan analogues tested, indicating a strong preference at the 6-substituted position. Interestingly, our data shows that VioA has only 47% relative activity against 5MeT compared to L-tryptophan, in contrast to previous characterisation (Füller et al., 2016), but this is perhaps due to difference in kinetic models chosen for our data (substrate inhibition) as opposed to Michaelis-Menten in the previous study.

**Figure 2.**
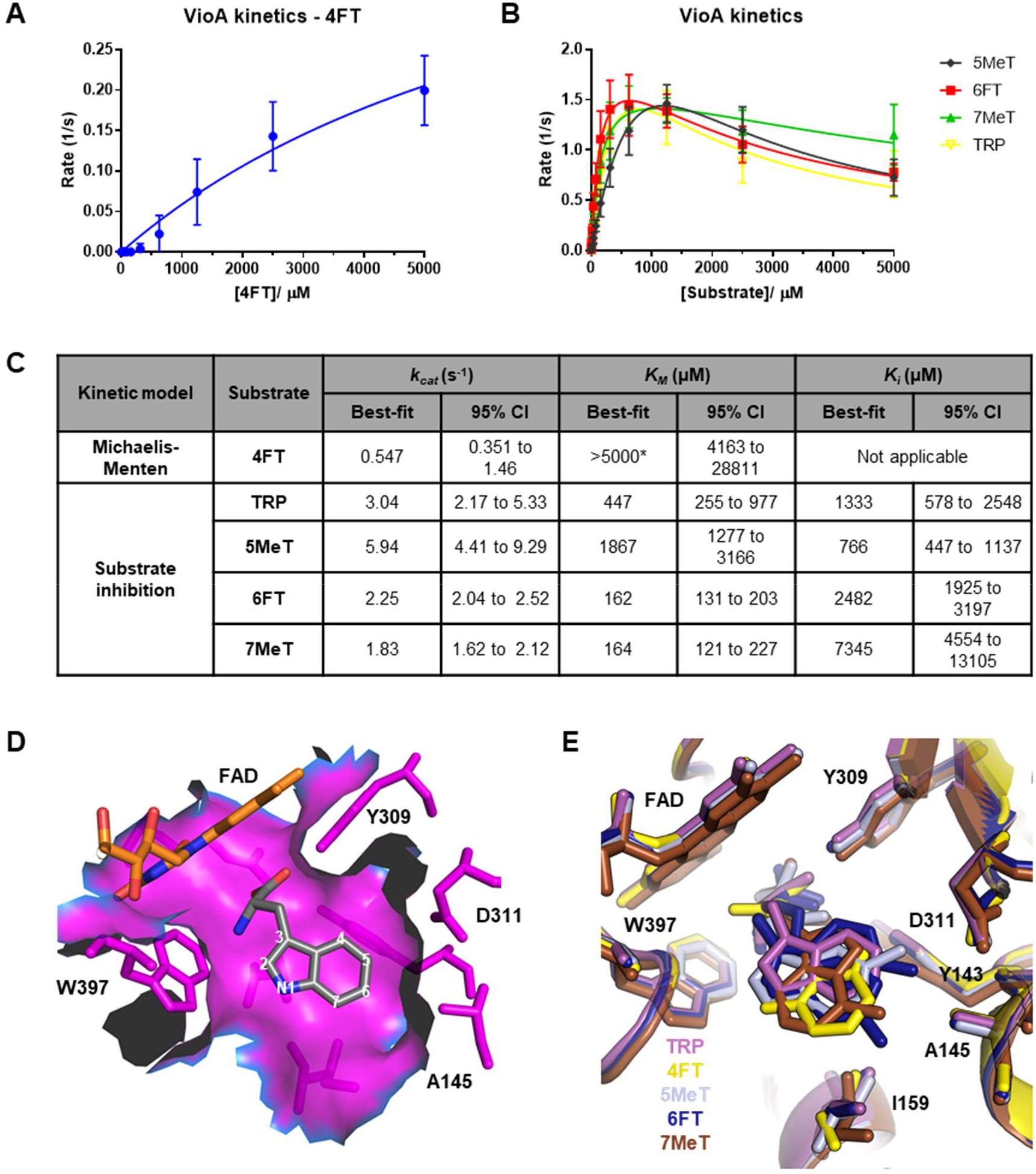
VioA substrate kinetics and structure. See also Figures S1 and S2, and Table S1. **(A)** Michaelis-Menten model is fitted on the VioA kinetics data against 4-fluoro-dl-tryptophan (4FT). **(B)** Substrate inhibition model fitted on VioA kinetics data against l-tryptophan (TRP), 5-methyl-dl-tryptophan (5MeT), 6-fluoro-dl-tryptophan (6FT) and 7-methyl-dl-tryptophan (7MeT). Data represents mean and error bars represent SD of three independent experiments carried out on three separate days. **(C)** Parameters of VioA kinetics models fitted in **A** and **B**, where best-fit values and 95% CI of *k_cat_*, *K_M_* and *K_i_* are calculated using GraphPad Prism 9.0.1 software. \**K_M_* value of 4FT is estimated to be 8307 μM which is beyond the range of substrate concentration tested (up to 5000 μM). **(D)** Binding site pocket showing space enclosed by residues (magenta) around l-tryptophan (grey with elemental colour). The carbon positions of tryptophan indole ring are labelled. **(E)** Superimposition of active sites of VioA structures complexed with l-tryptophan (TRP, magenta), 4-fluoro-l-tryptophan (4FT, yellow), 5-methyl-l-tryptophan (5MeT, light grey), 6-fluoro-ltryptophan (6FT, dark blue) and 7-methyl-l-tryptophan (7MeT, brown) in stick representation. Side chains surrounding the tryptophan ligand are labelled. Note that 6FT has 2 conformers fitted in the density map (see **Figure S2**).

### VioA ligand-soaked structures show conserved active site conformation

Although VioA has shown to be receptive towards various tryptophan analogues, there has been no wild-type VioA structure complexed with the native substrate tryptophan or its substituted analogues to date. A previous study elucidated structure of VioA complexed with a competitive inhibitor 2-[(1H-indol-3-yl)methyl]prop-2-enoic acid (IEA) (PDB **<u>5G3U</u>**), and generated a collection of VioA active site mutants to probe their substrate promiscuity (Füller et al., 2016). However, these VioA mutants exhibited decreased activity with most tryptophan analogues tested compared to wild-type VioA, demonstrating the difficulty in rationally engineering VioA towards increased substrate promiscuity. Another study reported the structure of VioA C395A mutant (PDB **<u>5ZBD</u>**) which demonstrated improved thermostability but no significantly improved activity (Yamaguchi et al., 2018).

To determine if the substrate kinetics differences were due to alterations in substrate binding, we next solved the crystal structures of apo VioA (PDB **<u>6ESD</u>**) and VioA complexed with TRP (PDB **<u>6G2P</u>**), 4FT (PDB **<u>6FW7</u>**), 5MeT (PDB **<u>6FW8</u>**), 6FT (PDB **<u>6FW9</u>**) and 7MeT (PDB **<u>6FWA</u>**) with resolutions ranging from 2.4 Å to 3.0 Å (**Figure 2D, 2E**, data statistics in **Table S1**). Our apo VioA structure is nearly identical to the published apo VioA structure (PDB **<u>5G3T</u>**, RMSD = 0.940 Å), whereas our tryptophan-bound structure (PDB **<u>6G2P</u>**) is also very similar to both inhibitor-bound VioA (PDB **<u>5G3U</u>**, 0.586 Å) and tryptophan-bound C395A VioA mutant (PDB **<u>5ZBD</u>**, 0.689 Å). The four substituted tryptophan substrate-bound VioA structures are also virtually identical to the tryptophan-bound VioA structure, with RMSD in the range of 0.304 Å to 0.419 Å. The active site of VioA reveals several key residues surrounding the C4- to C7-positions of tryptophan substrate indole ring. Of the four positions, C4 and C7 have the most space around the binding site pocket compared to C5 and C6 due to Asp311 and Ala145 side chains (**Figure 2D**), suggesting that VioA would have the lowest affinity for tryptophan analogues with substituent groups at the C5- and C6-positions. However, the interatomic distances between respective Cα atom or amine nitrogen atom and N5 of FAD cofactor is conserved for all the analogues (between 3.93 Å to 4.69 Å for Cα atom, 3.62 Å to 5.63 Å for amine nitrogen) and is similar to the TRP-bound structure (3.93 Å for Cα atom and 5.07 Å for amine nitrogen) (**Figure S2**). This showed that all the analogues can bind VioA in a catalytically active, but slightly different conformation. Interestingly, the hydrogen bonding between Arg64 and Tyr309 and the carboxylate group of all substrate analogues are conserved, as are the positions of the other active site residues (**Figure 2E**). Of the four structures with tryptophan analogues, 4FT-bound VioA structure has the lowest resolution at 3.0 Å and weakest density around the 4FT ligand compared to the other structures (**Figure S2**), suggesting lower occupancy binding which also correlates with the observed poorer enzyme kinetics compared to TRP. In addition, we observe two alternative conformers for the bound 6FT ligand demonstrating further conformational flexibility of tryptophan analogue binding. In summary, our structural analyses show that a variety of tryptophan analogues can bind VioA in different conformations, although the positions of active site residues and associated hydrogen bonding patterns are similar to TRP-bound VioA. Our data clearly show that the VioA active site can flexibly accommodate a variety of tryptophan substrate analogues for catalysis.

### Hybrid violacein-rebeccamycin biosynthetic pathway leads to 7-chlorinated analogues of violacein and deoxyviolacein

Recently, we have generated a wide range of violacein and deoxyviolacein analogues from *E. coli* cells that were expressing a synthetic violacein biosynthesis pathway *vioABCDE* with added substituted tryptophans, and these crude extracts were tested against malarial parasite *Plasmodium falciparum* (Wilkinson et al., 2020). Due to the promiscuity of the enzymes involved in the biosynthesis of violacein, we rationalized that a combination of enzymes from closely related bisindole biosynthesis pathways could generate compatible tryptophan analogues leading to other interesting new-to-nature analogues of violacein that might be otherwise difficult to synthesise. Rebeccamycin is a closely related bisindole where the biosynthetic pathway contains several genes including *rebO* and *rebD* which are homologous to *vioA* and *vioB* respectively. In addition, the pathway also consists of a tryptophan 7-halogenase RebH and its associated flavin reductase RebF that are responsible for generating 7-chloro-l-tryptophan from l-tryptophan via a chloraminelysine intermediate (Yeh et al., 2005, 2007). As a proof-of-principle, we combined the rebeccamycin (Reb) and violacein (Vio) biosynthetic operons to generate C7-chlorinated violacein analogues, which we were able to successfully produce in *E. coli* cells (**Figure 3A, 3B**) identified via LC-HRMS/MS (**Figure 3C**, **Table S2**). We found that although the RebOD+VioCDE strain (lacking RebFH) did not produce any chlorinated analogues of violacein, it was able to functionally substitute VioAB to produce violacein and deoxyviolacein (Balibar and Walsh, 2006). Next, by comparing RebFH+VioABCDE and RebFHOD+VioCDE (where RebOD replaced VioAB) strains, we observed that RebOD increased the proportion of chlorinated analogues from 6% to 33% when compared with VioAB (**Figure 3D, 3E**). This was likely due to the fact that RebO has k_cat_/K_m_ value 57-times higher for 7-Cl-l-tryptophan than that of l-tryptophan (Nishizawa et al., 2006), thus RebO and RebD preferentially accept the 7-chloro analogue of l-tryptophan and IPA imine respectively, producing the 7-chlorinated intermediate for downstream enzymes in the violacein biosynthesis pathway. By replacing the weak promoter J23114 for RebF and RebH with a medium strength promoter J23108, we also observed a total analogue increase of nearly 30% (**Figure 3F**). This shows that it is possible to fine tune the proportion of chlorinated analogue by changing the promoter strength of pathway enzymes, and optimization of fermentation protocols such as length of incubation, temperature, media conditions might further increase both proportion and the total amount of chlorinated analogues (Jones et al., 2015).

**Figure 3.**
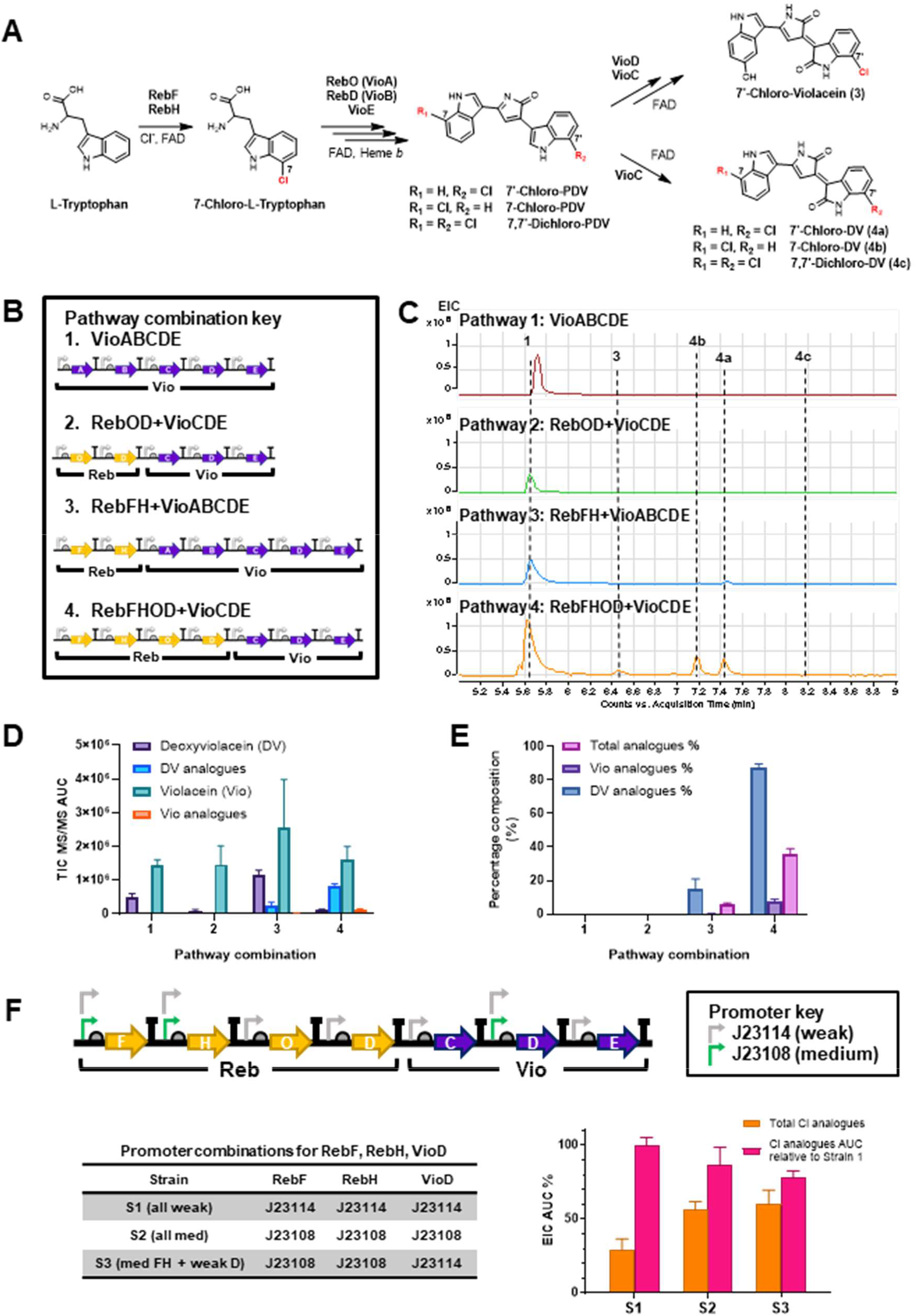
Generation of 7-chloro analogues of violacein and deoxyviolacein via in vivo halogenation. See also Table S2. **(A)** Schematic of the hybrid rebeccamycin-violacein pathway enzymes that lead to 7-chloro analogues of violacein (**3**) and deoxyviolacein (**4a-c**), as analysed by LC-HRMS/MS (**Table S2**). **(B)** Four constructs constituting various combinations of violacein and/or rebeccamycin biosynthetic genes to generate chlorinated violacein analogues. Pathway symbols were drawn using SBOL visual standard for biological parts. **(C)** Representative extracted ion chromatograms (EIC) targeting [M+H]^+^ m/z of **1**, **3**, **4a-c** detected from ethanol extracts of *E. coli* cells harbouring one of the pathway combinations in **(B)**. Compounds detected are shown with dotted line to indicate approximate retention times. Compound **2** is not shown here as it elutes at the same retention time as **3**. VioABCDE sample was run on a separate day so the retention time of **1** is slightly later than the other three samples. **(D)** Area under curves of m/z species extracted at the MS/MS level from TIC of ethanol extracts from cells harbouring respective pathway combination in **B**. Vio analogue refers to **3**, whereas DV analogues refer to **4a-c**. **(E)** Percentage compositions of Vio and DV analogues from different *E. coli* strains expressing hybrid Reb/Vio pathway variants. Data shows the mean and SD of biological duplicates. **(F)** Percentage compositions of 7-chloro analogues of violacein and deoxyviolacein extracted from *E. coli* expressing RebFHOD+VioCDE with different combinations of medium (J23108, green) and weak (J23114, grey) promoters controlling the expression of RebF, RebH and VioD. Data shows the mean and SD of biological triplicates.

### Generation of cross-coupling derivatives directly in brominated violacein crude extracts

Further derivatisation, through chemical cross-coupling, of new-to-nature, halogenated natural product analogues generated by a synthetic biological strain, was first demonstrated with pacidamycins (Deb Roy et al., 2010). We applied a similar strategy by first feeding 5-bromo-dl-tryptophan to *E. coli* cells expressing VioABCDE to generate 5’-bromo-violacein and 5’-bromo-deoxyviolacein which were detected by LC-HRMS/MS (**Table S2**). We then proceeded to screen a series of conditions that might facilitate Suzuki-Miyaura cross-coupling. However, due to low titres of 5’-bromo-violacein and 5’-bromo-deoxyviolacein available in the crude extract, and challenges relating to the separation of these compounds that showed propensity for pi-stacking, isolation of products for further characterisation and bioassays of the cross-coupling product proved difficult. We thus focused on systemically identifying conditions that would allow direct cross-coupling in crude extracts which identified optimal conditions for derivatization. The corresponding cross-coupling products 5’-(*p*-tolyl)-violacein (**7**) and 5’-(*p*-tolyl)-deoxyviolacein (**8**) were detected in the cross-coupling mixture by LC-HRMS/MS (**Figure 4A**, **Table S2**). Having identified the optimised conditions for cross-coupling at low concentrations, we performed cross-coupling reaction with a selection of 15 aryl boronic acids chosen to sample a wide range of steric and electronic variations, which gave a total of 20 new cross-coupled analogues of violacein and deoxyviolacein (**Figure 4B**). We observed that while boronic acids with electron-donating groups (*e.g. p*-methoxyphenylboronic acid) gave better yield and MS/MS data were obtained, for those with electron-withdrawing groups (*e.g. p*-formylphenylboronic acid), only MS data were obtained due to lower conversion and more difficult electrospray ionisation of corresponding violacein analogues (**Table S2**). Pleasingly, even boronic acid pinacol esters were successfully converted into the corresponding violacein product (*e.g. p*-cyanophenylboronic acid pinacol ester). In summary, the optimised conditions showed that a variety of cross-coupling analogues could be synthesised without the need for purification prior to cross-coupling, allowing for rapid access to potential cross-coupling products.

**Figure 4.**
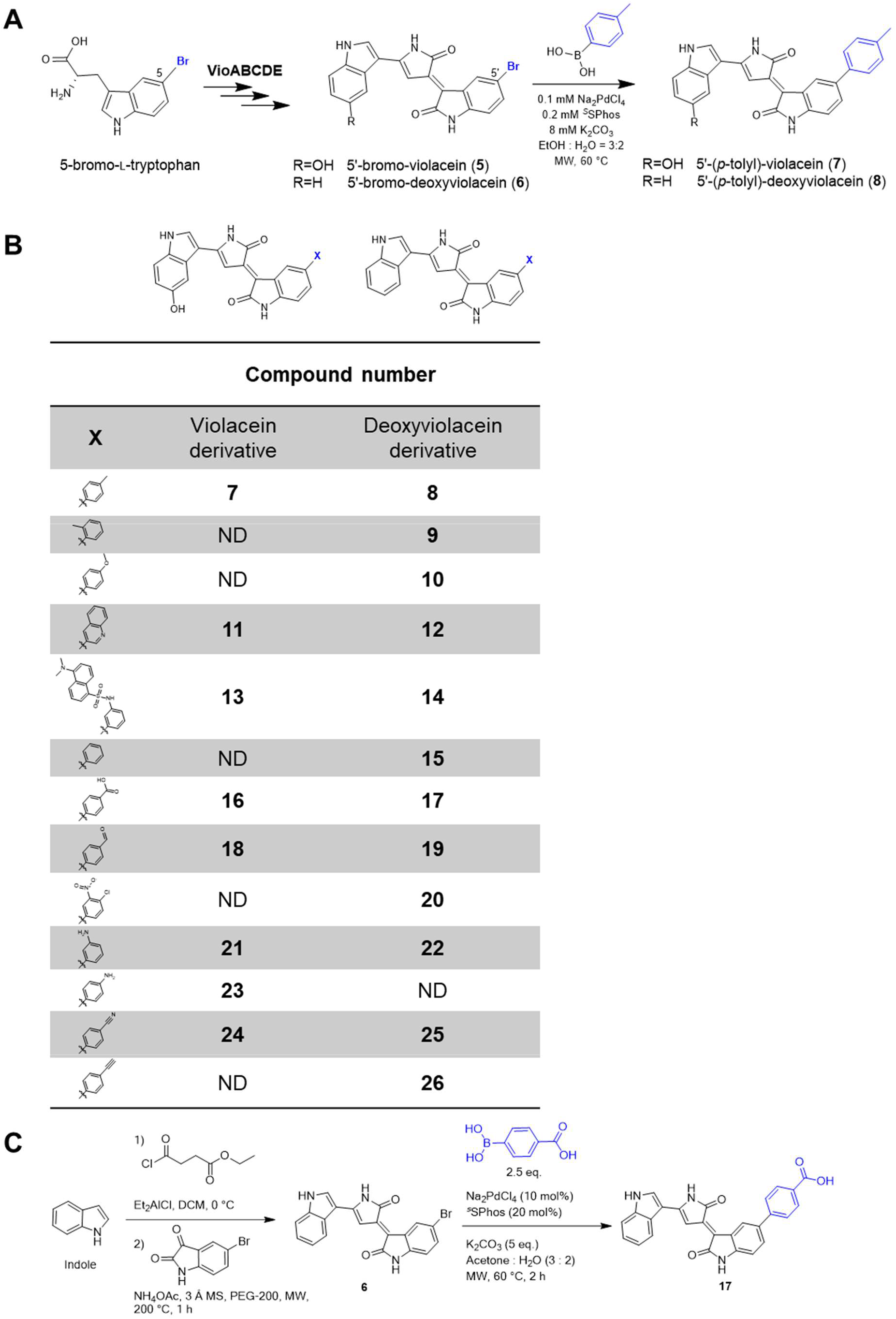
Further derivatisation of violacein and deoxyviolacein analogues via Suzuki-Miyaura cross-coupling reactions directly in crude extracts. See also Figures S3, S4 and Table S2. **(A)** Optimised conditions for cross-coupling reaction directly in crude extracts. Coupling partner indicated is *p*-tolyl-boronic acid. **(B)** Compound numbers of full range of cross-coupled products as detected by LC-HRMS or LC-HRMS/MS (see **Table S2**). ND, not detected. **(C)** Starting from indole, 5’-bromo-deoxyviolacein (**6**) was synthesised in two steps, and used for preparative scale cross-coupling to afford the product (**17**) which was isolated and characterised by ^1^H NMR analysis.

The attempts to isolate brominated violaceins, from the cell culture extracts, proved challenging due to low titres of desired compounds and co-elution as mixtures during chromatographic purifications. Hence, we decided to synthesise 5’-bromo-deoxyviolacein (**6**), which will be useful as a synthetic standard and for preparative scale cross-coupling reaction. To this end, we adapted a two-step protocol reported for synthesis of violacein analogues (McLaughlin et al., 2014). The condensation of the 3-acylated intermediate (characterised with ^1^H and ^13^C NMR in **Figure S5**) with 5-bromo-isatin proved challenging due to formation of a mixture of products. However, the desired product **6** (characterised with ^1^H NMR in **Figure S6**) was obtained in a 15% yield after two difficult chromatographic purification steps, further highlighting the benefits of the engineered biosynthesis. The LC retention time and LC-HRMS/MS analysis of the synthetic product matched with the biosynthetic **6**, and confirmed the identity as depicted (**Figure S3**). Furthermore, cross-coupling of synthetic **6** with *p*-carboxyphenylboronic acid proceeded with >70% yield based on LC-HRMS analysis (**Figure S4**); the product **17** was isolated (41% yield) from the crude reaction by extractive work-up (**Figure 4C**). The product was characterised by LC-HRMS/MS (**Figure S4**) and ^1^H and COSY NMR (**Figure S7**) analyses. These cross-coupling conditions provide a route to generating a diverse library of violacein analogues.

## Limitations of the study

Combining the strengths of biosynthesis and organic synthesis, our GenoChemetic approach in generating diverse compounds has proven fruitful in exploring chemical space of natural products and their derivatives. In this study, inspection of the biosynthetic assembly of violacein and crystal structure analyses of the first enzyme l-tryptophan oxidase VioA suggested that various tryptophan substrate analogues could be enzymatically processed. We utilized this finding and designed experiments combining biosynthesis and chemical synthesis to enable rapid access to 6 new halogenated violacein and deoxyviolacein analogues. Furthermore, we have expanded this collection of compounds further via Suzuki-Miyaura cross-coupling reactions exploiting a brominated analogue of violacein, accessing an additional series of 20 violacein or deoxyviolacein coupling products. However, the ability to carry out these synthetic diversifications selectively on the halometabolites as components of a crude cell extract is limited by the poor titres of desired halogenated compounds and complex mixtures which render isolation more difficult than traditional synthetic routes. Therefore, further optimisation of the biosynthesis of halometabolites to increase purity or yield of the desired compounds, or fine-tuning of cross-coupling conditions, should pave towards improved derivatisation efficiency via cross-coupling in crude extracts. Nonetheless, the combination of synthetic chemistry with synthetic biology, provides a more sustainable approach with the potential for application towards generating analogues of natural products in order to access greater chemical diversity rapidly and predictably.

## Acknowledgements

HEL was supported by an Imperial College President’s PhD Scholarship. We thank UKRI EPSRC (EP/K038648/1, EP/L011573/1 to PSF) and the European Union's Seventh Framework Programme (FP7/2007–2013/ERC grant agreement no 614779 GenoChemetics to RJMG) for funding. AMCO receives funding from EPSRC CRITICAT, EP/L016419/1. The authors thank all staff involved in Diamond Light Source Beamline I04-1 for assistance in VioA crystal data collection.

## Author Contributions

HEL, SJM, KMP, SVS, AMCO, RJMG and PSF designed the study, analysed the data, and wrote the manuscript. HEL performed experiments under guidance of SJM, KMP and PSF. AMCO performed cross-coupling reactions under guidance of SVS and RJMG. RL prepared and characterised the synthetic 5’-bromo-deoxyviolacein. AMCO and SMC performed LC-HRMS/MS analysis. RMM assisted on VioA crystallography, data collection and structure refinement. All authors read and approved the final draft of the manuscript.

## Resource Availability

### Lead Contact

Enquiries and requests for resources and reagents should be corresponded to the Lead Contact, Paul S. Freemont.

### Materials Availability

Plasmids and strains generated from this study will be made available by the Lead Contact upon reasonable request.

### Data and Code Availability

The authors declare that data supporting findings in this study is included in this paper and Supplemental Information. Sequences of plasmids used for expressing the violacein and the rebeccamycin/violacein hybrid pathways are available in the Supplemental Information. Raw data for NMR and LC-HRMS/MS analyses are available upon request. This study did not generate code.

## Declaration of Interests

The authors declare no competing interests.

## Supplemental Information

Supplemental Information is supplied as a separate PDF and an Excel file (Table S2 – Violacein and deoxyviolacein analogues MSMS fragment analysis. Related to Figures 3 and 4).

## Methods / Experimental Procedures

### Materials

Restriction enzymes (NEB), T4 DNA ligase (Promega), Phusion (Agilent) and Q5 DNA polymerase (NEB) were used for routine cloning. QIAprep Miniprep Kit (Qiagen) and Zymoclean Gel DNA Recovery Kit (Zymogen) were used for plasmid extraction and purification. All chemicals, including tryptophan analogues, were purchased from either Sigma-Aldrich, Fluorochem or Acros Organics. All tryptophan analogues used are dl-racemic mixtures except l-tryptophan and 5-hydroxy-l-tryptophan. Primers were synthesized by Integrated DNA Technologies and sequencing was performed by Eurofins Genomics. Chemically competent DH10B (NEB) or JM109 (Promega) *Escherichia coli* strains were used for plasmid maintenance, cloning and expression of pathway enzymes. Depending on the plasmid antibiotic resistance marker, LB broth supplemented by carbenicillin (50 *μ*g/mL), chloramphenicol (35 *μ*g/mL) and kanamycin (50 *μ*g/mL) was used unless stated otherwise.

### VioA purification and crystallisation

The *vioA* gene was amplified and cloned between NdeI and BamHI sites of pET-15b expression vector (Novagen) which contained an N-terminal hexahistidine tag. The vector was transformed into BL21 DE3 pLysS *Escherichia coli* (NEB) strain, grown in 1 L LB broth supplemented with 50 *μ*g/mL ampicillin at 37 °C until OD_600_ reached 0.6, and then induced with 0.2 mM IPTG and further incubation at 20 °C for 18 hours. Cells were then pelleted by centrifugation (4k rpm, 4 °C, 20 min) and resuspended in Binding Buffer (20 mM Tris pH 8, 500 mM NaCl, 10 mM Imidazole). Cells were sonicated (2 s on 2 s off at 70% amplitude for 4 min) and cell debris pelleted by centrifugation (15k rpm, 4 °C, 30 min). The supernatant was subjected to affinity purification by injecting into HisTrap HP column (GE Healthcare), with Buffer A as Binding Buffer and Buffer B as Elution Buffer (20 mM Tris pH 8, 500 mM NaCl, 500 mM Imidazole). Elution fractions were checked with SDS-PAGE, and fractions containing most VioA (49 kDa) were pooled and concentrated with Amicon Ultra-15 10K MWCO concentrator (EMD Millipore) and desalted with PD-10 Desalting Column (GE Healthcare) into GF Buffer (20 mM CHES pH 9.0, 25 mM NaCl). Protein solution was then subjected to SEC by injecting into HiLoad 16/60 Superdex 200 column (GE Healthcare) equilibrated with GF Buffer. Peak fractions were pooled and concentrated to 5 mg mL^−1^ measured on spectrophotometer with molar extinction coefficient from ProtParam (ExPASy). Concentrated proteins were then screened initially with sparse matrix (Crystal Screen 1 & 2 from Hampton Research), and hit condition was optimized in 24-well sitting drop plates with protein-to-buffer ratio 2:1. The best crystals were obtained with 100 mM HEPES pH 7.5, 9% PEG 8000 and 9% ethylene glycol incubated at 20 °C.

### Horseradish peroxidase-coupled VioA kinetic assay

The Infrared Fluorimetric Hydrogen Peroxide Assay Kit (Sigma-Aldrich #MAK166) was used for VioA kinetic assays. In each reaction well in 96-well plate, 50 *μ*L of 2X master mix (infrared peroxidase substrate, final concentration 0.4 U/mL horseradish peroxidase in 100 mM CHES pH 9 buffer) was added to 25 *μ*L tryptophan (TRP, 4FT, 5MeT, 6FT and 7MeT, diluted with 100 mM CHES pH 9 buffer) such that final concentrations would range from 2.4 *μ*M to 5 mM. The reaction is then initiated by injecting 25 *μ*L of VioA (final concentration 25 nM) in 100 mM CHES pH 9 buffer into each well (total volume per reaction is 100 *μ*L) and fluorescence is monitored for 30 s at 1-second intervals using 640/680 nm as excitation/emission wavelengths, with temperature maintained at 25 °C. A hydrogen peroxide standard calibration curve was also determined by adding 50 *μ*L of 2X master mix to 50 *μ*L hydrogen peroxide of known concentrations, incubated in the dark for 1 minute, and measured at the same conditions as above. Linear regression analysis was used to determine the slopes of linear region, and the slopes were used to fit either Michaelis-Menten model (4FT) or Substrate inhibition model (TRP, 5MeT, 6FT, 7MeT) to calculate the best-fit values and 95% confidence intervals of *k_cat_*, *K_M_* and *K_i_* (where applicable) in GraphPad Prism 9.0.1.

### Determination of apo VioA and tryptophan-bound VioA structures

Due to poor crystal growth and multiple needle formation was observed when we attempted to co-crystallise VioA with L-tryptophan, VioA crystal was first grown without tryptophan and then subsequently soaked in tryptophan solution. Briefly, VioA crystal was fished and cryoprotected with 20% ethylene glycol, and then snap frozen in liquid nitrogen before sending to Diamond Light Source for X-ray diffraction. Tryptophan-soaked crystals (l-tryptophan, 4-fluoro-dl-tryptophan, 5-methyl-dl-tryptophan, 6-fluoro-dl-tryptophan and 7-methyl-dl-tryptophan) were prepared by immersing crystal in mother liquor with added respective 1 mM l-tryptophan or 2 mM racemic tryptophan analogues for 30 minutes at room temperature. Datasets collected from DLS (i04-1 or i03 beamlines) were processed with in-house DIALS/Xia2 algorithm, and phase information was solved by PHASER (McCoy et al., 2007) molecular replacement with a model structure with PDB ID **<u>3X0V</u>** (l-lysine oxidase, 17% sequence identity). Further rounds of refinement and model building was done in Phenix (Adams et al., 2010) and *Coot* (Emsley et al., 2010) respectively (see **Table S1** for data collection and refinement statistics). l-tryptophan and its analogues-bound VioA structures were solved using molecular replacement with the apoenzyme structure as model. Protein structures are visualized using PyMOL Molecular Graphics System, Version 1.8 Schrödinger, LLC.

### Constructing Rebeccamycin/Violacein hybrid pathway

The sequences of Rebeccamycin and Violacein pathway genes are available from the UniProt database. Violacein genes were PCR-amplified from <u>Bba_K274002</u> plasmid (iGEM Parts Registry), adding NdeI and BamHI sites on the 5’ and 3’ ends respectively. RebF (<u>Q8KI76</u>), RebH (<u>Q8KHZ8</u>), RebO (<u>Q8KHS0</u>) and RebD (<u>Q8KHV6</u>) were synthesised commercially (GeneArt, Thermo Fisher) and cloned into appropriate pBP-ORF vector between NdeI and BamHI sites. Assembly variants of either the violacein pathway (pTU2S(VioAE)-b-VioBCD) or the violacein/rebeccamycin hybrid pathway (pTU2-A-RebOD-VioCDE, pTU2S(VioABEDC)-a-RebFH and pTU3-A-RebFHOD-VioCDE-KanR) were constructed using a modular cloning toolkit EcoFlex^22^, and most of the genetic parts are available on Addgene (<u>Kit #1000000080</u>). Briefly, all genes, including a kanamycin resistance marker KanR, were cloned into holding vector pBP-ORF, and then assembled into individual transcription units with defined promoters (BBa_J23114, BBa_J23108, SJM932, SJM933, SJM935, T7-RBS-His), pET-RBS and terminators (BBa_B0015, L2U2H09, L2U5H08 and L3S2P21). For full assembly protocol please refer to **SI Protocol**. Plasmid and protein sequences are available in **SI Appendices**.

### Culture growth and violacein extraction protocol

For test cultures to check for presence of novel analogues, typically 10 mL of autoclaved LB media in 50 mL sterile tubes were used. *E. coli* JM109 strain (Promega) harbouring the violacein or hybrid pathways was inoculated into overnight culture, which was then diluted 1:100 with fresh media and grown for 4 to 6 hours at 37 °C until reaching OD 0.5. Cultures were then incubated at 25 °C for up to 65 hours. Cells were harvested by centrifugation (4000 x *g*, 4 °C, 10 min) and supernatant was removed. For each 10 mL of culture grown, 1 mL of ethanol was added to the cell pellet, and the mixture was transferred into smaller tubes and incubated at 75 °C for about 30 min, or until cell pellet turns greyish-white. The mixture was then centrifuged (15,000 x *g*, RT, 1 min) and the supernatant transferred to fresh tube for storage at −20 °C or diluted 1:50 with ethanol for injection into LC-HRMS/MS for detection of analogues.

### LC-HRMS/MS analyses of violacein analogues

Violacein analogues were detected by Agilent 1290 LC and 6550 Q-ToF mass spectrometer with electrospray ionization, using Agilent Zorbax Extend C-18 column (2.1 x 50 mm, 1.8 *μ*m particle size). For LC, buffer A was water with 0.1% (v/v) formic acid and buffer B was acetonitrile with 0.1% (v/v) formic acid, for total running time of 13 minutes including 3 min post-run time. 2 *μ*L diluted samples were injected, and the column was washed at 0.35 ml/min with the following gradient: 0 min 5% B, 1 min 5% B, 9 min 70% B, 10 min 70% B. Quantification was by peak area of integrated chromatograms at targeted [M+H]^+^ mass with ±20 ppm window. The protonated molecules of each analyte, [M+H]^+^, were targeted and subjected to collision-induced dissociation (collision energy 30 eV), with product ions accumulated throughout the analysis. In calculating percentage composition of analogues, all compounds were assumed to have the same mass response as there is no mass spectrometry standard available for every compound detected.

### Generation and extraction of brominated violacein analogues

*E. coli* JM109 cells (harbouring the pTU2S(VioAE)-b-VioBCD plasmid) were cultured in TB media (1 L in 2 L flask) at 37 °C for 9 hours until OD600 reaches 0.4-0.6. Commercial 5-bromo-dl-tryptophan (Sigma) was added (1 mM final concentration) and cultures were further incubated for at 25 °C for 2 days. Cells were harvested by centrifugation (4000 rpm, 20 min) and stored at −20 °C. For extraction, 30 mL ethanol was added to 4 mL of *E. coli* frozen pellets containing brominated violacein, and the solution was stirred (500 rpm) at 50 °C for 1 hour. After cooling to room temperature, the extract was centrifuged (3000 rpm, 10 mins) and the supernatant crude extract containing violacein species were aliquoted (1 mL) and stored at −20 °C.

### Cross-coupling of brominated violacein from crude extract and LC-HRMS/MS analysis

In a glass microwave vial, Na_2_PdCl_4_ (100 *μ*L, 1mM stock in water) and ^*S*^SPhos (100 *μ*L, 2 mM stock in water) were mixed for 5 minutes in ethanol (300 *μ*L). Then, freshly prepared solution of appropriate boronic acid (200 *μ*L, 20 mM stock in ethanol), K_2_CO_3_ (200 *μ*L, 40 mM stock in water) and crude violacein extract (100 *μ*L in ethanol) were sequentially added, so the final concentration of ethanol was 60%. The vial was sealed and heated in a microwave reactor at 60 °C for 2 hours (microwave reactions were carried out using Biotage Initiator+ in appropriate Biotage Microwave Vials sealed with aluminium crimp caps). The palladium was then quenched with a solution of DTT (1:1 volume, 200 mM in water) and left stirring for 30 minutes. The solution was finally diluted by 5-fold (1:1 water:ethanol), centrifuged (13000 rpm, 10 min) and clear supernatant was analysed by LC-HRMS/MS. High resolution mass spectra were recorded on a Waters Micromass LCT time of flight mass spectrometer coupled to a Waters 2975 HPLC system or an Orbitrap VELOS pro. Data analysis was performed with Thermo Xcalibur 3.0.63.3 software. Samples were prepared in a 1:1 mixture of methanol water and injected with a 100 *μ*L syringe. The solvent system consisted of 0.1% formic acid in water (A) and acetonitrile (B). Products were eluted using a standard gradient from 5% B to 100% B over 15 minutes with a flow rate of 0.35 mL/min. The column used was a Kinetex 2.6 *μ*m EVO C18 100 Å, LC Column 100 × 2.1 mm.

### Synthesis of 5’-bromo-deoxyviolacein (6)

Step 1: Indole (586 mg, 5.0 mmol, 1.0 eq.) was dissolved in dichloromethane in a 3-necked flask and cooled to 0 °C, the flask was flushed with argon and sealed with a Suba-Seal. To this was added Et_2_AlCl (7.5 mL, 7.5 mmol, 1.5 eq.) as a 1.0 M solution in hexanes, via syringe through the seal. The resulting mixture was stirred at 0 °C for 25 mins. After this time the ethyl-4-chloro-4-oxobutyrate was added slowly (over 5 mins) via syringe through the seal, maintaining the reaction temperature between 0 and 2 °C. The reaction mixture was stirred for a further 2 hours allowing warming to room temperature. The resulting dark red solution was quenched by slowly adding NH_4_Cl (Sat. aq.) (5 mL). Water was added to the resulting suspension and this was then extracted with ethyl acetate, washed with Rochelle’s salt twice, washed NaHCO_3_ (aq.) twice and dried over magnesium sulfate, then filtered and evaporated *in vacuo* to give a pale-yellow solid. The solid was further purified by flash chromatography eluting gradient 20-70% ethyl acetate in hexane to give yellow solid 0.58 g, 47% yield. ^**1**^**H NMR (500 MHz, CDCl_3_)** δ 8.60 (s, 1H), 8.42 – 8.34 (m, 1H), 7.92 (d, *J* = 3.0 Hz, 1H), 7.46 – 7.39 (m, 1H), 7.32 – 7.28 (m, 2H), 4.17 (q, *J* = 7.1 Hz, 2H), 3.24 (t, *J* = 6.9 Hz, 2H), 2.80 (t, *J* = 6.9 Hz, 2H), 1.27 (t, *J* = 7.1 Hz, 3H). ^**13**^**C NMR (126 MHz, CDCl_3_)** δ 194.0 (C=O), 173.9 (C=O), 136.5 (C_*arom*_), 131.8 (CH_*arom*_), 125.5 (C_*arom*_), 123.7 (CH_*arom*_), 122.7 (CH_*arom*_), 122.2 (CH_*arom*_), 117.4 (C_*arom*_), 111.7 (CH_*arom*_), 60.9 (CH_2_), 34.1 (CH_2_), 28.6 (CH_2_), 14.3 (CH_3_). **HRMS (ESI+)** *m/z* calculated for C_14_H_16_NO_3_ [M+H]^+^ 246.1125, found 246.1122.

Step 2: Above intermediate (50 mg, 0.203 mmol, 1.0 eq.) was weighed into a MW vial, followed by 3Å molecular sieves (pre-dried, 260 mg), 5-bromo-isatin (53 mg, 0.23 mmol, 1.15 eq.), NH_4_OCOMe (79 mg, 0.815 mmol, 4.0 eq.) and PEG-200 (1.5 mL, pre-dried over 3Å molecular sieves for 24 hours). The vial was sealed and then evacuated and filled with argon 3 times and stirred at room temperature for 30 mins. The sealed vial was then heated in the microwave at 200 °C for 3 hours. The resulting dark blue solution was diluted with water and ethyl acetate forming an emulsion, brine was added to break the emulsion and the mixture was extracted with ethyl acetate, organic layers were combined and washed with brine and then dried over magnesium sulfate. The resulting purple solid was further purified by flash chromatography 10-100% ethyl acetate in hexane. The bromo-violacein enriched material was further purified by HPLC reversed phase C_18_ column (Kinetex^®^ 5 μm EVO C18 100 Å, 250×10.0 mm) on an HPLC, eluting with water (containing 0.1% formic acid) and acetonitrile. Starting conditions at 25% acetonitrile, with a one-hour gradient to 50% acetonitrile. Purple fractions were collected and combined to yield 5’-bromo-deoxyviolacein **6**(12 mg, 15% yield). ^**1**^**H NMR (500 MHz, Acetone-*d*_6_)** δ 11.39 (s, 1H), 9.89 (s, 1H), 9.82 (s, 1H), 9.31 (d, *J* = 2.0 Hz, 1H), 8.29 – 8.22 (m, 1H), 8.05 – 7.93 (m, 1H), 7.86 (d, *J* = 2.0 Hz, 1H), 7.61 (dd, *J* = 6.6, 1.8 Hz, 1H), 7.38 (dd, *J* = 8.3, 2.1 Hz, 1H), 7.36 – 7.31 (m, 1H), 6.90 (d, *J* = 8.3 Hz, 1H), 5.62 (s, 1H). **HRMS (ESI^+^)** *m/z* calculated for C_20_H_13_BrN_3_O_2_ [M+H]^+^ 406.0186, found 406.0182.

### Cross-coupling of synthetic 6 on preparative scale

In a glass microwave vial, Na_2_PdCl_4_ (100 μL from 5 mM stock solution, 10 mol%) and ^*S*^SPhos (100 μL from 10 mM stock solution, 20 mol%) were mixed for 5 minutes in water:acetone (1:1, 400 μL). Then a freshly prepared solution of *p*-carboxyphenylboronic acid (200 μL from 60 mM stock solution in ethanol, 12.3 μmol, 2.5 eq.) was added followed by K_2_CO_3_ (3.4 mg, 24.6 μmol, 5 eq.) and 5’-bromo-deoxyviolacein (**6**) (2 mg, 4.9 mmol, in 200 μL of acetone, 1 eq.). The vial was sealed and heated at 60 °C for 2 hours in a microwave reactor. The yield was estimated to be >70% from LC-MS analysis. The solvent was removed under vacuum. The solid was washed with ethanol, before being partially solubilised in acetone. The acetone was transferred to a clean vial and dried under vacuum, to afford 5’-(*p*-carboxyphenyl)-deoxyviolacein **17** (0.9 mg, 41% yield). ^**1**^**H NMR (500 MHz, Acetone-*d*_6_)** δ 9.32 (d, *J* = 2.0 Hz, 1H), 8.28 (s, 1H), 8.17 (d, *J* = 8.5 Hz, 1H), 8.04 (m, 1H), 8.01 (d, *J* = 7.6 Hz, 1H), 7.91 (d, *J* = 8.8 Hz, 1H), 7.90 (s, 1H), 7.62 (m, 2H), 7.52 (t, *J* = 7.7 Hz, 1H), 7.40 (d, *J* = 2.1 Hz, 1H), 7.38 (d, *J* = 2.0 Hz, 1H), 7.35 (m, 2H), 6.92 (d, *J* = 8.8 Hz, 1H), 6.89 (d, *J* = 8.2 Hz, 1H). **HRMS (ESI^+^)** *m/z* calculated for C_27_H_18_N_3_O_4_ [M+H]^+^ 448.1292, found 448.1286.

## Notes

### Competing Interest Statement

The authors have declared no competing interest.

### Summary of Updates

Updated compound number and figures; updated VioA kinetics parameter values with CI range.

