## Supplementary Information for "A GenoChemetic strategy for derivatization of the violacein natural product scaffold"

### SI Figures
#### Figure S1

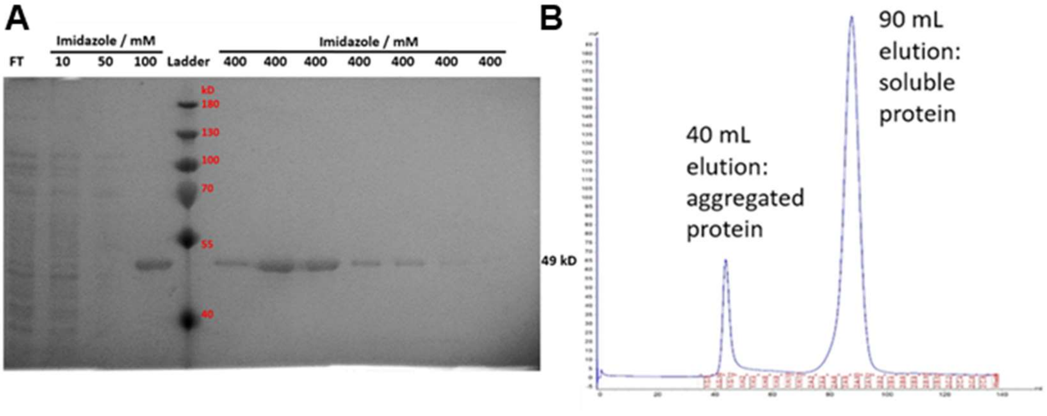

**Figure S1. His-tagged VioA (49 kD) was purified to homogeneity and subjected to horseradish peroxidase-coupled kinetics assay. Related to Figure 2. (A)** SDS-PAGE gel showing purification of His-tagged VioA using affinity chromatography. FT, flow through. Protein ladder sizes are marked in red in kD. **(B)** Size exclusion chromatograph showing protein elution peaks on the Superdex 200 column. The first peak at about 40 mL is likely protein aggregates, while the second peak at about 90 mL is likely the soluble VioA.

#### Figure S2

**A**

**Interatomic distance with FAD N5**

| Ligand | Cα (Å) | Amine N (Å) |
|---|---|---|
| TRP | 3.93 | 5.07 |
| 4FT | 4.11 | 3.62 |
| 5MeT | 4.69 | 5.48 |
| 6FT* | 3.93, 4.07 | 4.65, 5.34 |
| 7MeT | 4.68 | 5.63 |

**B**

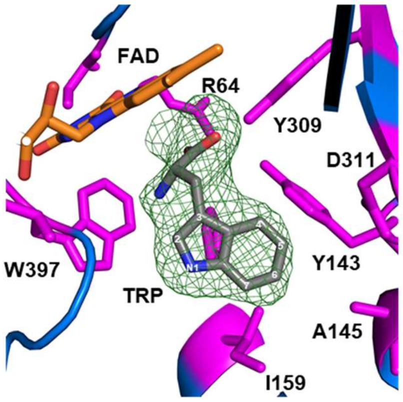

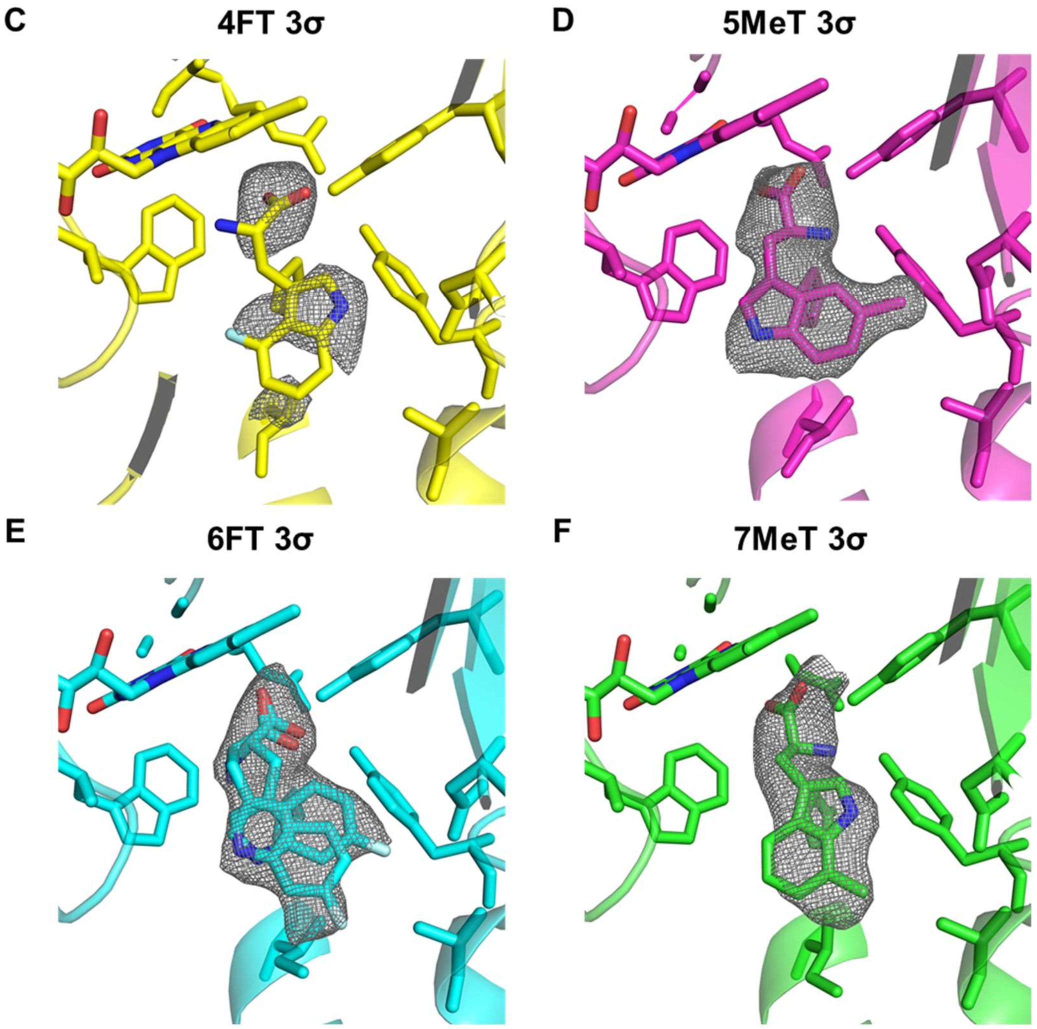

**Figure S2. VioA crystal structure in complex with L-tryptophan and tryptophan analogues. Related to Figure 2. (A)** Interatomic distance of N5 of cofactor FAD with either alpha carbon (Cα) or amine nitrogen (N) of tryptophan ligands in ligand-bound VioA active site. *Two conformers of the 6FT ligand are fitted in the omit map. **(B)** VioA active site residues (magenta) with L-tryptophan (TRP grey with carbon positions of indole ring labelled) and FAD cofactor (orange with elemental colour). In this view, H163 is located behind TRP. Green mesh is the $mF_o$-$DF_c$ omit map of TRP contoured at σ=1.5. **(C-F)** VioA active site complexed with tryptophan analogues. PDB accession codes as followed: 4FT (PDB **6FW7**), 5MeT (PDB **6FW8**), 6FT (PDB **6FW9**) and 7MeT (PDB **6FWA**). Grey mesh is the $mF_o$-$DF_c$ omit map of TRP contoured at 3σ. For 6FT structure, two conformations of the 6-fluoro-L-tryptophan are fitted into the larger-than-expected omit map.

#### Figure S3

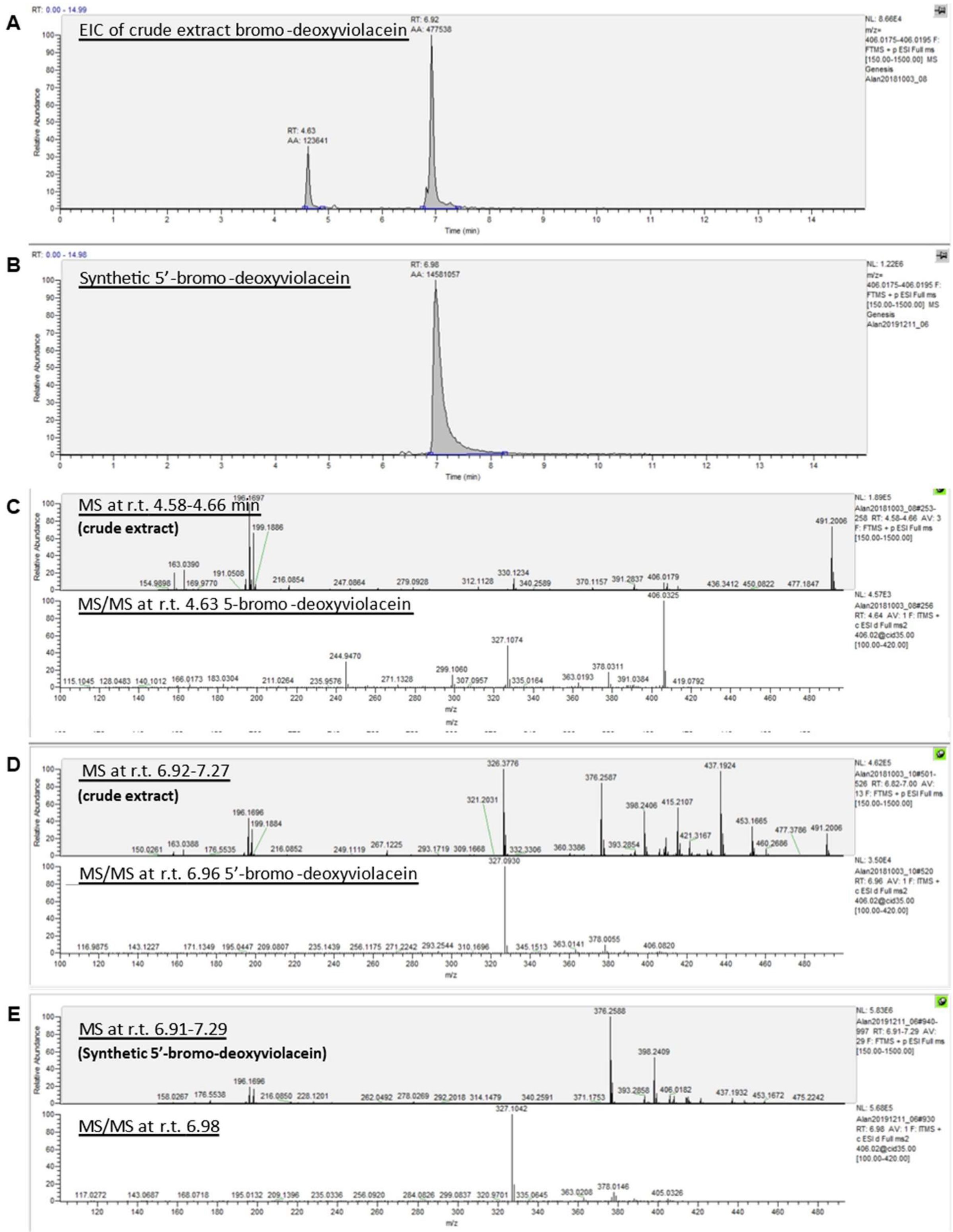

**Figure S3. LCMS analysis of the crude culture extracts containing bromo-deoxyviolaceins.** **Related to Figure 4.** **(A)** EIC for crude extract of bromo-deoxyviolacein. **(B)** EIC for synthetic 5’-bromo-deoxyviolacein. **(C)** Accurate mass spectra and HCD MS/MS fragmentation spectra for 5-bromo-deoxyviolacein $C_{20}H_{13}{}^{79}BrN_3O_2{}^+$ isotope at retention time 4.63 min from crude extract. **(D)** Accurate mass spectra and HCD MS/MS fragmentation spectra for 5'-bromo-deoxyviolacein $C_{20}H_{13}{}^{79}BrN_3O_2{}^+$ isotope at retention time 6.92 min from crude extract. **(E)** Accurate mass spectra and HCD MS/MS fragmentation spectra for synthetic 5'-bromo-deoxyviolacein $C_{20}H_{13}{}^{79}BrN_3O_2{}^+$ isotope at retention time 6.92 min.

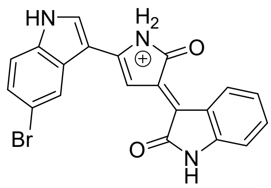

**5-bromo-deoxyviolacein**
Chemical Formula: $C_{20}H_{13}BrN_3O_2{}^+$
Exact Mass: 406.02

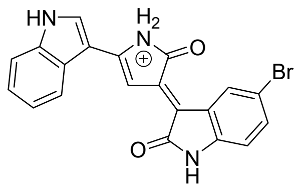

**5'-bromo-deoxyviolacein**
Chemical Formula: $C_{20}H_{13}BrN_3O_2{}^+$
Exact Mass: 406.02

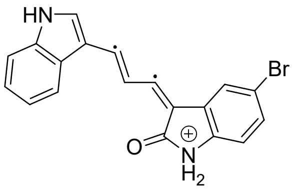

Chemical Formula: $C_{19}H_{12}BrN_2O^{2\bullet+}$
Exact Mass: 363.01

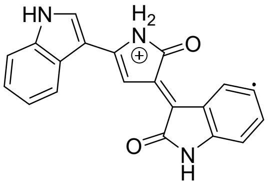

Chemical Formula: $C_{20}H_{13}N_3O_2{}^{\bullet+}$
Exact Mass: 327.10

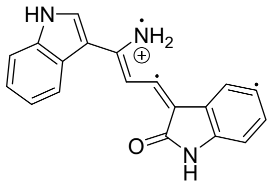

Chemical Formula: $C_{19}H_{13}N_3O^{3\bullet+}$
Exact Mass: 299.11

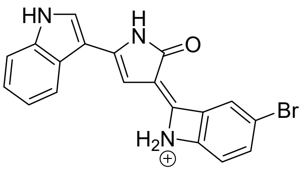

Chemical Formula: $C_{19}H_{13}BrN_3O^+$
Exact Mass: 378.02

MS/MS fragmentation pattern for 5'-bromo-deoxyviolacein. Isatin derivatives are known to undergo loss of CO and ring contraction during fragmentation.

#### Figure S4

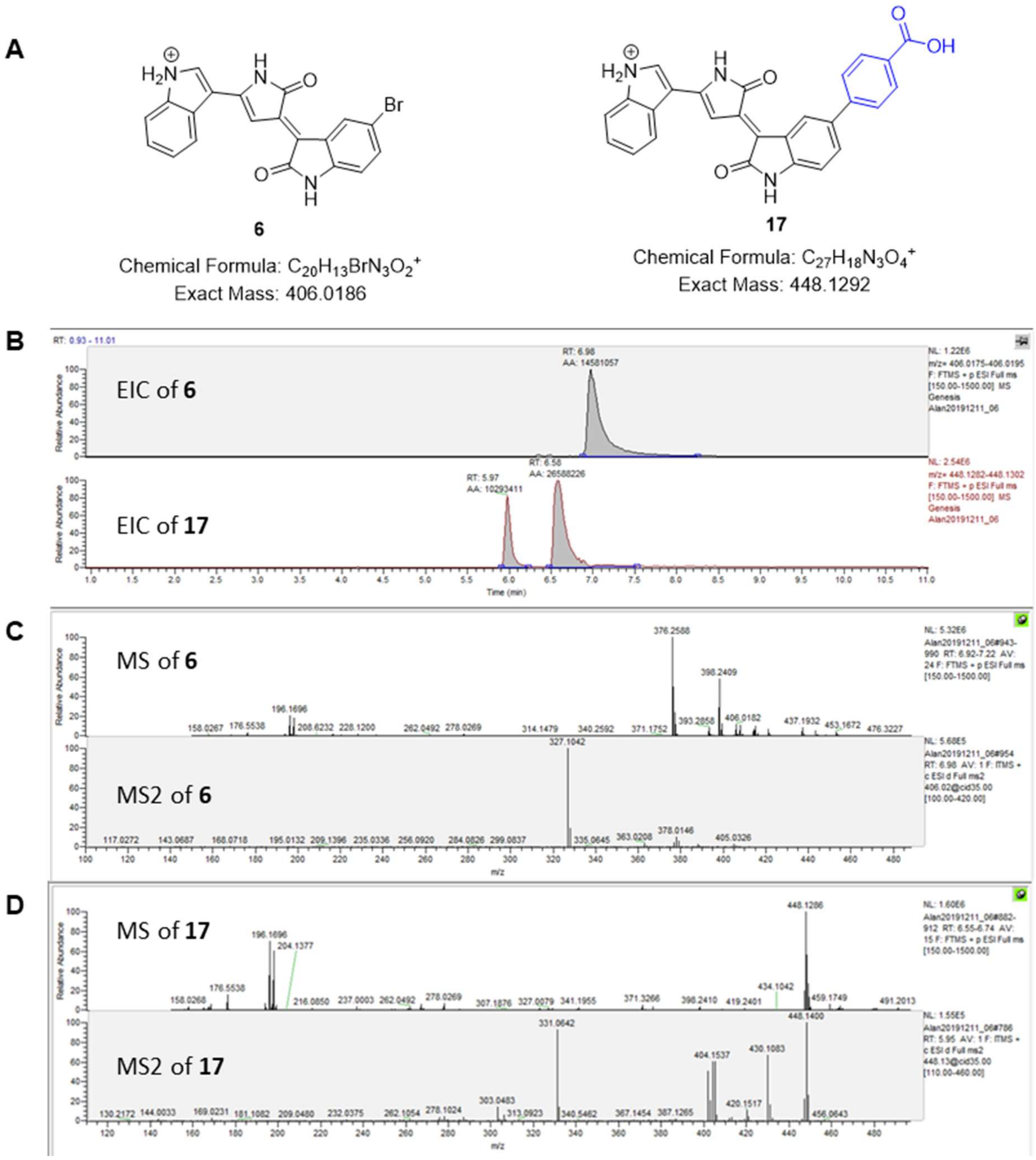

**Figure S4. LC-HRMS/MS analysis of synthetic 5'-bromo-deoxyviolacein (6) and 5'-(*p*-carboxyphenyl)-deoxyviolacein (17). Related to Figure 4. (A)** Structures of **6** and **17**. **(B)** EIC for **6** and **17**. **(C)** Accurate mass spectra and HCD MS/MS fragmentation spectra for **6** $C_{20}H_{13}{}^{79}BrN_3O_2^+$ isotope at retention time 6.92 min. **(D)** Accurate mass spectra and HCD MS/MS fragmentation spectra for **17** $C_{27}H_{18}N_3O_4^+$ isotope at retention time 6.55 min.

### Figure S5

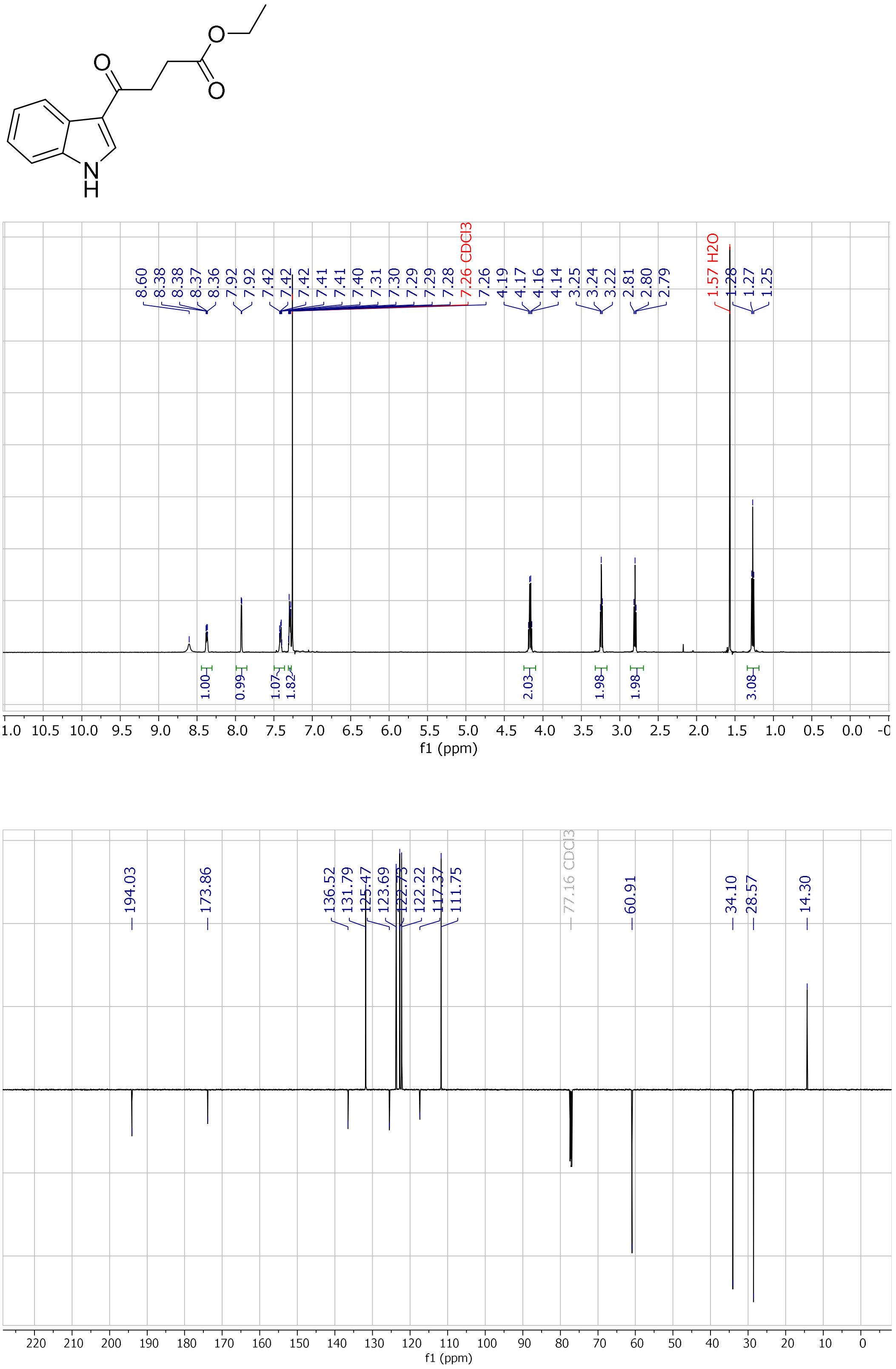

**Figure S5.** $^{1}H$-NMR ($CDCl_3$, 500 MHz) and $^{13}C$-NMR ($CDCl_3$, 126 MHz) of 3-acylated indole intermediate used for synthetic preparation of **6**, related to **Figure 4**.

### Figure S6

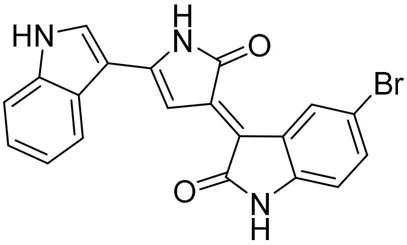

**6**

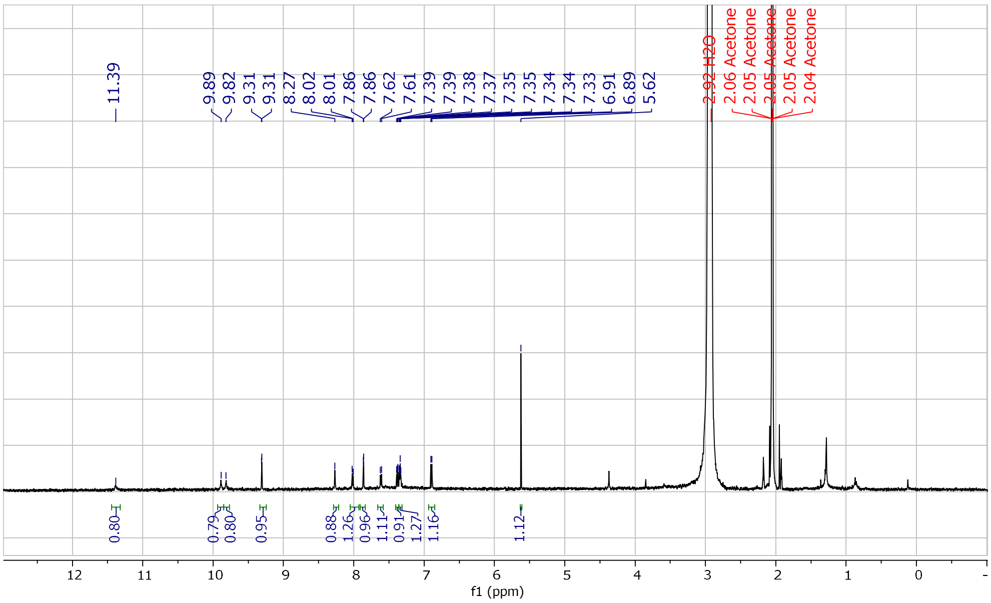

**Figure S6.** $^{1}$H-NMR (Acetone-$d_6$, 500 MHz) of synthetic 5'-bromo-deoxyviolacein **6.** Related to **Figure 4**.

### Figure S7

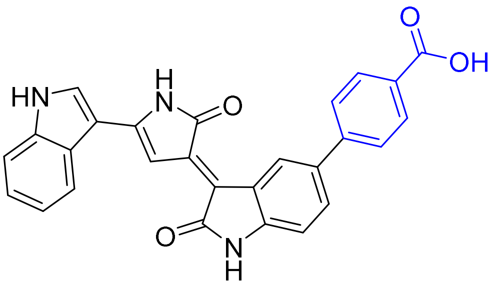

**17**

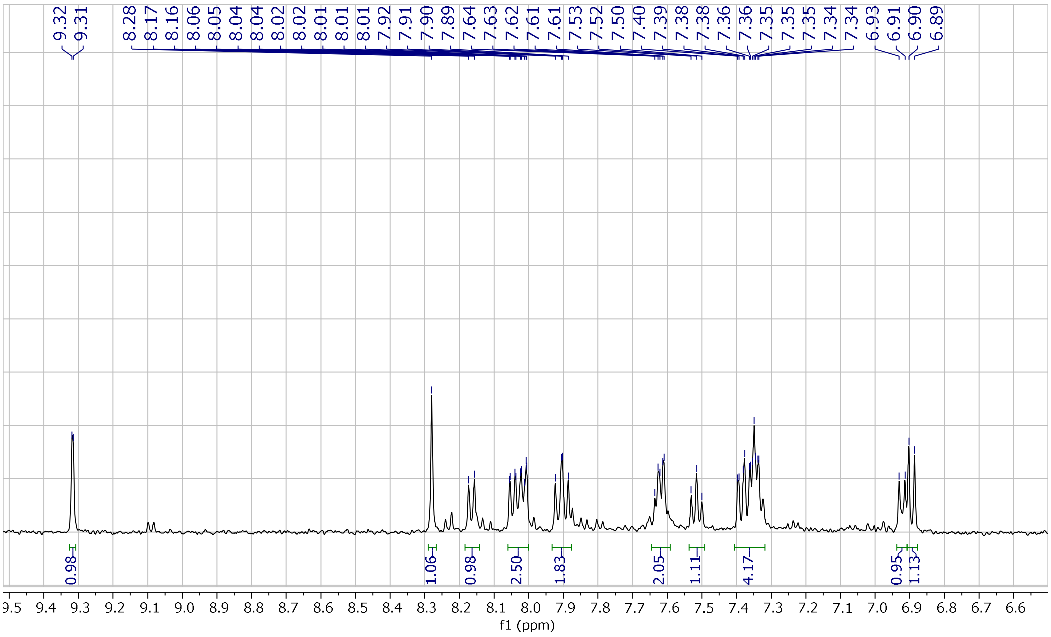

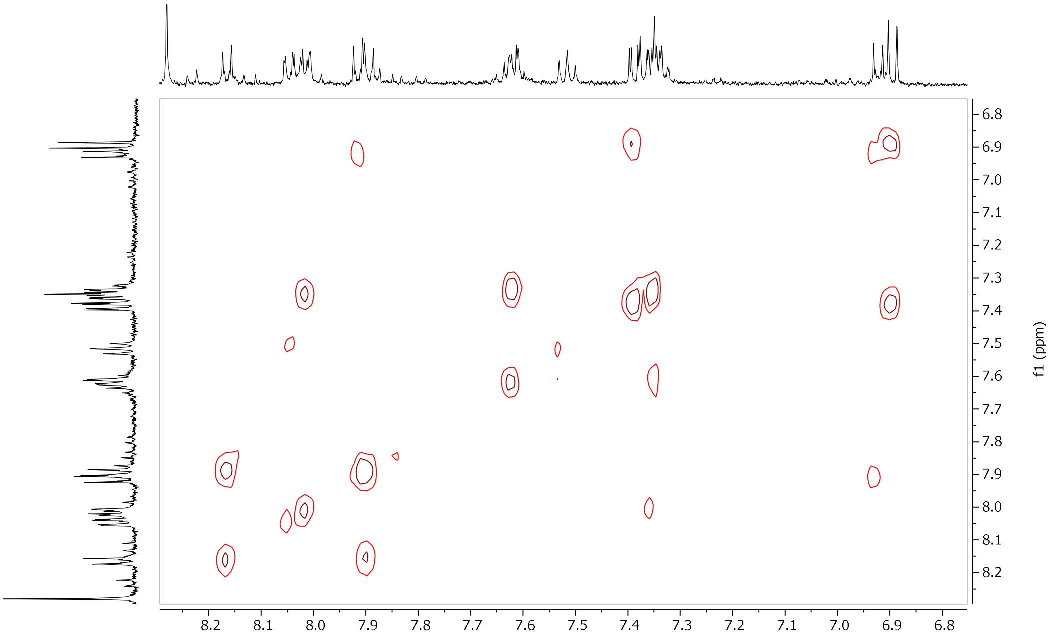

**Figure S7.** ¹H-NMR (Acetone-*d*₆, 500 MHz) of and COSY spectrum of 5'-(*p*-carboxyphenyl)-deoxyviolacein **17**. Related to **Figure 4**.

### SI Tables

#### Table S1 – VioA crystal structure data collection and refinement statistics. Related to Figure 2.

| | VioA_Trp | VioA_apo |
|---|---|---|
| PDB accession code | **6G2P** | **6ESD** |
| Wavelength (Å) | 0.9282 | 0.9282 |
| Resolution range (Å) | 76.046 - 2.6 (2.693 - 2.6) | 63.78 - 2.6 (2.693 - 2.6) |
| Space group | *C* 2 2 $2_1$ | *C* 2 2 $2_1$ |
| Unit cell dimension (*a* / *b* / *c*) (Å) | 152.09 / 175.34 / 94.44 | 151.61 / 172.6 / 94.67 |
| Unit cell angles ($\alpha$ / $\beta$ / $\gamma$) (˚) | 90 / 90 / 90 | 90 / 90 / 90 |
| Total reflections | 168072 (14462) | 173410 (14994) |
| Unique reflections | 38572 (3709) | 38134 (3731) |
| Multiplicity | 4.4 (3.9) | 4.5 (4.1) |
| Completeness (%) | 0.98 (0.95) | 0.99 (0.96) |
| Mean I/sigma(I) | 10.09 (2.43) | 24.35 (4.34) |
| Wilson B-factor | 45.96 | 48.36 |
| $R_{merge}$ | 0.04712 (0.2496) | 0.04776 (0.302) |
| $R_{meas}$ | 0.05349 (0.2891) | 0.05391 (0.3465) |
| $CC_{1/2}$ | 0.999 (0.927) | 0.999 (0.907) |
| CC* | 1 (0.981) | 1 (0.975) |
| Reflections used in refinement | 38554 (3708) | 38070 (3732) |
| Reflections used for R-free | 1851 (177) | 1958 (182) |
| $R_{work}$ | 0.1776 (0.2138) | 0.2089 (0.3065) |
| $R_{free}$ | 0.2236 (0.2728) | 0.2562 (0.4204) |
| $CC_{work}$ | 0.952 (0.913) | 0.867 (0.677) |
| $CC_{free}$ | 0.912 (0.846) | 0.849 (0.539) |
| Number of non-hydrogen atoms | 6791 | 6333 |
| Macromolecules | 6432 | 6176 |
| Ligands | 116 | 108 |
| Protein residues | 828 | 827 |
| $RMS_{bonds}$ | 0.740 | 0.758 |
| $RMS_{angles}$ | 5.16 | 5.33 |
| Ramachandran favoured (%) | 95 | 95 |
| Ramachandran allowed (%) | 4.6 | 4.9 |
| Ramachandran outliers (%) | 0 | 0.25 |
| Rotamer outliers (%) | 2 | 0.68 |
| Clashscore | 4.61 | 7.78 |
| Average B-factor | 47.66 | 58.12 |
| macromolecules | 47.89 | 58.41 |
| Ligands | 35.70 | 44.95 |
| Solvent | 47.32 | 50.43 |

Statistics for the highest-resolution shell are shown in parentheses.

| | VioA_4FT | VioA_5MeT |
|---|---|---|
| **PDB accession code** | **6FW7** | **6FW8** |
| **Wavelength (Å)** | 0.98 | 0.98 |
| **Resolution range (Å)** | 59.04 - 3.0 (3.120 - 3.000) | 58.875 - 2.400 (2.449 - 2.400) |
| **Space group** | *C* 2 2 $2_1$ | *C* 2 2 $2_1$ |
| **Unit cell dimension (*a* / *b* / *c*) (Å)** | 151.81 / 174.88 / 93.92 | 151.13 / 174.09 / 93.92 |
| **Unit cell angles (*α* / *β* / *γ*) (˚)** | 90 / 90 / 90 | 90 / 90 / 90 |
| **Total reflections** | 164133 (16850) | 318997 (28735) |
| **Unique reflections** | 25435 (2508) | 48699 (4768) |
| **Multiplicity** | 6.5 (6.7) | 6.6 (6.0) |
| **Completeness (%)** | 1.00 (1.00) | 0.98 (1.00) |
| **Mean I/sigma(I)** | 5.47 (2.47) | 9.44 (2.18) |
| **Wilson B-factor** | 61.24 | 52.43 |
| **$R_{merge}$** | 0.2695 (0.8365) | 0.1053 (0.7248) |
| **$R_{meas}$** | 0.2939 (0.9061) | 0.1147 (0.7947) |
| **$CC_{1/2}$** | 0.848 (0.61) | 0.979 (0.7) |
| **CC*** | 0.958 (0.871) | 0.995 (0.907) |
| **Reflections used in refinement** | 25342 (2480) | 47819 (4507) |
| **Reflections used for R-free** | 1229 (134) | 2327 (226) |
| **$R_{work}$** | 0.1959 (0.2807) | 0.1895 (0.3452) |
| **$R_{free}$** | 0.2592 (0.3221) | 0.2449 (0.4278) |
| **$CC_{work}$** | 0.904 (0.879) | 0.953 (0.843) |
| **$CC_{free}$** | 0.911 (0.851) | 0.933 (0.648) |
| **Number of non-hydrogen atoms** | 6507 | 6707 |
| **macromolecules** | 6359 | 6470 |
| **Ligands** | 140 | 140 |
| **Protein residues** | 827 | 834 |
| **$RMS_{bonds}$** | 0.736 | 0.731 |
| **$RMS_{angles}$** | 5.22 | 5.13 |
| **Ramachandran favoured (%)** | 91 | 95 |
| **Ramachandran allowed (%)** | 8.6 | 4.5 |
| **Ramachandran outliers (%)** | 0.72 | 0 |
| **Rotamer outliers (%)** | 1.4 | 1.2 |
| **Clashscore** | 13.46 | 8.92 |
| **Average B-factor** | 65.69 | 66.14 |
| **macromolecules** | 65.98 | 66.55 |
| **Ligands** | 53.42 | 54.34 |
| **Solvent** | 46.32 | 55.88 |

Statistics for the highest-resolution shell are shown in parentheses.

| | VioA_6FT | VioA_7MeT |
|---|---|---|
| **PDB accession code** | **6FW9** | **6FWA** |
| **Wavelength (Å)** | 0.98 | 0.98 |
| **Resolution range (Å)** | 63.86 - 2.739 (2.828 - 2.739) | 47.134 - 2.846 (2.947 - 2.846) |
| **Space group** | *C* 2 2 $2_1$ | *C* 2 2 $2_1$ |
| **Unit cell dimension (*a* / *b* / *c*) (Å)** | 151.48 / 174.32 / 93.82 | 151.34 / 172.42 / 94.27 |
| **Unit cell angles (*α* / *β* / *γ*) (˚)** | 90 / 90 / 90 | 90 / 90 / 90 |
| **Total reflections** | 212945 (20401) | 188810 (18086) |
| **Unique reflections** | 33026 (3237) | 29251 (2837) |
| **Multiplicity** | 6.4 (6.3) | 6.5 (6.4) |
| **Completeness (%)** | 0.99 (1.00) | 0.95 (0.98) |
| **Mean I/sigma(I)** | 8.05 (1.91) | 5.90 (1.62) |
| **Wilson B-factor** | 57.76 | 66.48 |
| **$R_{merge}$** | 0.1456 (0.7696) | 0.1922 (0.8586) |
| **$R_{meas}$** | 0.1587 (0.8394) | 0.2095 (0.9355) |
| **$CC_{1/2}$** | 0.97 (0.615) | 0.975 (0.548) |
| **CC*** | 0.992 (0.873) | 0.994 (0.842) |
| **Reflections used in refinement** | 32621 (3094) | 27963 (2675) |
| **Reflections used for R-free** | 1595 (146) | 1445 (158) |
| **$R_{work}$** | 0.1983 (0.3413) | 0.2343 (0.3782) |
| **$R_{free}$** | 0.2617 (0.4097) | 0.3121 (0.4342) |
| **$CC_{work}$** | 0.944 (0.804) | 0.936 (0.792) |
| **$CC_{free}$** | 0.944 (0.682) | 0.883 (0.488) |
| **Number of non-hydrogen atoms** | 6686 | 6529 |
| **macromolecules** | 6412 | 6296 |
| **Ligands** | 156 | 140 |
| **Protein residues** | 832 | 832 |
| **$RMS_{bonds}$** | 0.736 | 0.738 |
| **$RMS_{angles}$** | 5.15 | 5.24 |
| **Ramachandran favoured (%)** | 91 | 90 |
| **Ramachandran allowed (%)** | 8.5 | 10 |
| **Ramachandran outliers (%)** | 0.12 | 0.12 |
| **Rotamer outliers (%)** | 1.4 | 1.6 |
| **Clashscore** | 10.04 | 28.12 |
| **Average B-factor** | 65.26 | 75.29 |
| **macromolecules** | 65.65 | 75.81 |
| **Ligands** | 53.85 | 59.60 |
| **Solvent** | 58.87 | 63.70 |

Statistics for the highest-resolution shell are shown in parentheses.

#### Table S2 – Violacein and deoxyviolacein analogues MSMS fragment analysis. Related to Figures 3 and 4.

See Table S2 excel file.

There are two excel sheets in this file:

1. "Halogenated analogues" contains violacein (**1**) and deoxyviolacein (**2**) from violacein standard (Sigma), analogues produced from feeding 5-bromo-DL-tryptophan (**5, 6**), as well as *in vivo* halogenation via RebF and RebH (**3, 4a, 4b, 4c**).
2. "Cross-coupling products" contains violacein and deoxyviolacein analogues produced from Suzuki-Miyaura cross-coupling reactions with brominated violacein or deoxyviolacein and various coupling ligands (**7** to **26**).

In some cases, structural isomers of mono-substituted analogues are distinguished by a specific fragment in MS/MS spectra. For example, 7-chloro-deoxyviolacein (chlorine attached to C7 position of indole, **4b**) was identified by fragment with *m/z* = 183, while 7'-chloro-violacein (chlorine attached to C7 position of keto-indole, **4b**) was identified by fragment with *m/z* = 217, which is the same fragment with chlorine substituting a hydrogen atom. This is due to the subtle difference in fragmentation pattern between analogues with substituted group attached to the indole or keto-indole moiety.

In addition, monochlorinated analogues demonstrate distinct M:M+2 = 3:1 isotopic pattern, whereas dichlorinated analogues demonstrate M:M+2:M+4 = 9:6:1 isotopic pattern in MS1 spectra. Monobrominated analogues demonstrate M:M+2 = 1:1 isotopic pattern. This is shown on the rightmost column with arrows pointing to the halogen isotopic peaks in MS1 ESI+ spectra.

### SI Protocol - EcoFlex assembly for Reb/Vio hybrid pathway. Related to Figures 2, 3 and 4.

Cloning of the full-length Reb/Vio hybrid pathway is carried out using EcoFlex MoClo kit (Moore et al. 2016) (available on Addgene (Kit #1000000080). This is a hierarchical assembly method utillising Type IIS restriction enzymes BsaI and BsmBI, starting from regulatory elements such as promoters and terminators (Level 0) to transcription unit (Level 1), and subsequently to sub-pathway (Level 2) and larger pathways (Level 3) with multiple transcription units. Level 0 parts are stored in pBP plasmids, which are derived from the pSB1C3 vector with pMB1 origin and chloramphenicol resistance. Level 1 plasmids are denoted as pTU1, and they are derived from pSB1A2 vector with pMB1 origin and ampicillin resistance. Level 1 plasmids also have RFP or LacZ for red/white or blue/white screening on transformation plate. Level 2 plasmids have the same pSB1C3 backbone, while Level 3 plasmids are derived from pSB1A2.

Genes were cloned from PCR product of violacein plasmid (http://parts.igem.org/Part:BBa_K274002) or synthesized with flanking NdeI and BamHI restriction sites into pBP-ORF vector. Other Level 0 parts including pBP-J23114, pBP-J23108, pET-RBS and pBP-B0015 are available from Addgene (https://www.addgene.org/cloning/moclo/freemont-ecoflex/).

To construct pTU3-A-RebFHOD-VioCDE/KanR plasmid with the full-length pathway, first assemble the Level 1 plasmids harboring the individual genes of interest with their own Level 0 parts including promoter, RBS and terminator. Assembly of Level 1 plasmid was carried out as followed:

1. Combine ~100 ng of gene of interest in pBP vector with 50 ng each of promoter, RBS, terminator and Level 1 acceptor plasmid.
2. Add 1X T4 ligase buffer, 1X BSA, 20 units BsaI and 2-3 units T4 ligase, top up to 10 or 15 $\mu$L reaction volume.
3. Incubate reaction in a thermal cycler at 37 ˚C for 5 min then 16 ˚C for 10 min, repeat cycles for 15 to 30 times, then heat inactivate the enzymes by incubating at 50 ˚C and 80 ˚C for 5 min each.
4. Transform 5 $\mu$L to 50 $\mu$L DH10β or JM109 *E. coli* cells onto LB + carbenicillin plate, selecting for white colonies.
5. Check plasmid integrity by test digestion and sequencing if necessary (see below for sequencing primers used).

The order of assembly into higher hierarchy plasmids (Level 2 and 3) are dictated by the type of pTU1 vector for each gene. For pTU2 vector, each can take 4 or 5 pTU1 vectors, while pTU3A vector can accept two pTU2 vectors. Assembly into pTU2 vector follows a similar protocol as above, except adding pTU1 plasmids instead of pBP parts, changing BsaI to BsmBI, and incubating the reaction at 37 ˚C overnight instead of thermal cycling. Assembly into pTU3 vector follows the same as Level 1 assembly protocol, except adding pTU2 plasmids instead of pTU1 plasmids.

For pTU3-A-RebFHOD-VioCDE-KanR, the parts used for assembly and the respective vectors used at Level 1, 2 and 3 are as followed.

| Gene | Promoter | RBS | Terminator | pTU1 vector | pTU2 vector | pTU3 vector |
|---|---|---|---|---|---|---|
| **RebF*** | BBa_J23114 or BBa_J23108 | pET | B0015 | pTU1-A | pTU2-A | pTU3-A |
| **RebH*** | BBa_J23114 or BBa_J23108 | pET | B0015 | pTU1-B | | |
| **RebO** | BBa_J23114 | pET | B0015 | pTU1-C | | |
| **RebD** | BBa_J23114 | pET | B0015 | pTU1-D | | |
| **VioC** | BBa_J23114 | pET | B0015 | pTU1-A | pTU2-B | |
| **VioD*** | BBa_J23114 or BBa_J23108 | pET | B0015 | pTU1-B | | |
| **VioE** | BBa_J23114 | pET | B0015 | pTU1-C | | |
| **KanR**** | BBa_J23114 | pET | B0015 | pTU1-D | | |

*The RebF, RebH and VioD genes are assembled with either BBa_J23114 or BBa_J23108 promoters depending on the combination required as shown in **Figure 3F**.

**For KanR, it is encoded downstream of another constitutive promoter (minimal FRT) and in the reverse strand with respect to the other genes, but in order to be compatible with the assembly protocol, the same promoter, RBS and terminator parts were assembled.

For pTU2-A-RebOD-VioCDE, the types of vectors at Level 1 and 2 used are as followed.

| Gene | Promoter | RBS | Terminator | pTU1 vector | pTU2 vector |
|---|---|---|---|---|---|
| **RebO** | BBa_J23114 | pET | B0015 | pTU1-A | pTU2-A |
| **RebD** | BBa_J23114 | pET | B0015 | pTU1-B | |
| **VioE** | BBa_J23114 | pET | B0015 | pTU1-C | |
| **VioD** | BBa_J23114 | pET | B0015 | pTU1-D1 | |
| **VioC** | BBa_J23114 | pET | B0015 | pTU1-E | |

Apart from assembling multiple Level 2 plasmids into Level 3 plasmid to construct a larger pathway, it is also possible to clone a Level 2 plasmid into a secondary cloning site upstream of the main assembly site in a pTU2S plasmid. These secondary sites can be accessed by cutting with BpiI and then ligate with BsaI-cut pTU2 plasmid. The main assembly site is then used for the other genes required for the full pathway.

For pTU2S-a(VioABEDC)-RebFH, the types of vectors at Level 1 and 2 used are as followed. Genes in bracket are cloned into the secondary site. T7-RBS-His promoter/RBS was used for VioABEDC to attempt minimizing conversion of tryptophan by VioA before chlorination by RebH.

| Gene | Promoter | RBS | Terminator | pTU1 vector | pTU2 vector |
|---|---|---|---|---|---|
| **VioA** | T7-RBS-His | | B0015 | pTU1-A | pTU2-A (then clone into pTU2S-a secondary site) |
| **VioB** | T7-RBS-His | | B0015 | pTU1-B | |
| **VioE** | T7-RBS-His | | B0015 | pTU1-C | |
| **VioD** | T7-RBS-His | | B0015 | pTU1-D1 | |
| **VioC** | T7-RBS-His | | B0015 | pTU1-E | |
| **RebF** | BBa_J23114 | pET | B0015 | pTU1-A | pTU2S-a (with VioABEDC in secondary site) |
| **RebH** | BBa_J23114 | pET | B0015 | pTU1-B | |

For pTU2S-b(VioAE)-VioBCD, the types of vectors at Level 1 and 2 used are as followed. Genes in bracket are cloned into the secondary site. Different promoters and terminators were used for VioBCD to minimize recombination events.

| Gene | Promoter | RBS | Terminator | pTU1 vector | pTU2 vector |
|---|---|---|---|---|---|
| **VioA** | BBa_J23114 | pET | B0015 | pTU1-A | pTU2-a (then clone into pTU2S-b secondary site) |
| **VioE** | BBa_J23114 | pET | B0015 | pTU1-B | |
| **VioB** | SJM933 | pET | L3S2P21 | pTU1-A | pTU2S-b (with VioAE in secondary site) |
| **VioC** | SJM935 | pET | L2U2H09 | pTU1-B | |
| **VioD** | SJM932 | pET | L2U5H08 | pTU1-C | |

**Part sequence**

Highlighted 4bp indicates linker that directs order of assembly in a typical Golden Gate reaction. ATG start codon is bold and underlined.

| Part Name | Sequence |
|---|---|
| **Weak promoter BBa_J23114** | CTATTTTATGGCTAGCTCAGTCCTAGGTACAATGCTAGCGTAC |
| **Medium promoter BBa_J23108** | CTATCTGACAGCTAGCTCAGTCCTAGGTATAATGCTAGCGTAC |
| **Synthetic Promoter SJM932** | CTATTTTACAGCCCGAAGAGTCCTAGGTATTGTACGGACGTAC |
| **Synthetic Promoter SJM933** | CTATTTTACGGCTCAAGCAGTCCTAGGTATAGTAGCAACGTAC |
| **Synthetic Promoter SJM935** | CTATTTGACAGCCGGAAAAGTCCTAGGTATTGTCTAGACGTAC |
| **T7-RBS-His (with in-frame His$_6$-tag)** | CTATCCCGCGAAATTAATACGACTCACTATAGGGGAATTGTGAGCGGATAACAATTCCCCTCTAGAAATAATTTTGTTTAACTTTAAGAAGGAGATATACC**A****TG**GGCAGCAGCCATCATCATCATCATCACAGCAGCGGCCTGGTGCCGCGCGGCAGCCATA |
| **pET-RBS** | GTACTTTAACTTTAAGAAGGAGATATACATA |

| | |
|---|---|
| **Synthetic terminator L2U2H09** | TCGAACGGCCCTCGCAAGGGCCGTTTTTTTGTATGTT |
| **Synthetic terminator L2U5H08** | TCGATGCCGGGAGAACATCCCGGCATTGTTGTATTGTT |
| **Synthetic terminator L3S2P21** | TCGACTCGGTACCAAATTCCAGAAAAGAGGCCTCCCGAAAGGGGGGCCT<br>TTTTTCGTTTTGGTCCTGTT |
| **Terminator BBa_B0015** | TCGACCAGGCATCAAATAAAACGAAAGGCTCAGTCGAAAGACTGGGCCTT<br>TCGTTTTATCTGTTGTTTGTCGGTGAACGCTCTCTACTAGAGTCACACTGG<br>CTCACCTTCGGGTGGGCCTTTCTGCGTTTATATGTT |
| **vioA** | CAT**ATG**AAACATTCTTCCGATATCTGCATTGTTGGTGCTGGTATTTCTGGT<br>TTGACGTGCGCAAGCCATCTGCTGGACAGCCCGGCATGCCGTGGTCTGA<br>GCCTGCGTATCTTTGACATGCAGCAAGAAGCCGGTGGCCGTATCCGCAG<br>CAAAATGCTGGATGGTAAGGCAAGCATTGAACTGGGCGCAGGTCGCTAC<br>TCCCCTCAGTTGCACCCGCATTTCCAAAGCGCAATGCAGCACTATAGCCA<br>AAAGAGCGAAGTCTATCCGTTCACCCAGTTGAAGTTCAAATCTCACGTGC<br>AGCAAAAGCTGAAGCGCGCCATGAATGAACTGTCCCCGCGTCTGAAAGA<br>GCATGGTAAAGAGAGCTTTTTGCAGTTTGTCAGCCGTTATCAAGGTCACG<br>ATAGCGCGGTTGGTATGATCCGCTCTATGGGTTACGACGCACTGTTCCTG<br>CCGGATATCAGCGCAGAAATGGCCTACGACATTGTGGGTAAGCACCCGG<br>AGATCCAGAGCGTGACGGACAACGACGCGAACCAATGGTTTGCAGCGGA<br>AACGGGCTTTGCTGGTCTGATTCAGGGCATCAAGGCTAAGGTTAAGGCG<br>GCAGGTGCGCGTTTTAGCCTGGGTTATCGTCTGCTGAGCGTCCGTACCG<br>ACGGTGACGGCTACCTGCTGCAACTGGCAGGTGACGACGGCTGGAAACT<br>GGAGCACCGTACCCGCCATCTGATTCTGGCGATTCCGCCGAGCGCGATG<br>GCGGGTTTGAATGTTGATTTTCCAGAAGCCTGGTCCGGTGCGCGCTATGG<br>CAGCCTGCCGCTGTTTAAGGGCTTTCTGACGTACGGTGAGCCGTGGTGG<br>TTGGACTACAAACTGGACGATCAGGTGCTGATTGTTGACAACCCGCTGCG<br>CAAAATCTATTTCAAAGGCGATAAGTACCTGTTCTTCTATACCGATAGCGA<br>GATGGCGAATTACTGGCGCGGTTGTGTCGCGGAGGGCGAGGACGGTTAC<br>CTGGAGCAAATTCGCACCCATTTGGCTAGCGCACTGGGTATCGTCCGTGA<br>ACGTATCCCGCAACCGCTGGCACACGTTCACAAGTATTGGGCGCACGGC<br>GTTGAGTTTTGCCGTGATTCTGATATTGACCACCCGAGCGCACTGTCTCA<br>TCGCGACAGCGGTATCATCGCGTGCTCCGATGCGTACACGGAGCATTGT<br>GGTTGGATGGAGGGCGGTCTGCTGAGCGCCCGTGAGGCAAGCCGTCTG<br>CTGTTGCAGCGTATCGCCGCGTGA |
| **vioB** | CAT**ATG**AGCATTCTGGATTTCCCGCGTATCCACTTCCGTGGCTGGGCCCG<br>TGTCAATGCGCCGACCGCGAACCGCGATCCGCACGGCCACATCGATATG<br>GCCAGCAATACCGTGGCGATGGCGGGTGAGCCGTTCGACCTGGCACGC<br>CATCCTACGGAGTTCCACCGTCACCTGCGCTCCCTGGGTCCGCGCTTCG<br>GCTTGGATGGTCGTGCTGACCCGGAAGGCCCGTTCAGCCTGGCCGAGG<br>GCTACAACGCTGCCGGTAACAACCACTTTTCGTGGGAGAGCGCAACCGTT<br>AGCCACGTGCAATGGGATGGCGGTGAGGCGGATCGTGGTGACGGTCTG<br>GTCGGTGCTCGTTTGGCACTGTGGGGTCACTACAATGATTATCTGCGTAC<br>CACCTTCAATCGTGCTCGTTGGGTCGACAGCGACCCGACGCGCCGTGAC<br>GCTGCACAAATCTATGCGGGCCAATTCACCATTAGCCCGGCTGGTGCCG<br>GTCCGGGTACGCCGTGGCTGTTTACGGCAGACATTGATGATAGCCATGG<br>TGCACGTTGGACGCGTGGCGGCCACATTGCAGAGCGTGGCGGCCACTTC<br>TTGGATGAAGAGTTTGGTCTGGCACGCCTGTTTCAGTTCTCTGTGCCGAA<br>AGATCACCCACATTTTCTGTTTCACCCGGGTCCGTTTGATTCCGAGGCCT<br>GGCGTCGTCTGCAATTGGCTCTGGAGGATGACGACGTTCTGGGTCTGAC<br>CGTGCAATATGCGTTGTTCAATATGAGCACCCCGCCTCAGCCGAACAGCC<br>CGGTTTTTCACGATATGGTCGGTGTTGTCGGTCTGTGGCGTCGTGGTGAA<br>CTGGCGAGCTACCCGGCTGGTCGTCTGCTGCGTCCGCGTCAACCGGGTC<br>TGGGTGACCTGACCCTGCGCGTCAACGGTGGTCGCGTTGCGCTGAATTT<br>GGCGTGTGCCATTCCGTTCAGCACTCGTGCCGCGCAGCCAAGCGCACCG<br>GACCGCCTGACCCCGGACCTGGGTGCCAAACTGCCGCTGGGCGATCTG<br>CTGCTGCGTGATGAGGACGGCGCACTGTTGGCACGTGTGCCGCAGGCTC |

| | |
|---|---|
| | TGTACCAAGACTATTGGACGAATCACGGTATTGTGGACCTGCCGCTGCTG<br>CGCGAACCGCGTGGTAGCTTGACCCTGAGCAGCGAACTGGCGGAGTGG<br>CGTGAGCAAGACTGGGTCACCCAAAGCGACGCGTCTAACCTGTACCTGG<br>AGGCACCGGATCGCCGTCACGGTCGCTTTTTCCCTGAGAGCATCGCGCT<br>GCGCAGCTACTTTCGCGGTGAAGCGCGTGCGCGTCCGGATATCCCGCAT<br>CGTATCGAGGGCATGGGCCTGGTCGGCGTCGAATCTCGTCAGGATGGCG<br>ACGCTGCGGAATGGCGTCTGACGGGTCTGCGTCCGGGTCCGGCACGCA<br>TTGTTCTGGACGATGGTGCCGAGGCGATCCCTCTGCGTGTTCTGCCTGAC<br>GATTGGGCGCTGGATGACGCGACCGTCGAAGAAGTGGATTACGCCTTTTT<br>GTACCGCCACGTTATGGCGTATTACGAGCTGGTGTATCCATTCATGAGCG<br>ACAAGGTGTTTTCCCTGGCTGATCGTTGCAAATGTGAAACGTACGCACGT<br>CTGATGTGGCAGATGTGTGATCCGCAGAACCGCAACAAGTCCTATTACAT<br>GCCGAGCACCCGCGAACTGTCGGCACCGAAAGCTCGTTTGTTCTTGAAG<br>TATCTGGCCCACGTGGAAGGCCAGGCACGCCTGCAAGCACCTCCGCCAG<br>CGGGTCCGGCACGCATTGAATCTAAAGCCCAGTTGGCGGCAGAGCTGCG<br>TAAAGCCGTCGACCTGGAGCTGTCTGTGATGCTGCAATACCTGTACGCGG<br>CGTATAGCATTCCGAACTATGCACAGGGCCAACAACGTGTTCGTGACGGT<br>GCGTGGACCGCCGAGCAGCTGCAACTGGCGTGCGGTAGCGGTGACCGT<br>CGCCGTGATGGCGGTATTCGTGCAGCACTGCTGGAAATTGCTCATGAAGA<br>AATGATTCATTACCTGGTCGTTAACAACCTGCTGATGGCCCTGGGCGAGC<br>CGTTCTACGCGGGTGTCCCGCTGATGGGCGAAGCGGCACGTCAGGCGTT<br>TGGCCTGGACACCGAGTTCGCTCTGGAACCGTTTAGCGAAAGCACGCTG<br>GCACGTTTTGTTCGTCTGGAATGGCCGCACTTTATCCCAGCACCGGGCAA<br>ATCCATCGCGGACTGCTATGCCGCCATTCGTCAGGCGTTTTTGGATCTGC<br>CGGACTTGTTTGGTGGCGAGGCAGGTAAGCGTGGCGGTGAACACCACCT<br>GTTCCTGAATGAGCTGACCAACCGTGCGCATCCGGGTTATCAACTGGAAG<br>TTTTCGATCGCGACTCGGCGCTGTTTGGTATTGCATTTGTGACCGATCAG<br>GGCGAAGGTGGCGCTCTGGACAGCCCGCACTACGAACATAGCCATTTTC<br>AACGTCTGCGTGAAATGAGCGCGCGTATCATGGCTCAAAGCGCACCGTT<br>CGAACCGGCGCTGCCGGCGTTGCGTAATCCGGTTCTGGATGAGAGCCCG<br>GGTTGCCAACGTGTCGCAGACGGTCGTGCGCGTGCGCTGATGGCATTGT<br>ACCAAGGCGTTTATGAGCTGATGTTTGCGATGATGGCGCAGCACTTCGCC<br>GTGAAACCGCTGGGTAGCTTGCGTCGCAGCCGCCTGATGAACGCAGCAA<br>TCGATCTGATGACCGGTCTGTTGCGTCCGCTGAGCTGCGCGCTGATGAA<br>CCTGCCAAGCGGCATCGCCGGTCGCACGGCCGGTCCGCCGCTGCCGGG<br>TCCGGTTGACACCCGTAGCTATGACGACTACGCGCTGGGCTGTCGCATG<br>CTGGCACGCCGTTGCGAGCGTCTGCTGGAGCAGGCGAGCATGCTGGAA<br>CCGGGTTGGCTGCCGGATGCGCAGATGGAGCTGCTGGATTTCTATCGTC<br>GCCAAATGCTGGACTTGGCGTGCGGCAAACTGAGCCGCGAGGCCTAA |
| **vioC** | CAT**<u>ATG</u>**AAACGTGCGATTATCGTTGGTGGCGGCCTGGCGGGTGGCCTGA<br>CCGCGATCTACCTGGCGAAGCGTGGCTACGAAGTGCACGTCGTGGAGAA<br>GCGTGGTGATCCTCTGCGCGATCTGAGCTCTTACGTGGACGTTGTTAGCA<br>GCCGTGCGATCGGCGTGAGCATGACCGTTCGTGGTATCAAGAGCGTTTT<br>GGCTGCGGGCATTCCGCGTGCAGAGCTGGATGCGTGTGGCGAACCGAT<br>CGTGGCAATGGCTTTCTCCGTGGGTGGTCAGTATCGCATGCGCGAACTG<br>AAGCCGTTGGAGGATTTCCGTCCGCTGAGCTTGAACCGTGCGGCGTTTC<br>AAAAGCTGCTGAACAAATACGCGAACCTGGCAGGCGTTCGTTACTACTTT<br>GAGCATAAGTGCCTGGATGTTGACCTGGATGGTAAGAGCGTGTTGATTCA<br>GGGCAAAGATGGTCAGCCGCAGCGTCTGCAAGGTGACATGATTATCGGT<br>GCGGATGGCGCCCACAGCGCCGTCCGTCAGGCGATGCAGAGCGGCCTG<br>CGTCGTTTCGAGTTCCAGCAAACGTTCTTCCGCCATGGCTACAAAACCCT<br>GGTTTTGCCGGACGCGCAAGCACTGGGTTACCGTAAAGACACGCTGTAC<br>TTTTTCGGCATGGATTCCGGTGGCCTGTTCGCGGGTCGTGCGGCTACGA<br>TCCCAGATGGTAGCGTCAGCATCGCCGTTTGCCTGCCGTACTCGGGTAG<br>CCCTTCCCTGACGACCACCGACGAACCGACGATGCGTGCGTTCTTCGAT<br>CGTTACTTCGGTGGCCTGCCGCGTGACGCGCGTGACGAAATGCTGCGTC<br>AGTTTCTGGCGAAGCCGAGCAACGACCTGATTAACGTGCGCTCTAGCACC<br>TTTCACTATAAGGGTAATGTGCTGTTGCTGGGTGATGCTGCGCATGCGAC<br>TGCGCCGTTCCTGGGTCAGGGTATGAACATGGCGCTGGAGGACGCCCGC<br>ACGTTTGTCGAGCTGCTGGACCGCCACCAGGGCGACCAAGACAAAGCCT |

| | |
|---|---|
| | TTCCGGAGTTCACGGAGCTGCGCAAAGTCCAGGCAGACGCAATGCAAGA<br>CATGGCTCGCGCCAACTATGACGTTTTGAGCTGCTCGAACCCGATCTTTT<br>TCATGCGTGCGCGTTACACGCGTTACATGCATTCCAAGTTTCCGGGCCTG<br>TATCCGCCGGATATGGCCGAGAAACTGTACTTTACGAGCGAGCCGTACGA<br>TCGTCTGCAACAAATCCAGCGTAAACAGAATGTTTGGTACAAGATTGGTC<br>GCGTGAATTGA |
| **vioD** | CAT**ATG**AAGATTCTGGTCATTGGTGCTGGTCCAGCTGGTCTGGTTTTCGC<br>ATCCCAACTGAAGCAGGCACGCCCTTTGTGGGCCATTGACATCGTGGAG<br>AAGAATGACGAGCAAGAAGTGCTGGGCTGGGGTGTCGTGCTGCCTGGCC<br>GTCCGGGTCAGCACCCGGCGAACCCGCTGTCCTATCTGGATGCACCGGA<br>GCGTCTGAATCCGCAATTTCTGGAGGACTTCAAACTGGTGCATCATAATG<br>AGCCGTCCTTGATGTCCACGGGCGTTTTGTTGTGCGGCGTGGAGCGTCG<br>CGGTCTGGTTCACGCGCTGCGCGATAAGTGCCGCAGCCAAGGCATTGCT<br>ATTCGTTTCGAAAGCCCGTTGCTGGAACACGGTGAGCTGCCGCTGGCGG<br>ACTATGATCTGGTGGTCCTGGCTAATGGTGTTAATCACAAAACCGCGCAT<br>TTCACCGAGGCTCTGGTCCCGCAGGTGGACTACGGCCGCAATAAGTACA<br>TTTGGTATGGCACTAGCCAGCTGTTCGATCAGATGAATCTGGTTTTTCGTA<br>CCCATGGTAAAGATATCTTTATCGCGCATGCCTATAAGTATAGCGATACCA<br>TGAGCACGTTCATTGTCGAATGTAGCGAAGAGACTTACGCACGCGCACGC<br>CTGGGCGAAATGTCCGAAGAGGCGAGCGCAGAATACGTTGCTAAGGTGT<br>TCCAGGCCGAGCTGGGTGGTCACGGCCTGGTGAGCCAGCCGGGTCTGG<br>GTTGGCGTAACTTCATGACGTTGTCTCATGACCGTTGTCATGATGGTAAGT<br>TGGTTCTGCTGGGTGACGCGCTGCAAAGCGGTCACTTTAGCATCGGCCA<br>CGGCACCACGATGGCCGTGGTGGTGGCGCAGCTGCTGGTTAAAGCGCT<br>GTGTACCGAAGATGGTGTGCCTGCCGCGCTGAAACGTTTCGAAGAGCGT<br>GCCCTGCCGCTGGTGCAGTTGTTCCGTGGCCACGCAGACAACAGCCGCG<br>TTTGGTTCGAAACCGTCGAAGAGCGCATGCACCTGTCCTCGGCGGAATTT<br>GTGCAAAGCTTCGACGCACGCCGCAAAAGCCTGCCGCCGATGCCGGAAG<br>CACTGGCGCAGAATCTGCGTTATGCTTTGCAGCGCTGA |
| **vioE** | CAT**ATG**GAGAACCGTGAGCCACCACTGTTGCCAGCCCGTTGGAGCAGCG<br>CCTATGTCTCTTATTGGAGCCCGATGCTGCCGGATGACCAGCTGACCAGC<br>GGCTATTGCTGGTTCGACTATGAACGTGACATCTGTCGTATTGACGGCCT<br>GTTCAATCCGTGGAGCGAGCGTGATACTGGTTATCGCCTGTGGATGTCG<br>GAGGTTGGTAATGCGGCCAGCGGCCGTACCTGGAAACAAAAAGTCGCCT<br>ATGGTCGTGAGCGTACCGCCCTGGGTGAACAGCTGTGTGAGCGTCCGCT<br>GGATGATGAGACTGGCCCTTTTGCCGAATTGTTCCTGCCACGCGATGTCC<br>TGCGCCGTCTGGGTGCCCGTCACATTGGCCGTCGCGTGGTTCTGGGTCG<br>CGAAGCGGACGGTTGGCGTTACCAGCGCCCAGGTAAAGGTCCGAGCACC<br>CTGTACCTGGATGCGGCGAGCGGCACTCCACTGCGCATGGTCACCGGCG<br>ATGAAGCGTCGCGTGCAAGCCTGCGTGATTTTCCGAATGTGAGCGAGGC<br>GGAGATCCCGGACGCGGTTTTCGCGGCCAAGCGCTAA |
| **RebF** | CAT**ATG**ACCATCGAATTTGATCGTCCGGGTGCACATGTTACCGCAGCAGA<br>TCATCGTGCCCTGATGAGCCTGTTTCCGACCGGTGTTGCAGTTATTACCG<br>CAATTGATGAAGCAGGTACACCGCATGGTATGACCTGTACCAGCCTGACC<br>AGCGTTACCCTGGACCCTCCGACCCTGCTGGTTTGTCTGAATCGTGCAAG<br>CGGCACCCTGCATGCCGTTCGTGGTGGTCGTTTTGGTGTTAATCTGCTGC<br>ATGCACGTGGTCGTCGTGCAGCAGAAGTTTTTAGCACCGCAGTTCAGGAT<br>CGCTTTGGTGAAGTTCGTTGGGAACATAGTGATGTTACCGGTATGCCGTG<br>GCTGGCCGAAGATGCACATGCATTTGCAGGTTGTGTTGTTCGTAAAAGCA<br>CCGTTGTTGGTGATCATGAAATTGTTCTGGGTGAAGTGCATGAAGTTGTTC<br>GTGAACATGATCTGCCGCTGCTGTATGGTATGCGTGAATTTGCAGTTTGG<br>ACACCGGAAGGTTAA |
| **RebH** | CAT**ATG**AGCGGCAAAATCGACAAAATTCTGATTGTTGGTGGTGGCACCGC<br>AGGTTGGATGGCAGCAAGCTATCTGGGTAAAGCACTGCAGGGTACAGCA<br>GATATTACCCTGCTGCAGGCACCGGATATTCCGACCCTGGGTGTTGGTGA<br>AGCAACCATTCCGAATCTGCAGACCGCATTTTTTGATTTTCTGGGTATTCC<br>GGAAGATGAATGGATGCGTGAATGTAATGCAAGCTATAAAGTGGCCATCA<br>AATTCATTAATTGGCGTACCGCAGGCGAAGGCACCAGCGAAGCACGTGA<br>ACTGGATGGTGGTCCGGATCATTTTTATCATAGCTTTGGTCTGCTGAAATA<br>CCATGAGCAGATTCCGCTGAGCCATTATTGGTTTGATCGTAGCTATCGTG |

| | |
|---|---|
| | GTAAAACCGTTGAACCGTTTGATTACGCCTGTTATAAAGAACCGGTTATTC<br>TGGATGCAAATCGTAGTCCGCGTCGTCTGGATGGTAGCAAAGTTACCAAT<br>TATGCATGGCATTTTGATGCACATCTGGTTGCAGATTTTCTGCGTCGTTTT<br>GCAACCGAAAAACTGGGTGTTCGTCATGTTGAAGATCGTGTTGAACATGT<br>GCAGCGTGATGCAAATGGTAATATTGAAAGCGTTCGTACCGCAACCGGTC<br>GTGTTTTTGATGCCGACCTGTTTGTTGATTGTAGCGGTTTTCGTGGCCTGC<br>TGATTAACAAAGCAATGGAAGAACCGTTTCTGGATATGAGCGATCATCTG<br>CTGAATGATAGCGCAGTTGCAACCCAGGTTCCGCATGATGATGATGCCAA<br>TGGTGTGGAACCGTTTACCAGCGCAATTGCAATGAAAAGCGGTTGGACCT<br>GGAAAATTCCGATGCTGGGTCGTTTTGGCACCGGTTATGTTTATAGCAGC<br>CGCTTTGCCACCGAAGATGAAGCAGTTCGTGAATTTTGTGAAATGTGGCA<br>TCTGGACCCGGAAACCCAGCCGCTGAATCGTATTCGTTTTCGTGTTGGTC<br>GTAATCGTCGTGCATGGGTTGGTAATTGTGTTAGCATTGGCACCAGCAGC<br>TGTTTTGTTGAACCGCTGGAAAGCACCGGTATCTATTTTGTTTATGCAGCA<br>CTGTATCAGCTGGTGAAACATTTTCCGGATAAAAGCCTGAATCCGGTTCT<br>GACCGCACGTTTTAATCGTGAAATTGAAACCATGTTCGATGACACCCGTG<br>ATTTTATTCAGGCCCACTTTTATTTCAGTCCGCGTACCGATACCCCGTTTT<br>GGCGTGCAAATAAAGAACTGCGTCTGGCAGATGGTATGCAAGAAAAAATT<br>GATATGTATCGTGCCGGTATGGCAATTAATGCACCGGCAAGTGATGATGC<br>ACAGCTGTATTATGGCAACTTTGAAGAAGAATTTCGCAACTTCTGGAACAA<br>CAGCAACTATTATTGTGTTCTGGCAGGTCTGGGTCTGGTTCCGGATGCAC<br>CGTCACCGCGTCTGGCCCACATGCCGCAGGCAACCGAATCAGTTGATGA<br>AGTTTTTGGTGCAGTTAAAGATCGTCAGCGTAACCTGCTGGAAACCCTGC<br>CGAGCCTGCATGAATTTCTGCGCCAGCAGCATGGTCGTTAA |
| **RebO** | CAT**ATG**AGCCGTGGTCATAAAAAGATTACCGTTCTGGGTGCCGGTGTTGC<br>AGGTCTGGTTGCAGCACATGAACTGGAAGAACTGGGTCATGAAGTTGAAG<br>TTCTGGAAGGTAGCGATCGTCTGGGTGGTCGTGTTCATACCCATCGTTTT<br>GGTGAAGGTGGTAGCGTTCCGTTTGTTGAACTGGGTGCAATGCGTATTCC<br>GACCAAACATCGTCATACCATTGATTATATTGGTAAACTGGGTCTGACCCC<br>GAAACTGAAAGAGTTTAAAACCCTGTTTAGTGATGATGGTGCCTATCATAC<br>CACCAGTGCAGGTTTTGTTCGTGTTCGTGATGCAGCAAAAGTTCTGGTTG<br>ATGAATTTCGTCTGCTGATGAGCGGTCGTGATCTGCGTGAAGAAACCATT<br>CTGTTTGGTGCATGGCTGACCGCAGTTGGTGATGCAATTGCACCGGCAG<br>ATTTTCGTGCAGCACTGCGTACCGATTTTACCGCAGATCTGCTGGAAGTT<br>GTTGATCGTATTGATCTGGACCCGTTTCTGGTTGGTGCAGCACGTGATCA<br>GTTTGATCTGCATGCATTTTTTGCAGCACATCCGGAAGTTCGTACCAGCTG<br>TACCGGTAAACTGAATCGTTTTGTTGATGATATCCTGGATGAAACCAGTCC<br>GCGTCTGCTGCGTCTGGAAGGTGGTATGGATCAGCTGGTTGATGCACTG<br>GTTGAACGTATTCGTGGTGATATTCGTACCGGTCATGAAGTGAGCGCAAT<br>TGATGTTCGTGAAGATCATGTTGCAGTTACCGTTCATAATGGTCATGGTGT<br>TAATACCCTGCGTAGCGATCATGTTCTGTGTACCATTCCGTTTAGCGTTCT<br>GCGTAATCTGCGTCTGACCGGTCTGAGCACCGATAAACTGGAAATTATTC<br>ACGACGTGAAATATTGGAGCGCAACCAAAGTTGCATTTCGTTGTCGTGAA<br>CCGTTTTGGGAACGTGATGGTATTAATGGTGGTGCAAGCTTTGGTGGTGG<br>TCGTATTCGTCAGACCTATTATCCGCCTGTTGAAGGTGATCCGACCCGTG<br>GTGCAGTTCTGCTGGCAAGCTATACCATGGGTGATGATGCAGATGTTCTG<br>GGTGGTATGCCGGAAGCACAGCGTCATGAAGTGGTTCTGGATGAAGTTG<br>GTCGTATGCATCCGGAACTGCATGAACCGGGTATGGTTGTTGAAGCAGTT<br>AGCCGTGCATGGGGTGAAGATCGTTGGAGCAATGGTGCGGGTGTTACCC<br>GTTGGGGTAAAGATGTTGCAGCATGTGAAGAAGAACGCGATCGCGCAGC<br>CCGTCCGGAAGGTCGTCTGTATTTTGCCGGTGAACATTGTAGCAGCACCA<br>CCGCATGGATTGATGGTGCAGTTGAAAGCGCACTGGCAGCAGTTCGTGC<br>AATTGAAGCCGGTGATGGTCGTGGATCCTCGAGCTAA |
| **RebD (with C-terminal *thrombin cleavage site* and His6 tag)** | CAT**ATG**AGCGTTTTTGATCTGCCTCGTCTGCATTTTGCAGGCACCGCAAC<br>CACCCGTCTGCCGACCGGTCCGCGTAATGGTCTGGTTGATCTGAGCACC<br>CATAGCGTTGTTATGGATGGTGAACGTTTTCCGGCAAGCCGTCCGGCAGC<br>AGAATATCATGCATATCTGGATCGTGTTGGTGGTAAAGGCACCGCATTTG<br>CAGGTAATGGTTATTTTGCAATTGATGCCGGTATTACCGCAGTTGAACGTG<br>CAGCCGGTGAAGTTGATACCGGTGATCTGCTGGTTGGTCGTGCAGTTGAT<br>GTTTGGGGTCATTATAACGAATATCTGGCCACCACCTTTAATCGTGCACGT |

| | |
|---|---|
| | ATTTTTGATGTTGATCCGAGCAGCAGCTGGACCAGCACCGTTATGATTGG<br>TCAGTTTGGTTTTGGTCGTCTGGGTCGTAGCCATGATGTTGGTTATGTTTT<br>TACCGGTGGTGTTCATGGTATGCAGCCTCCGCGTTGGCATGAAGATGGTC<br>GTGTTCTGCATCAGTTTACCGTTCCGGCAGGCGAAGATATGACCTGGTTT<br>GGTAGCGCAGCAGATTCACCGGCAGCAGCACGTCTGCGTGAACTGGTTG<br>AAAGCGGTGAAGCAGATGGTCTGGTGGTTCAGCTGGCACTGAGTGATGC<br>AGGTCCGGCACCGATGCCGCATGCACAGCAGTGGCGTCTGCGTGGCAC<br>CATTGCACCGTGGCATGCCGGTGAACCGCGTACCTGTCCGGCAGGTCGT<br>CTGCTGACACCGCATAATCTGACAGCAGATCTGCGTGGTGATCATGTTAG<br>CCTGAATCTGATTAGCTTTCGTCCGCCTACCGGTATTAGCGGTCTGGAAC<br>TGCGTACCGCAGATACCGATCGTTTTATTGCACGTGTTCCGGCAGATGAT<br>CCGCATGGTGTTGTTACAGTTCCGGCAGCGGAAGGTGGTGATGAAGCAC<br>TGTGTGTTGTTGGCACCACCGCAGCGGGTGAACGTATTGTTGTTAGCCGT<br>GAACGTGAAGTTACCGTTCATGTTGATGATGCAAGCGTTTTTCTGGAACAT<br>CCGCGTGGTCCGGGTGATAGCGATCAGGATGCAGAAATTGCAGTTCGTA<br>CCTATGTTCGTGGTGAACCGGCAGCCGCAACCATTCATATTGGTCAGTAT<br>TTCAATCCGCGTGCATTTCCGCTGGATGAACATGCAACCGCAGCAAGCGC<br>AACACCGGAAGATCTGGATGTTGTTGCCCTGTGTGTTGATGGTACACGTT<br>GGAGCCGTCATTGTGTTATTAGCACCGATGAAAATGGTGATGGTCGCTTT<br>CTGCTGCGTGGTGCACGTCCGGGTGCAACCCGTCTGCTGCTGAGCGCAG<br>AAGGTGCAACCCCGTTTGATGGTCTGACCGCAGCAGCAGCCTATGATAAT<br>GATGATAGCCTGGGTCTGTGGTCAGGTCTGGCAAGCGTTGCAGTTCGTG<br>TTCTGCCGGATCATTGGTGGATGGATGATATTCCGCGTGATAAAGTTACC<br>TTCGATCTGCTGTATCGTGAAGTGTTTGCATTTTATGAACTGCTGTATAGC<br>TTTATGGGCGAAGAAGTTTTTAGCCTGGCAGATCGTTTTCGTGTTGAAACC<br>CATCCGCGTCTGATTTGGCAGATGTGTGATCCGCGTAATCGTGCAAAAAC<br>CTATTATATGCCTCCGACCCGTGATCTGACCGGTCCGCAGGCACGTCTGC<br>TGTTAGCATATCTGCGTGCACAGAATAGTGATGTTGTTGTTCCGGTTATTG<br>AACCGAGCCATACCCGTAGCGGCACCCCGATTAGCACCCGTACCGATCT<br>GGTTCGTGCACTGCGTCATGGTGTTGCAATTGAACTGGCAGTTATGCTGC<br>AGTATCTGTATGCAGCATTTAGCATTCCGACCCATGGTGCAGGTCAAGAA<br>CTGGTTAGCCGTGGTGATTGGACACCGGAACAGCTGCGTCTGATGTGTG<br>GTGATGGTGGTGAAACCACCGATGGTGGTGTGCGTGGTAGCCTGCTGGG<br>TGTTGCACGTGAAGAAATGATTCATTTTCTGGTGGTGAATAATGTTCTGAT<br>GGCAGTTGGTGAACCGTTTCATGTTCCGGATCTGGATTTTGGCACCATTA<br>ATGATACCCTGATGGTTCCGCTGGATTTTAGCCTGGAAGCACTGGGTCTG<br>GGTAGCGTTCAGCGTTTTATTCAGATTGAACAGCCGGAAGGTCTGACCGG<br>TGCAGTTCGTCTGGGTGATCTGCCGGTTCCGGTTCGTGAAGCCGAAGATT<br>TTCATTATGCAAGCCTGAGCGAACTGTATGGTGATATTCGTGAAGGTCTG<br>CAGCGTGTTCCGGGTCTGTTTCTGGTTGAACGTGGTCGTGGTGGTGGTG<br>AACATCACCTGTTTCTGCGTGAAAGCGTTAATGCAGTTCATCCGGATTATC<br>AGCTGGAAGTTGATGATCTGAGCAGCGCACTGTTTGCCATTGATTTTGTTA<br>CCGAACAGGGTGAAGGTCATGTTCTGACCGATGAAGATACCGGTGAAGA<br>AAGCCATTATGATACCTTTGTTCGTGTTGCCGATCTGCTGATGAAAGAACG<br>CCTGACCGCAGCCGATACCCGTCGTGCACAGTGGTCACCGGCATATCCG<br>GTTGCACGTAATCCGACCGTTCATGGTGGTGGTCAGAGCAAAGAACTGGT<br>GACCAGTCCGGTTGCCCGTGAACTGATGGTTCTGTTTAACAAAAGCTACT<br>TCATGATGCTGCAGCTGATGGTTCAGCATTTTGGTGGTAGTCCGGATGCA<br>AGCCTGCGTCGTAGCAAACTGATGAATGCAGCAATTGATGTTATGACCGG<br>TGTTATGCGTCCGCTGGCAGAACTGCTGGTTACCGTTCCGAGCGGTCGT<br>CATGGTCGTACCGCAGGTCCGTCATTTGAACTGGATGAAAAACCGGCATT<br>TATTCCGCGTGCAGATGTTGCACGTCGTGCAATTAGCCTGCGTTTTCGTC<br>ATCTGGCAGAAAGCGCACGTACCTGTGCACTGGTTCCGGATAAAGTTGTT<br>CGTAATCTGGATTTTCTGGCAGATCAGTTTGCAACCGAAGGTCCGCGTGG<br>ATCCTCGAGC*ctggtgccgcgcggcagc*catcatcatcatcatcacTAA |

**EcoFlex vectors**

Promoters, RBSs and terminators are cloned into pBP vector between NdeI and SphI sites. Genes are cloned into pBP-ORF vector between NdeI and BamHI sites.

Destination vectors pTU1, pTU2 and pTU3 contains negative selection marker (RFP with constitutive promoter) flanked by corresponding 4bp linker that directs assembly.

Destination vectors pTU2S contains both main Golden Gate assembly site (cut by BsmBI) as well as a secondary cloning site which is cut by BpiI. All vectors (except the pTU2S vectors) are available from the EcoFlex kit on Addgene.

**Primers for sequencing**

| Name | Sequence |
|---|---|
| **VF2** | CTGACGTCTAAGAAACCATT |
| **VR** | AACCGTATTACCGCCTTTGA |
| **VioA_Rev** | TTGCTGCGGATACGGCCACC |
| **VioB_Fwr** | GTTTCACCCGGGTCCGTTTG |
| **VioB_Rev** | ATAACCCGGATGCGCACGGTTG |
| **VioC_Fwr** | GAAACGTGCGATTATCGTTGGTG |
| **VioC_Fwr_2** | TGACGAAATGCTGCGTCAG |
| **VioD_Fwr** | GAAGATTCTGGTCATTGGTGCTG |
| **VioD_Fwr_2** | TGCAAAGCGGTCACTTTAGC |
| **VioE_Fwr** | AACCGTGAGCCACCACTGTTG |
| **RebD_Fwr** | GAGCGTTTTTGATCTGCCTCG |
| **RebD_Fwr_2** | GAATCTGATTAGCTTTCGTCCGC |
| **RebD_Fwr_3** | GCGTCTGATTTGGCAGATGTG |
| **RebD_Fwr_4** | GGAAGTTGATGATCTGAGCAGC |
| **RebH_Fwr** | GAGCGGCAAAATCGACAAAATTC |
| **RebH_Fwr2** | GCAATGAAAAGCGGTTGGAC |
| **RebO_Fwr** | GCCGTGGTCATAAAAAGATTACCG |
| **RebO_Fwr2** | CCTGCGTAGCGATCATGTTC |

### SI Appendices – Plasmid and protein sequence for Rebeccamycin-Violacein hybrid pathway

Sequences are supplied as gbk file which can be imported and visualized on various molecular biology software such as Genious, Snapgene or Benchling.

#### pTU3-A-RebFHOD-VioCDE-KanR

```
LOCUS       Exported File         14756 bp ds-DNA    circular SYN 22-SEP-2017
DEFINITION  .
ACCESSION   .
VERSION     .
KEYWORDS    pTU3-A-RebFHOD_VioCDE_KanR
SOURCE      synthetic DNA construct
  ORGANISM  synthetic DNA construct
REFERENCE   1  (bases 1 to 14756)
  AUTHORS   .
  TITLE     Direct Submission
  JOURNAL   Exported 22 Sep 2017 from SnapGene 1.1.3
            http://www.snapgene.com
COMMENT
FEATURES             Location/Qualifiers
     source          1..14756
                     /organism="synthetic DNA construct"
                     /mol_type="other DNA"
     misc_feature    1..6
                     /note="BBa Suffix"
                     /note="color: #ff00ff"
     misc_feature    70..89
                     /note="VR Primer"
                     /note="color: #00ffff"
     rep_origin      complement(165..847)
                     /direction=LEFT
                     /note="ColE1 origin"
                     /note="color: #808080"
     misc_feature    1931..1950
                     /note="VF2 Primer"
                     /note="color: #00ffff"
     misc_feature    2058..2063
                     /note="BBa Prefix"
                     /note="color: #ff00ff"
     misc_feature    2080..2083
                     /note="Scar"
                     /note="color: #00ffff"
     CDS             2150..2671
                     /note="RebF_reductase"
                     /note="color: #ffc0cb; direction: RIGHT"
     misc_feature    2809..2812
                     /note="Scar"
                     /note="color: #00ffff"
     CDS             2882..4480
                     /note="halogenase_RebH"
                     /note="color: #ffc0cb; direction: RIGHT"
     misc_feature    2963..2968
                     /note="BBa Suffix"
                     /note="color: #ff00ff"
     misc_feature    2990..2995
                     /note="BBa Suffix"
                     /note="color: #ff00ff"
```

```
     misc_feature    3044..3049
                     /note="BBa Suffix"
                     /note="color: #ff00ff"
     misc_feature    4618..4621
                     /note="Scar"
                     /note="color: #00ffff"
     CDS             4691..6124
                     /codon_start=1
                     /note="RebO7ClTrpoxidaseQ8K"
                     /note="color: #993366"
                     /translation="MSRGHKKITVLGAGVAGLVAAHELEELGHEVEVLEGSDRLGGRVH
                     THRFGEGGSVPFVELGAMRIPTKHRHTIDYIGKLGLTPKLKEFKTLFSDDGAYHTTSAG
                     FVRVRDAAKVLVDEFRLLMSGRDLREETILFGAWLTAVGDAIAPADFRAALRTDFTADL
                     LEVVDRIDLDPFLVGAARDQFDLHAFFAAHPEVRTSCTGKLNRFVDDILDETSPRLLRL
                     EGGMDQLVDALVERIRGDIRTGHEVSAIDVREDHVAVTVHNGHGVNTLRSDHVLCTIPF
                     SVLRNLRLTGLSTDKLEIIHDVKYWSATKVAFRCREPFWERDGINGGASFGGGRIRQTY

YPPVEGDPTRGAVLLASYTMGDDADVLGGMPEAQRHEVVLDEVGRMHPELHEPGMVVEA

VSRAWGEDRWSNGAGVTRWGKDVAACEEERDRAARPEGRLYFAGEHCSSTTAWIDGAVE
                     SALAAVRAIEAGDGRGSSS"
     misc_feature    6258..6261
                     /note="Scar"
                     /note="color: #00ffff"
     CDS             6331..9420
                     /codon_start=1
                     /note="RebDDichlorochromopy"
                     /note="color: #993366"
                     /translation="MSVFDLPRLHFAGTATTRLPTGPRNGLVDLSTHSVVMDGERFPAS

RPAAEYHAYLDRVGGKGTAFAGNGYFAIDAGITAVERAAGEVDTGDLLVGRAVDVWGHY

NEYLATTFNRARIFDVDPSSSWTSTVMIGQFGFGRLGRSHDVGYVFTGGVHGMQPPRWH

EDGRVLHQFTVPAGEDMTWFGSAADSPAAARLRELVESGEADGLVVQLALSDAGPAPMP
                     HAQQWRLRGTIAPWHAGEPRTCPAGRLLTPHNLTADLRGDHVSLNLISFRPPTGISGLE

LRTADTDRFIARVPADDPHGVVTVPAAEGGDEALCVVGTTAAGERIVVSREREVTVHVD
                     DASVFLEHPRGPGDSDQDAEIAVRTYVRGEPAAATIHIGQYFNPRAFPLDEHATAASAT

PEDLDVVALCVDGTRWSRHCVISTDENGDGRFLLRGARPGATRLLLSAEGATPFDGLTA

AAAYDNDDSLGLWSGLASVAVRVLPDHWWMDDIPRDKVTFDLLYREVFAFYELLYSFMG

EEVFSLADRFRVETHPRLIWQMCDPRNRAKTYYMPPTRDLTGPQARLLLAYLRAQNSDV
                     VVPVIEPSHTRSGTPISTRTDLVRALRHGVAIELAVMLQYLYAAFSIPTHGAGQELVSR

GDWTPEQLRLMCGDGGETTDGGVRGSLLGVAREEMIHFLVVNNVLMAVGEPFHVPDLDF
                     GTINDTLMVPLDFSLEALGLGSVQRFIQIEQPEGLTGAVRLGDLPVPVREAEDFHYASL
                     SELYGDIREGLQRVPGLFLVERGRGGGEHHLFLRESVNAVHPDYQLEVDDLSSALFAID

FVTEQGEGHVLTDEDTGEESHYDTFVRVADLLMKERLTAADTRRAQWSPAYPVARNPTV

HGGGQSKELVTSPVARELMVLFNKSYFMMLQLMVQHFGGSPDASLRRSKLMNAAIDVMT
                     GVMRPLAELLVTVPSGRHGRTAGPSFELDEKPAFIPRADVARRAISLRFRHLAESARTC
                     ALVPDKVVRNLDFLADQFATEGPRGSSSLVPRGSHHHHHH"
     misc_feature    9554..9557
                     /note="Scar"
                     /note="color: #00ffff"
     CDS             9635..10924
```

/codon_start=1
/note="VioC"
/note="color: #00ffff"
/translation="MKRAIIVGGGLAGGLTAIYLAKRGYEVHVVEKRGDPLRDLSSYVD
VVSSRAIGVSMTVRGIKSVLAAGIPRAELDACGEPIVAMAFSVGGQYRMRELKPLEDFR
PLSLNRAAFQKLLNKYANLAGVRYYFEHKCLDVDLDGKSVLIQGKDGQPQRLQGDMIIG

ADGAHSAVRQAMQSGLRRFEFQQTFFRHGYKTLVLPDAQALGYRKDTLYFFGMDSGGLF
AGRAATIPDGSVSIAVCLPYSGSPSLTTTDEPTMRAFFDRYFGGLPRDARDEMLRQFLA

KPSNDLINVRSSTFHYKGNVLLLGDAAHATAPFLGQGMNMALEDARTFVELLDRHQGDQ

DKAFPEFTELRKVQADAMQDMARANYDVLSCSNPIFFMRARYTRYMHSKFPGLYPPDMA
EKLYFTSEPYDRLQQIQRKQNVWYKIGRVN"
CDS 11141..12262
/codon_start=1
/note="VioD"
/note="color: #00ff00"
/translation="MKILVIGAGPAGLVFASQLKQARPLWAIDIVEKNDEQEVLGWGVV

LPGRPGQHPANPLSYLDAPERLNPQFLEDFKLVHHNEPSLMSTGVLLCGVERRGLVHAL
RDKCRSQGIAIRFESPLLEHGELPLADYDLVVLANGVNHKTAHFTEALVPQVDYGRNKY
IWYGTSQLFDQMNLVFRTHGKDIFIAHAYKYSDTMSTFIVECSEETYARARLGEMSEEA

SAEYVAKVFQAELGGHGLVSQPGLGWRNFMTLSHDRCHDGKLVLLGDALQSGHFSIGHG

TTMAVVVAQLLVKALCTEDGVPAALKRFEERALPLVQLFRGHADNSRVWFETVEERMHL
SSAEFVQSFDARRKSLPPMPEALAQNLRYALQR"
CDS 12479..13054
/codon_start=1
/note="VioE"
/note="color: #ffff00"
/translation="MENREPPLLPARWSSAYVSYWSPMLPDDQLTSGYCWFDYERDICR

IDGLFNPWSERDTGYRLWMSEVGNAASGRTWKQKVAYGRERTALGEQLCERPLDDETGP

FAELFLPRDVLRRLGARHIGRRVVLGREADGWRYQRPGKGPSTLYLDAASGTPLRMVTG
DEASRASLRDFPNVSEAEIPDAVFAAKR"
CDS complement(13384..14178)
/codon_start=1
/note="KanR"
/note="color: #999999"
/translation="MIEQDGLHAGSPAAWVERLFGYDWAQQTIGCSDAAVFRLSAQGRP

VLFVKTDLSGALNELQDEAARLSWLATTGVPCAAVLDVVTEAGRDWLLLGEVPGQDLLS

SHLAPAEKVSIMADAMRRLHTLDPATCPFDHQAKHRIERARTRMEAGLVDQDDLDEEHQ

GLAPAELFARLKARMPDGEDLVVTHGDACLPNIMVENGRFSGFIDCGRLGVADRYQDIA
LATRDIAEELGGEWADRFLVLYGIAAPDSQRIAFYRLLDEFF"
ORIGIN
1 ctgcaggctt cctcgctcac tgactcgctg cgctcggtcg ttcggctgcg gcgagcggta
61 tcagctcact caaaggcggt aatacggtta tccacagaat caggggataa cgcaggaaag
121 aacatgtgag caaaaggcca gcaaaaggcc aggaaccgta aaaaggccgc gttgctggcg
181 tttttccata ggctccgccc ccctgacgag catcacaaaa atcgacgctc aagtcagagg
241 tggcgaaacc cgacaggact ataaagatac caggcgtttc cccctggaag ctccctcgtg
301 cgctctcctg ttccgaccct gccgcttacc ggatacctgt ccgcctttct cccttcggga
361 agcgtggcgc tttctcatag ctcacgctgt aggtatctca gttcggtgta ggtcgttcgc
421 tccaagctgg gctgtgtgca cgaacccccc gttcagcccg accgctgcgc cttatccggt
481 aactatcgtc ttgagtccaa cccggtaaga cacgacttat cgccactggc agcagccact

541 ggtaacagga ttagcagagc gaggtatgta ggcggtgcta cagagttctt gaagtggtgg
601 cctaactacg gctacactag aaggacagta tttggtatct gcgctctgct gaagccagtt
661 accttcggaa aaagagttgg tagctcttga tccggcaaac aaaccaccgc tggtagcggt
721 ggtttttttg tttgcaagca gcagattacg cgcagaaaaa aaggatctca agaagatcct
781 ttgatctttt ctacggggtc tgacgctcag tggaacgaaa actcacgtta agggattttg
841 gtcatgagat tatcaaaaag gatcttcacc tagatccttt taaattaaaa atgaagtttt
901 aaatcaatct aaagtatata tgagtaaact tggtctgaca gttaccaatg cttaatcagt
961 gaggcaccta tctcagcgat ctgtctattt cgttcatcca tagttgcctg actccccgtc
1021 gtgtagataa ctacgatacg ggagggctta ccatctggcc ccagtgctgc aatgataccg
1081 cgagagccac gctcaccggc tccagattta tcagcaataa accagccagc cggaagggcc
1141 gagcgcagaa gtggtcctgc aactttatcc gcctccatcc agtctattaa ttgttgccgg
1201 gaagctagag taagtagttc gccagttaat agtttgcgca acgttgttgc cattgctaca
1261 ggcatcgtgg tgtcacgctc gtcgtttggt atggcttcat tcagctccgg ttcccaacga
1321 tcaaggcgag ttacatgatc ccccatgttg tgcaaaaaag cggttagctc cttcggtcct
1381 ccgatcgttg tcagaagtaa gttggccgca gtgttatcac tcatggttat ggcagcactg
1441 cataattctc ttactgtcat gccatccgta agatgctttt ctgtgactgg tgagtactca
1501 accaagtcat tctgagaata gtgtatgcgg cgaccgagtt gctcttgccc ggcgtcaata
1561 cgggataata ccgcgccaca tagcagaact ttaaaagtgc tcatcattgg aaaacgttct
1621 tcggggcgaa aactctcaag gatcttaccg ctgttgagat ccagttcgat gtaacccact
1681 cgtgcaccca actgatcttc agcatctttt actttcacca gcgtttctgg gtgagcaaaa
1741 acaggaaggc aaaatgccgc aaaaaaggga ataagggcga cacggaaatg ttgaatactc
1801 atactcttcc tttttcaata ttattgaagc atttatcagg gttattgtct catgagcgga
1861 tacatatttg aatgtattta gaaaaataaa caaatagggg ttccgcgcac atttccccga
1921 aaagtgccac ctgacgtcta agaaaccatt attatcatga cattaaccta taaaaatagg
1981 cgtatcacga ggcagaattt cagataaaaa aaatccttag ctttcgctaa ggatgatttc
2041 tggaattcgc ggccgcttct agaacgtctc aatctctata tctctatttt atggctagct
2101 cagtcctagg tacaatgcta gcgtacttta actttaagaa ggagatatac atatgaccat
2161 cgaatttgat cgtccgggtg cacatgttac cgcagcagat catcgtgccc tgatgagcct
2221 gtttccgacc ggtgttgcag ttattaccgc aattgatgaa gcaggtacac cgcatggtat
2281 gacctgtacc agcctgacca gcgttaccct ggaccctccg accctgctgg tttgtctgaa
2341 tcgtgcaagc ggcaccctgc atgccgttcg tggtggtcgt tttggtgtta atctgctgca
2401 tgcacgtggt cgtcgtgcag cagaagtttt tagcaccgca gttcaggatc gctttggtga
2461 agttcgttgg gaacatagtg atgttaccgg tatgccgtgg ctggccgaag atgcacatgc
2521 atttgcaggt tgtgttgttc gtaaaagcac cgttgttggt gatcatgaaa ttgttctggg
2581 tgaagtgcat gaagttgttc gtgaacatga tctgccgctg ctgtatggta tgcgtgaatt
2641 tgcagtttgg acaccggaag gttaaggatc ctcgaccagg catcaaataa aacgaaaggc
2701 tcagtcgaaa gactgggcct ttcgttttat ctgttgtttg tcggtgaacg ctctctacta
2761 gagtcacact ggctcacctt cgggtgggcc tttctgcgtt tatatgtttg ccctatttta
2821 tggctagctc agtcctaggt acaatgctag cgtactttaa ctttaagaag gagatataca
2881 tatgagcggc aaaatcgaca aaattctgat tgttggtggt ggcaccgcag gttggatggc
2941 agcaagctat ctgggtaaag cactgcaggg tacagcagat attaccctgc tgcaggcacc
3001 ggatattccg accctgggtg ttggtgaagc aaccattccg aatctgcaga ccgcattttt
3061 tgattttctg ggtattccgg aagatgaatg gatgcgtgaa tgtaatgcaa gctataaagt
3121 ggccatcaaa ttcattaatt ggcgtaccgc aggcgaaggc accagcgaag cacgtgaact
3181 ggatggtggt ccggatcatt ttatcatag ctttggtctg ctgaaatacc atgagcagat
3241 tccgctgagc cattattggt ttgatcgtag ctatcgtggt aaaaccgttg aaccgtttga
3301 ttacgcctgt tataaagaac cggttattct ggatgcaaat cgtagtccgc gtcgtctgga
3361 tggtagcaaa gttaccaatt atgcatggca ttttgatgca catctggttg cagattttct
3421 gcgtcgtttt gcaaccgaaa aactgggtgt tcgtcatgtt gaagatcgtg ttgaacatgt
3481 gcagcgtgat gcaaatggta atattgaaag cgttcgtacc gcaaccggtc gtgtttttga
3541 tgccgacctg tttgttgatt gtagcggttt tcgtggcctg ctgattaaca aagcaatgga
3601 agaaccgttt ctggatatga gcgatcatct gctgaatgat agcgcagttg caacccaggt
3661 tccgcatgat gatgatgcca atggtgtgga accgtttacc agcgcaattg caatgaaaag
3721 cggttggacc tggaaaattc cgatgctggg tcgttttggc accggttatg tttatagcag
3781 ccgctttgcc accgaagatg aagcagttcg tgaattttgt gaaatgtggc atctggaccc
3841 ggaaacccag ccgctgaatc gtattcgttt tcgtgttggt cgtaatcgtc gtgcatgggt
3901 tggtaattgt gttagcattg gcaccagcag ctgttttgtt gaaccgctgg aaagcaccgg
3961 tatctatttt gtttatgcag cactgtatca gctggtgaaa cattttccgg ataaaagcct
4021 gaatccggtt ctgaccgcac gttttaatcg tgaaattgaa accatgttcg atgacacccg
4081 tgattttatt caggcccact tttatttcag tccgcgtacc gataccccgt tttggcgtgc

4141 aaataaagaa ctgcgtctgg cagatggtat gcaagaaaaa attgatatgt atcgtgccgg
4201 tatggcaatt aatgcaccgg caagtgatga tgcacagctg tattatggca actttgaaga
4261 agaatttcgc aacttctgga acaacagcaa ctattattgt gttctggcag gtctgggtct
4321 ggttccggat gcaccgtcac cgcgtctggc ccacatgccg caggcaaccg aatcagttga
4381 tgaagttttt ggtgcagtta aagatcgtca gcgtaacctg ctggaaaccc tgccgagcct
4441 gcatgaattt ctgcgccagc agcatggtcg ttaaggatcc tcgaccaggc atcaaataaa
4501 acgaaaggct cagtcgaaag actgggcctt tcgttttatc tgttgtttgt cggtgaacgc
4561 tctctactag agtcacactg gctcaccttc gggtgggcct ttctgcgttt atatgttccg
4621 gctattttat ggctagctca gtcctaggta caatgctagc gtactttaac tttaagaagg
4681 agatatacat atgagccgtg gtcataaaaa gattaccgtt ctgggtgccg gtgttgcagg
4741 tctggttgca gcacatgaac tggaagaact gggtcatgaa gttgaagttc tggaaggtag
4801 cgatcgtctg ggtggtcgtg ttcataccca tcgttttggt gaaggtggta gcgttccgtt
4861 tgttgaactg ggtgcaatgc gtattccgac caaacatcgt cataccattg attatattgg
4921 taaactgggt ctgaccccga aactgaaaga gtttaaaacc ctgtttagtg atgatggtgc
4981 ctatcatacc accagtgcag gttttgttcg tgttcgtgat gcagcaaaag ttctggttga
5041 tgaatttcgt ctgctgatga gcggtcgtga tctgcgtgaa gaaaccattc tgtttggtgc
5101 atggctgacc gcagttggtg atgcaattgc accggcagat ttcgtgcag cactgcgtac
5161 cgattttacc gcagatctgc tggaagttgt tgatcgtatt gatctggacc cgtttctggt
5221 tggtgcagca cgtgatcagt ttgatctgca tgcatttttt gcagcacatc cggaagttcg
5281 taccagctgt accggtaaac tgaatcgttt tgttgatgat atcctggatg aaaccagtcc
5341 gcgtctgctg cgtctggaag gtggtatgga tcagctggtt gatgcactgg ttgaacgtat
5401 tcgtggtgat attcgtaccg gtcatgaagt gagcgcaatt gatgttcgtg aagatcatgt
5461 tgcagttacc gttcataatg gtcatggtgt taataccctg cgtagcgatc atgttctgtg
5521 taccattccg tttagcgttc tgcgtaatct gcgtctgacc ggtctgagca ccgataaact
5581 ggaaattatt cacgacgtga aatattggag cgcaaccaaa gttgcatttc gttgtcgtga
5641 accgttttgg gaacgtgatg gtattaatgg tggtgcaagc tttggtggtg gtcgtattcg
5701 tcagacctat tatccgcctg ttgaaggtga tccgacccgt ggtgcagttc tgctggcaag
5761 ctataccatg ggtgatgatg cagatgttct gggtggtatg ccggaagcac agcgtcatga
5821 agtggttctg gatgaagttg gtcgtatgca tccggaactg catgaaccgg gtatggttgt
5881 tgaagcagtt agccgtgcat ggggtgaaga tcgttggagc aatggtgcgg gtgttacccg
5941 ttggggtaaa gatgttgcag catgtgaaga agaacgcgat cgcgcagccc gtccggaagg
6001 tcgtctgtat tttgccggtg aacattgtag cagcaccacc gcatggattg atggtgcagt
6061 tgaaagcgca ctggcagcag ttcgtgcaat tgaagccggt gatggtcgtg gatcctcgag
6121 ctaaccaggc atcaaataaa acgaaaggct cagtcgaaag actgggcctt tcgttttatc
6181 tgttgtttgt cggtgaacgc tctctactag agtcacactg gctcaccttc gggtgggcct
6241 ttctgcgttt atatgttgaa gctattttat ggctagctca gtcctaggta caatgctagc
6301 gtactttaac tttaagaagg agatatacat atgagcgttt ttgatctgcc tcgtctgcat
6361 tttgcaggca ccgcaaccac ccgtctgccg accggtccgc gtaatggtct ggttgatctg
6421 agcacccata gcgttgttat ggatggtgaa cgttttccgg caagccgtcc ggcagcagaa
6481 tatcatgcat atctggatcg tgttggtggt aaaggcaccg catttgcagg taatggttat
6541 tttgcaattg atgccggtat taccgcagtt gaacgtgcag ccggtgaagt tgataccggt
6601 gatctgctgg ttggtcgtgc agttgatgtt tggggtcatt ataacgaata tctggccacc
6661 acctttaatc gtgcacgtat ttttgatgtt gatccgagca gcagctggac cagcaccgtt
6721 atgattggtc agtttggttt tggtcgtctg ggtcgtagcc atgatgttgg ttatgttttt
6781 accggtggtg ttcatggtat gcagcctccg cgttggcatg aagatggtcg tgttctgcat
6841 cagtttaccg ttccggcagg cgaagatatg acctggtttg gtagcgcagc agattcaccg
6901 gcagcagcac gtctgcgtga actggttgaa agcggtgaag cagatggtct ggtggttcag
6961 ctggcactga gtgatgcagg tccggcaccg atgccgcatg cacagcagtg gcgtctgcgt
7021 ggcaccattg caccgtggca tgccggtgaa ccgcgtacct gtccggcagg tcgtctgctg
7081 acaccgcata atctgacagc agatctgcgt ggtgatcatg ttagcctgaa tctgattagc
7141 tttcgtccgc ctaccggtat tagcggtctg gaactgcgta ccgcagatac cgatcgtttt
7201 attgcacgtg ttccggcaga tgatccgcat ggtgttgtta cagttccggc agcggaaggt
7261 ggtgatgaag cactgtgtgt tgttggcacc accgcagcgg gtgaacgtat tgttgttagc
7321 cgtgaacgtg aagttaccgt tcatgttgat gatgcaagcg tttttctgga acatccgcgt
7381 ggtccgggtg atagcgatca ggatgcagaa attgcagttc gtacctatgt tcgtggtgaa
7441 ccggcagccg caaccattca tattggtcag tatttcaatc cgcgtgcatt tccgctggat
7501 gaacatgcaa ccgcagcaag cgcaacaccg gaagatctgg atgttgttgc cctgtgtgtt
7561 gatggtacac gttggagccg tcattgtgtt attagcaccg atgaaaatgg tgatggtcgc
7621 tttctgctgc gtggtgcacg tccgggtgca acccgtctgc tgctgagcgc agaaggtgca
7681 accccgtttg atggtctgac cgcagcagca gcctatgata atgatgatag cctgggtctg

7741 tggtcaggtc tggcaagcgt tgcagttcgt gttctgccgg atcattggtg gatggatgat
7801 attccgcgtg ataaagttac cttcgatctg ctgtatcgtg aagtgtttgc attttatgaa
7861 ctgctgtata gctttatggg cgaagaagtt tttagcctgg cagatcgttt tcgtgttgaa
7921 acccatccgc gtctgatttg gcagatgtgt gatccgcgta atcgtgcaaa aacctattat
7981 atgcctccga cccgtgatct gaccggtccg caggcacgtc tgctgttagc atatctgcgt
8041 gcacagaata gtgatgttgt tgttccggtt attgaaccga gccatacccg tagcggcacc
8101 ccgattagca cccgtaccga tctggttcgt gcactgcgtc atggtgttgc aattgaactg
8161 gcagttatgc tgcagtatct gtatgcagca tttagcattc cgacccatgg tgcaggtcaa
8221 gaactggtta gccgtggtga ttggacaccg gaacagctgc gtctgatgtg tggtgatggt
8281 ggtgaaacca ccgatggtgg tgtgcgtggt agcctgctgg gtgttgcacg tgaagaaatg
8341 attcattttc tggtggtgaa taatgttctg atggcagttg gtgaaccgtt tcatgttccg
8401 gatctggatt ttggcaccat taatgatacc ctgatggttc cgctggattt tagcctggaa
8461 gcactgggtc tgggtagcgt tcagcgtttt attcagattg aacagccgga aggtctgacc
8521 ggtgcagttc gtctgggtga tctgccggtt ccggttcgtg aagccgaaga ttttcattat
8581 gcaagcctga gcgaactgta tggtgatatt cgtgaaggtc tgcagcgtgt tccgggtctg
8641 tttctggttg aacgtggtcg tggtggtggt gaacatcacc tgtttctgcg tgaaagcgtt
8701 aatgcagttc atccggatta tcagctggaa gttgatgatc tgagcagcgc actgtttgcc
8761 attgattttg ttaccgaaca gggtgaaggt catgttctga ccgatgaaga taccggtgaa
8821 gaaagccatt atgatacctt tgttcgtgtt gccgatctgc tgatgaaaga acgcctgacc
8881 gcagccgata cccgtcgtgc acagtggtca ccggcatatc cggttgcacg taatccgacc
8941 gttcatggtg gtggtcagag caaagaactg gtgaccagtc cggttgcccg tgaactgatg
9001 gttctgttta acaaaagcta cttcatgatg ctgcagctga tggttcagca ttttggtggt
9061 agtccggatg caagcctgcg tcgtagcaaa ctgatgaatg cagcaattga tgttatgacc
9121 ggtgttatgc gtccgctggc agaactgctg gttaccgttc cgagcggtcg tcatggtcgt
9181 accgcaggtc cgtcatttga actggatgaa aaaccggcat ttattccgcg tgcagatgtt
9241 gcacgtcgtg caattagcct gcgttttcgt catctggcag aaagcgcacg tacctgtgca
9301 ctggttccgg ataaagttgt tcgtaatctg gattttctgg cagatcagtt tgcaaccgaa
9361 ggtccgcgtg gatcctcgag cctggtgccg cgcggcagcc atcatcatca tcatcactaa
9421 ccaggcatca aataaaacga aaggctcagt cgaaagactg ggcctttcgt tttatctgtt
9481 gtttgtcggt gaacgctctc tactagagtc acactggctc accttcgggt gggcctttct
9541 gcgtttatat gttttaggta catctctatt ttatggctag ctcagtccta ggtacaatgc
9601 tagcgtactt taactttaag aaggagatat acatatgaaa cgtgcgatta tcgttggtgg
9661 cggcctggcg ggtggcctga ccgcgatcta cctggcgaag cgtggctacg aagtgcacgt
9721 cgtggagaag cgtggtgatc ctctgcgcga tctgagctct tacgtggacg ttgttagcag
9781 ccgtgcgatc ggcgtgagca tgaccgttcg tggtatcaag agcgttttgg ctgcgggcat
9841 tccgcgtgca gagctggatg cgtgtggcga accgatcgtg gcaatggctt tctccgtggg
9901 tggtcagtat cgcatgcgcg aactgaagcc gttggaggat ttccgtccgc tgagcttgaa
9961 ccgtgcggcg tttcaaaagc tgctgaacaa atacgcgaac ctggcaggcg ttcgttacta
10021 ctttgagcat aagtgcctgg atgttgacct ggatggtaag agcgtgttga ttcagggcaa
10081 agatggtcag ccgcagcgtc tgcaaggtga catgattatc ggtgcggatg gcgcccacag
10141 cgccgtccgt caggcgatgc agagcggcct gcgtcgtttc gagttccagc aaacgttctt
10201 ccgccatggc tacaaaaccc tggttttgcc ggacgcgcaa gcactgggtt accgtaaaga
10261 cacgctgtac tttttcggca tggattccgg tggcctgttc gcgggtcgtg cggctacgat
10321 cccagatggt agcgtcagca tcgccgtttg cctgccgtac tcgggtagcc cttccctgac
10381 gaccaccgac gaaccgacga tgcgtgcgtt cttcgatcgt tacttcggtg gcctgccgcg
10441 tgacgcgcgt gacgaaatgc tgcgtcagtt tctggcgaag ccgagcaacg acctgattaa
10501 cgtgcgctct agcacctttc actataaggg taatgtgctg ttgctgggtg atgctgcgca
10561 tgcgactgcg ccgttcctgg gtcagggtat gaacatggcg ctggaggacg cccgcacgtt
10621 tgtcgagctg ctggaccgcc accagggcga ccaagacaaa gcctttccgg agttcacgga
10681 gctgcgcaaa gtccaggcag acgcaatgca agacatggct cgcgccaact atgacgtttt
10741 gagctgctcg aacccgatct ttttcatgcg tgcgcgttac acgcgttaca tgcattccaa
10801 gtttccgggc ctgtatccgc cggatatggc cgagaaactg tactttacga gcgagccgta
10861 cgatcgtctg caacaaatcc agcgtaaaca gaatgtttgg tacaagattg gtcgcgtgaa
10921 ttgaggatcc tcgaccaggc atcaaataaa acgaaaggct cagtcgaaag actgggcctt
10981 tcgttttatc tgttgtttgt cggtgaacgc tctctactag agtcacactg gctcaccttc
11041 gggtgggcct ttctgcgttt atatgtttgc cctattttat ggctagctca gtcctaggta
11101 caatgctagc gtactttaac tttaagaagg agatatacat atgaagattc tggtcattgg
11161 tgctggtcca gctggtctgg ttttcgcatc ccaactgaag caggcacgcc ctttgtgggc
11221 cattgacatc gtggagaaga atgacgagca agaagtgctg ggctggggtg tcgtgctgcc
11281 tggccgtccg ggtcagcacc cggcgaaccc gctgtcctat ctggatgcac cggagcgtct

11341 gaatccgcaa tttctggagg acttcaaact ggtgcatcat aatgagccgt ccttgatgtc
11401 cacgggcgtt ttgttgtgcg gcgtggagcg tcgcggtctg gttcacgcgc tgcgcgataa
11461 gtgccgcagc caaggcattg ctattcgttt cgaaagcccg ttgctggaac acggtgagct
11521 gccgctggcg gactatgatc tggtggtcct ggctaatggt gttaatcaca aaaccgcgca
11581 tttcaccgag gctctggtcc cgcaggtgga ctacggccgc aataagtaca tttggtatgg
11641 cactagccag ctgttcgatc agatgaatct ggtttttcgt acccatggta aagatatctt
11701 tatcgcgcat gcctataagt atagcgatac catgagcacg ttcattgtcg aatgtagcga
11761 agagacttac gcacgcgcac gcctgggcga aatgtccgaa gaggcgagcg cagaatacgt
11821 tgctaaggtg ttccaggccg agctgggtgg tcacggcctg gtgagccagc cgggtctggg
11881 ttggcgtaac ttcatgacgt tgtctcatga ccgttgtcat gatggtaagt tggttctgct
11941 gggtgacgcg ctgcaaagcg gtcactttag catcggccac ggcaccacga tggccgtggt
12001 ggtggcgcag ctgctggtta aagcgctgtg taccgaagat ggtgtgcctg ccgcgctgaa
12061 acgtttcgaa gagcgtgccc tgccgctggt gcagttgttc cgtggccacg cagacaacag
12121 ccgcgtttgg ttcgaaaccg tcgaagagcg catgcacctg tcctcggcgg aatttgtgca
12181 aagcttcgac gcacgccgca aaagcctgcc gccgatgccg gaagcactgg cgcagaatct
12241 gcgttatgct ttgcagcgct gaggatcctc gaccaggcat caaataaaac gaaaggctca
12301 gtcgaaagac tgggcctttc gttttatctg ttgtttgtcg gtgaacgctc tctactagag
12361 tcacactggc tcaccttcgg gtgggccttt ctgcgtttat atgttccggc tattttatgg
12421 ctagctcagt cctaggtaca atgctagcgt actttaactt taagaaggag atatacatat
12481 ggagaaccgt gagccaccac tgttgccagc ccgttggagc agcgcctatg tctcttattg
12541 gagcccgatg ctgccggatg accagctgac cagcggctat tgctggttcg actatgaacg
12601 tgacatctgt cgtattgacg gcctgttcaa tccgtggagc gagcgtgata ctggttatcg
12661 cctgtggatg tcggaggttg gtaatgcggc cagcggccgt acctggaaac aaaaagtcgc
12721 ctatggtcgt gagcgtaccg ccctgggtga acagctgtgt gagcgtccgc tggatgatga
12781 gactggccct tttgccgaat tgttcctgcc acgcgatgtc ctgcgccgtc tgggtgcccg
12841 tcacattggc cgtcgcgtgg ttctgggtcg cgaagcggac ggttggcgtt accagcgccc
12901 aggtaaaggt ccgagcaccc tgtacctgga tgcggcgagc ggcactccac tgcgcatggt
12961 caccggcgat gaagcgtcgc gtgcaagcct gcgtgatttt ccgaatgtga gcgaggcgga
13021 gatcccggac gcggttttcg cggccaagcg ctaaggatcc tcgaccaggc atcaaataaa
13081 acgaaaggct cagtcgaaag actgggcctt tcgttttatc tgttgtttgt cggtgaacgc
13141 tctctactag agtcacactg gctcaccttc gggtgggcct ttctgcgttt atatgttgaa
13201 gctattttat ggctagctca gtcctaggta caatgctagc gtactttaac tttaagaagg
13261 agatatacat atgagattgc agcattacac gtcttgagcg attgtgtagg ctggagctgc
13321 ttcgaagttc ctatactttc tagagaatag gaacttcgga ataggaactt caagatcccc
13381 ttattagaag aactcgtcaa gaaggcgata gaaggcgatg cgctgcgaat cgggagcggc
13441 gataccgtaa agcacgagga agcggtcagc ccattcgccg ccaagctctt cagcaatatc
13501 acgggtagcc aacgctatgt cctgatagcg gtccgccaca cccagccggc cacagtcgat
13561 gaatccagaa aagcggccat tttccaccat gatattcggc aagcaggcat cgccatgggt
13621 cacgacgaga tcctcgccgt cgggcatgcg cgccttgagc ctggcgaaca gttcggctgg
13681 cgcgagcccc tgatgctctt cgtccagatc atcctgatcg acaagaccgg cttccatccg
13741 agtacgtgct cgctcgatgc gatgtttcgc ttggtggtcg aatgggcagg tagccggatc
13801 aagcgtatgc agccgccgca ttgcatcagc catgatggat actttctcgg caggagcaag
13861 gtgagatgac aggagatcct gccccggcac ttcgcccaat agcagccagt cccttcccgc
13921 ttcagtgaca acgtcgagca cagctgcgca aggaacgccc gtcgtggcca gccacgatag
13981 ccgcgctgcc tcgtcctgca gttcattcag ggcaccggac aggtcggtct tgacaaaaag
14041 aaccgggcgc ccctgcgctg acagccggaa cacggcggca tcagagcagc cgattgtctg
14101 ttgtgcccag tcatagccga atagcctctc cacccaagcg gccggagaac ctgcgtgcaa
14161 tccatcttgt tcaatcatgc gaaacgatcc tcatcctgtc tcttgatcag atcttgatcc
14221 cctgcgccat cagatccttg gcggcaagaa agccatccag tttactttgc agggcttccc
14281 aaccttacca gagggcgccc cagctggcaa ttccggttcg cttgctgtcc ataaaaccgc
14341 ccagtctagc tatcgccatg taagcccact gcaagctacc tgctttctct ttgcgcttgc
14401 gttttccctt gtccagatag cccagtagct gacattcatc cggggtcagc accgtttctg
14461 cggactggct ttctacgtgt tccgcttcct ttagcagccc ttgcgccctg agtgcttgcg
14521 gcagcgtgag cttcaaaagc gctctgaagt tcctatactt tctagagaat aggaacttcg
14581 aactgcaggt cgacggatcc tcgaccaggc atcaaataaa acgaaaggct cagtcgaaag
14641 actgggcctt tcgttttatc tgttgtttgt cggtgaacgc tctctactag agtcacactg
14701 gctcaccttc gggtgggcct ttctgcgttt atatgtttta gggacttaga gagacg
//

#### pTU2-A-RebOD-VioCDE

```
LOCUS       Exported File         10652 bp ds-DNA     circular SYN 22-SEP-2017
DEFINITION  .
ACCESSION   .
VERSION     .
KEYWORDS    pTU2-A-RebOD-VioEDC
SOURCE      synthetic DNA construct
  ORGANISM  synthetic DNA construct
REFERENCE   1  (bases 1 to 10652)
  AUTHORS   TERRENCE_LAI
  TITLE     Direct Submission
  JOURNAL   Exported 22 Sep 2017 from SnapGene 1.1.3
            http://www.snapgene.com
FEATURES             Location/Qualifiers
     source          1..10652
                     /organism="synthetic DNA construct"
                     /mol_type="other DNA"
     misc_feature    2066..2069
                     /note="Scar"
                     /note="color: #00ffff"
     CDS             2139..3572
                     /codon_start=1
                     /note="RebO7ClTrpoxidaseQ8K"
                     /note="color: #993366"
                     /translation="MSRGHKKITVLGAGVAGLVAAHELEELGHEVEVLEGSDRLGGRVH
                     THRFGEGGSVPFVELGAMRIPTKHRHTIDYIGKLGLTPKLKEFKTLFSDDGAYHTTSAG
                     FVRVRDAAKVLVDEFRLLMSGRDLREETILFGAWLTAVGDAIAPADFRAALRTDFTADL
                     LEVVDRIDLDPFLVGAARDQFDLHAFFAAHPEVRTSCTGKLNRFVDDILDETSPRLLRL
                     EGGMDQLVDALVERIRGDIRTGHEVSAIDVREDHVAVTVHNGHGVNTLRSDHVLCTIPF
                     SVLRNLRLTGLSTDKLEIIHDVKYWSATKVAFRCREPFWERDGINGGASFGGGRIRQTY

YPPVEGDPTRGAVLLASYTMGDDADVLGGMPEAQRHEVVLDEVGRMHPELHEPGMVVEA

VSRAWGEDRWSNGAGVTRWGKDVAACEEERDRAARPEGRLYFAGEHCSSTTAWIDGAVE
                     SALAAVRAIEAGDGRGSSS"
     misc_feature    3706..3709
                     /note="Scar"
                     /note="color: #00ffff"
     CDS             3779..6868
                     /codon_start=1
                     /note="RebDDichlorochromopy"
                     /note="color: #993366"
                     /translation="MSVFDLPRLHFAGTATTRLPTGPRNGLVDLSTHSVVMDGERFPAS

RPAAEYHAYLDRVGGKGTAFAGNGYFAIDAGITAVERAAGEVDTGDLLVGRAVDVWGHY

NEYLATTFNRARIFDVDPSSSWTSTVMIGQFGFGRLGRSHDVGYVFTGGVHGMQPPRWH

EDGRVLHQFTVPAGEDMTWFGSAADSPAAARLRELVESGEADGLVVQLALSDAGPAPMP
                     HAQQWRLRGTIAPWHAGEPRTCPAGRLLTPHNLTADLRGDHVSLNLISFRPPTGISGLE

LRTADTDRFIARVPADDPHGVVTVPAAEGGDEALCVVGTTAAGERIVVSREREVTVHVD
                     DASVFLEHPRGPGDSDQDAEIAVRTYVRGEPAAATIHIGQYFNPRAFPLDEHATAASAT

PEDLDVVALCVDGTRWSRHCVISTDENGDGRFLLRGARPGATRLLLSAEGATPFDGLTA

AAAYDNDDSLGLWSGLASVAVRVLPDHWWMDDIPRDKVTFDLLYREVFAFYELLYSFMG

EEVFSLADRFRVETHPRLIWQMCDPRNRAKTYYMPPTRDLTGPQARLLLAYLRAQNSDV
```

VVPVIEPSHTRSGTPISTRTDLVRALRHGVAIELAVMLQYLYAAFSIPTHGAGQELVSR

GDWTPEQLRLMCGDGGETTDGGVRGSLLGVAREEMIHFLVVNNVLMAVGEPFHVPDLDF
GTINDTLMVPLDFSLEALGLGSVQRFIQIEQPEGLTGAVRLGDLPVPVREAEDFHYASL
SELYGDIREGLQRVPGLFLVERGRGGGEHHLFLRESVNAVHPDYQLEVDDLSSALFAID

FVTEQGEGHVLTDEDTGEESHYDTFVRVADLLMKERLTAADTRRAQWSPAYPVARNPTV

HGGGQSKELVTSPVARELMVLFNKSYFMMLQLMVQHFGGSPDASLRRSKLMNAAIDVMT
GVMRPLAELLVTVPSGRHGRTAGPSFELDEKPAFIPRADVARRAISLRFRHLAESARTC
ALVPDKVVRNLDFLADQFATEGPRGSSSLVPRGSHHHHHH"
misc_feature 7002..7005
/note="Scar"
/note="color: #00ffff"
CDS 7075..7650
/codon_start=1
/note="VioE"
/note="color: #ffff00"
/translation="MENREPPLLPARWSSAYVSYWSPMLPDDQLTSGYCWFDYERDICR

IDGLFNPWSERDTGYRLWMSEVGNAASGRTWKQKVAYGRERTALGEQLCERPLDDETGP

FAELFLPRDVLRRLGARHIGRRVVLGREADGWRYQRPGKGPSTLYLDAASGTPLRMVTG
DEASRASLRDFPNVSEAEIPDAVFAAKR"
misc_feature 7794..7797
/note="Scar"
/note="color: #00ffff"
CDS 7867..8988
/codon_start=1
/note="VioD"
/note="color: #00ff00"
/translation="MKILVIGAGPAGLVFASQLKQARPLWAIDIVEKNDEQEVLGWGVV

LPGRPGQHPANPLSYLDAPERLNPQFLEDFKLVHHNEPSLMSTGVLLCGVERRGLVHAL
RDKCRSQGIAIRFESPLLEHGELPLADYDLVVLANGVNHKTAHFTEALVPQVDYGRNKY
IWYGTSQLFDQMNLVFRTHGKDIFIAHAYKYSDTMSTFIVECSEETYARARLGEMSEEA

SAEYVAKVFQAELGGHGLVSQPGLGWRNFMTLSHDRCHDGKLVLLGDALQSGHFSIGHG

TTMAVVVAQLLVKALCTEDGVPAALKRFEERALPLVQLFRGHADNSRVWFETVEERMHL
SSAEFVQSFDARRKSLPPMPEALAQNLRYALQR"
misc_feature 9132..9135
/note="Scar"
/note="color: #00ffff"
CDS 9205..10494
/codon_start=1
/note="VioC"
/note="color: #00ffff"
/translation="MKRAIIVGGGLAGGLTAIYLAKRGYEVHVVEKRGDPLRDLSSYVD
VVSSRAIGVSMTVRGIKSVLAAGIPRAELDACGEPIVAMAFSVGGQYRMRELKPLEDFR
PLSLNRAAFQKLLNKYANLAGVRYYFEHKCLDVDLDGKSVLIQGKDGQPQRLQGDMIIG

ADGAHSAVRQAMQSGLRRFEFQQTFFRHGYKTLVLPDAQALGYRKDTLYFFGMDSGGLF
AGRAATIPDGSVSIAVCLPYSGSPSLTTTDEPTMRAFFDRYFGGLPRDARDEMLRQFLA

KPSNDLINVRSSTFHYKGNVLLLGDAAHATAPFLGQGMNMALEDARTFVELLDRHQGDQ

DKAFPEFTELRKVQADAMQDMARANYDVLSCSNPIFFMRARYTRYMHSKFPGLYPPDMA
EKLYFTSEPYDRLQQIQRKQNVWYKIGRVN"
misc_feature 10638..10641

```
                     /note="Scar"
                     /note="color: #00ffff"
ORIGIN
        1 ctgcagtccg gcaaaaaagg gcaaggtgtc accaccctgc cctttttctt taaaaccgaa
       61 aagattactt cgcgttatgc aggcttcctc gctcactgac tcgctgcgct cggtcgttcg
      121 gctgcggcga gcggtatcag ctcactcaaa ggcggtaata cggttatcca cagaatcagg
      181 ggataacgca ggaaagaaca tgtgagcaaa aggccagcaa aaggccagga accgtaaaaa
      241 ggccgcgttg ctggcgtttt tccacaggct ccgcccccct gacgagcatc acaaaaatcg
      301 acgctcaagt cagaggtggc gaaacccgac aggactataa agataccagg cgtttccccc
      361 tggaagctcc ctcgtgcgct ctcctgttcc gaccctgccg cttaccggat acctgtccgc
      421 ctttctccct tcgggaagcg tggcgctttc tcatagctca cgctgtaggt atctcagttc
      481 ggtgtaggtc gttcgctcca agctgggctg tgtgcacgaa ccccccgttc agcccgaccg
      541 ctgcgcctta tccggtaact atcgtcttga gtccaacccg gtaagacacg acttatcgcc
      601 actggcagca gccactggta acaggattag cagagcgagg tatgtaggcg gtgctacaga
      661 gttcttgaag tggtggccta actacggcta cactagaaga acagtatttg gtatctgcgc
      721 tctgctgaag ccagttacct tcggaaaaag agttggtagc tcttgatccg gcaaacaaac
      781 caccgctggt agcggtggtt tttttgtttg caagcagcag attacgcgca gaaaaaaagg
      841 atctcaagaa gatcctttga tcttttctac ggggtctgac gctcagtgga acgaaaactc
      901 acgttaaggg attttggtca tgagattatc aaaaaggatc ttcacctaga tccttttaaa
      961 ttaaaaatga agttttaaat caatctaaag tatatatgag taaacttggt ctgacagctc
     1021 gaggcttgga ttctcaccaa taaaaaacgc ccggcggcaa ccgagcgttc tgaacaaatc
     1081 cagatggagt tctgaggtca ttactggatc tatcaacagg agtccaagcg agctcgatat
     1141 caaattacgc cccgccctgc cactcatcgc agtactgttg taattcatta agcattctgc
     1201 cgacatggaa gccatcacaa acggcatgat gaacctgaat cgccagcggc atcagcacct
     1261 tgtcgccttg cgtataatat ttgcccatgg tgaaaacggg ggcgaagaag ttgtccatat
     1321 tggccacgtt taaatcaaaa ctggtgaaac tcacccaggg attggctgac acgaaaaaca
     1381 tattctcaat aaacccttta gggaaatagg ccaggttttc accgtaacac gccacatctt
     1441 gcgaatatat gtgtagaaac tgccggaaat cgtcgtggta ttcactccag agcgatgaaa
     1501 acgtttcagt ttgctcatgg aaaacggtgt aacaagggtg aacactatcc catatcacca
     1561 gctcaccgtc tttcattgcc atacgaaatt ccggatgagc attcatcagg cgggcaagaa
     1621 tgtgaataaa ggccggataa aacttgtgct tatttttctt tacggtcttt aaaaaggccg
     1681 taatatccag ctgaacggtc tggttatagg tacattgagc aactgactga aatgcctcaa
     1741 aatgttcttt acgatgccat tgggatatat caacggtggt atatccagtg attttttct
     1801 ccattttagc ttccttagct cctgaaaatc tcgataactc aaaaaatacg cccggtagtg
     1861 atcttatttc attatggtga aagttggaac ctcttacgtg cccgatcaac tcgagtgcca
     1921 cctgacgtct aagaaaccat tattatcatg acattaacct ataaaaatag gcgtatcacg
     1981 aggcagaatt tcagataaaa aaaatcctta gctttcgcta aggatgattt ctggaattcg
     2041 cggccgcttc tagaggtctc actatatctc tattttatgg ctagctcagt cctaggtaca
     2101 atgctagcgt actttaactt taagaaggag atatacatat gagccgtggt cataaaaaga
     2161 ttaccgttct gggtgccggt gttgcaggtc tggttgcagc acatgaactg gaagaactgg
     2221 gtcatgaagt tgaagttctg gaaggtagcg atcgtctggg tggtcgtgtt catacccatc
     2281 gttttggtga aggtggtagc gttccgtttg ttgaactggg tgcaatgcgt attccgacca
     2341 aacatcgtca taccattgat tatattggta aactgggtct gaccccgaaa ctgaaagagt
     2401 ttaaaaccct gtttagtgat gatggtgcct atcataccac cagtgcaggt tttgttcgtg
     2461 ttcgtgatgc agcaaaagtt ctggttgatg aatttcgtct gctgatgagc ggtcgtgatc
     2521 tgcgtgaaga aaccattctg tttggtgcat ggctgaccgc agttggtgat gcaattgcac
     2581 cggcagattt tcgtgcagca ctgcgtaccg attttaccgc agatctgctg gaagttgttg
     2641 atcgtattga tctggacccg tttctggttg gtgcagcacg tgatcagttt gatctgcatg
     2701 cattttttgc agcacatccg gaagttcgta ccagctgtac cggtaaactg aatcgttttg
     2761 ttgatgatat cctggatgaa accagtccgc gtctgctgcg tctggaaggt ggtatggatc
     2821 agctggttga tgcactggtt gaacgtattc gtggtgatat tcgtaccggt catgaagtga
     2881 gcgcaattga tgttcgtgaa gatcatgttg cagttaccgt tcataatggt catggtgtta
     2941 ataccctgcg tagcgatcat gttctgtgta ccattccgtt tagcgttctg cgtaatctgc
     3001 gtctgaccgg tctgagcacc gataaactgg aaattattca cgacgtgaaa tattggagcg
     3061 caaccaaagt tgcatttcgt tgtcgtgaac cgttttggga acgtgatggt attaatggtg
     3121 gtgcaagctt tggtggtggt cgtattcgtc agacctatta tccgcctgtt gaaggtgatc
     3181 cgacccgtgg tgcagttctg ctggcaagct ataccatggg tgatgatgca gatgttctgg
     3241 gtggtatgcc ggaagcacag cgtcatgaag tggttctgga tgaagttggt cgtatgcatc
     3301 cggaactgca tgaaccgggt atggttgttg aagcagttag ccgtgcatgg ggtgaagatc
     3361 gttggagcaa tggtgcgggt gttacccgtt ggggtaaaga tgttgcagca tgtgaagaag
```

3421 aacgcgatcg cgcagcccgt ccggaaggtc gtctgtattt tgccggtgaa cattgtagca
3481 gcaccaccgc atggattgat ggtgcagttg aaagcgcact ggcagcagtt cgtgcaattg
3541 aagccggtga tggtcgtgga tcctcgagct aaccaggcat caaataaaac gaaaggctca
3601 gtcgaaagac tgggcctttc gttttatctg ttgtttgtcg gtgaacgctc tctactagag
3661 tcacactggc tcaccttcgg gtgggccttt ctgcgtttat atgtttgccc tattttatgg
3721 ctagctcagt cctaggtaca atgctagcgt actttaactt taagaaggag atatacatat
3781 gagcgttttt gatctgcctc gtctgcattt tgcaggcacc gcaaccaccc gtctgccgac
3841 cggtccgcgt aatggtctgg ttgatctgag cacccatagc gttgttatgg atggtgaacg
3901 ttttccggca agccgtccgg cagcagaata tcatgcatat ctggatcgtg ttggtggtaa
3961 aggcaccgca tttgcaggta atggttattt tgcaattgat gccggtatta ccgcagttga
4021 acgtgcagcc ggtgaagttg ataccggtga tctgctggtt ggtcgtgcag ttgatgtttg
4081 gggtcattat aacgaatatc tggccaccac ctttaatcgt gcacgtattt ttgatgttga
4141 tccgagcagc agctggacca gcaccgttat gattggtcag tttggttttg gtcgtctggg
4201 tcgtagccat gatgttggtt atgtttttac cggtggtgtt catggtatgc agcctccgcg
4261 ttggcatgaa gatggtcgtg ttctgcatca gtttaccgtt ccggcaggcg aagatatgac
4321 ctggtttggt agcgcagcag attcaccggc agcagcacgt ctgcgtgaac tggttgaaag
4381 cggtgaagca gatggtctgg tggttcagct ggcactgagt gatgcaggtc cggcaccgat
4441 gccgcatgca cagcagtggc gtctgcgtgg caccattgca ccgtggcatg ccggtgaacc
4501 gcgtacctgt ccggcaggtc gtctgctgac accgcataat ctgacagcag atctgcgtgg
4561 tgatcatgtt agcctgaatc tgattagctt tcgtccgcct accggtatta gcggtctgga
4621 actgcgtacc gcagataccg atcgttttat tgcacgtgtt ccggcagatg atccgcatgg
4681 tgttgttaca gttccggcag cggaaggtgg tgatgaagca ctgtgtgttg ttggcaccac
4741 cgcagcgggt gaacgtattg ttgttagccg tgaacgtgaa gttaccgttc atgttgatga
4801 tgcaagcgtt tttctggaac atccgcgtgg tccgggtgat agcgatcagg atgcagaaat
4861 tgcagttcgt acctatgttc gtggtgaacc ggcagccgca accattcata ttggtcagta
4921 tttcaatccg cgtgcatttc cgctggatga acatgcaacc gcagcaagcg caacaccgga
4981 agatctggat gttgttgccc tgtgtgttga tggtacacgt tggagccgtc attgtgttat
5041 tagcaccgat gaaaatggtg atggtcgctt tctgctgcgt ggtgcacgtc cgggtgcaac
5101 ccgtctgctg ctgagcgcag aaggtgcaac cccgtttgat ggtctgaccg cagcagcagc
5161 ctatgataat gatgatagcc tgggtctgtg gtcaggtctg gcaagcgttg cagttcgtgt
5221 tctgccggat cattggtgga tggatgatat tccgcgtgat aaagttacct tcgatctgct
5281 gtatcgtgaa gtgtttgcat tttatgaact gctgtatagc tttatgggcg aagaagtttt
5341 tagcctggca gatcgttttc gtgttgaaac ccatccgcgt ctgatttggc agatgtgtga
5401 tccgcgtaat cgtgcaaaaa cctattatat gcctccgacc cgtgatctga ccggtccgca
5461 ggcacgtctg ctgttagcat atctgcgtgc acagaatagt gatgttgttg ttccggttat
5521 tgaaccgagc catacccgta gcggcacccc gattagcacc cgtaccgatc tggttcgtgc
5581 actgcgtcat ggtgttgcaa ttgaactggc agttatgctg cagtatctgt atgcagcatt
5641 tagcattccg acccatggtg caggtcaaga actggttagc cgtggtgatt ggacaccgga
5701 acagctgcgt ctgatgtgtg gtgatggtgg tgaaaccacc gatggtggtg tgcgtggtag
5761 cctgctgggt gttgcacgtg aagaaatgat tcattttctg gtggtgaata atgttctgat
5821 ggcagttggt gaaccgtttc atgttccgga tctggatttt ggcaccatta atgatacccct
5881 gatggttccg ctggatttta gcctggaagc actgggtctg ggtagcgttc agcgttttat
5941 tcagattgaa cagccggaag gtctgaccgg tgcagttcgt ctgggtgatc tgccggttcc
6001 ggttcgtgaa gccgaagatt ttcattatgc aagcctgagc gaactgtatg gtgatattcg
6061 tgaaggtctg cagcgtgttc cgggtctgtt tctggttgaa cgtggtcgtg gtggtggtga
6121 acatcacctg tttctgcgtg aaagcgttaa tgcagttcat ccggattatc agctggaagt
6181 tgatgatctg agcagcgcac tgtttgccat tgattttgtt accgaacagg gtgaaggtca
6241 tgttctgacc gatgaagata ccggtgaaga aagccattat gataccttttg ttcgtgttgc
6301 cgatctgctg atgaaagaac gcctgaccgc agccgatacc cgtcgtgcac agtggtcacc
6361 ggcatatccg gttgcacgta atccgaccgt tcatggtggt ggtcagagca aagaactggt
6421 gaccagtccg gttgcccgtg aactgatggt tctgtttaac aaaagctact tcatgatgct
6481 gcagctgatg gttcagcatt ttggtggtag tccggatgca agcctgcgtc gtagcaaact
6541 gatgaatgca gcaattgatg ttatgaccgg tgttatgcgt ccgctggcag aactgctggt
6601 taccgttccg agcggtcgtc atggtcgtac cgcaggtccg tcatttgaac tggatgaaaa
6661 accggcattt attccgcgtg cagatgttgc acgtcgtgca attagcctgc gttttcgtca
6721 tctggcagaa agcgcacgta cctgtgcact ggttccggat aaagttgttc gtaatctgga
6781 ttttctggca gatcagtttg caaccgaagg tccgcgtgga tcctcgagcc tggtgccgcg
6841 cggcagccat catcatcatc atcactaacc aggcatcaaa taaaacgaaa ggctcagtcg
6901 aaagactggg cctttcgttt tatctgttgt ttgtcggtga acgctctcta ctagagtcac
6961 actggctcac cttcgggtgg gcctttctgc gtttatatgt tccggctatt ttatggctag

7021 ctcagtccta ggtacaatgc tagcgtactt taactttaag aaggagatat acatatggag
7081 aaccgtgagc caccactgtt gccagcccgt tggagcagcg cctatgtctc ttattggagc
7141 ccgatgctgc cggatgacca gctgaccagc ggctattgct ggttcgacta tgaacgtgac
7201 atctgtcgta ttgacggcct gttcaatccg tggagcgagc gtgatactgg ttatcgcctg
7261 tggatgtcgg aggttggtaa tgcggccagc ggccgtacct ggaaacaaaa agtcgcctat
7321 ggtcgtgagc gtaccgccct gggtgaacag ctgtgtgagc gtccgctgga tgatgagact
7381 ggcccttttg ccgaattgtt cctgccacgc gatgtcctgc gccgtctggg tgcccgtcac
7441 attggccgtc gcgtggttct gggtcgcgaa gcggacggtt ggcgttacca gcgcccaggt
7501 aaaggtccga gcaccctgta cctggatgcg gcgagcggca ctccactgcg catggtcacc
7561 ggcgatgaag cgtcgcgtgc aagcctgcgt gattttccga atgtgagcga ggcggagatc
7621 ccggacgcgg ttttcgcggc caagcgctaa ggatcctcga ccaggcatca aataaaacga
7681 aaggctcagt cgaaagactg ggcctttcgt tttatctgtt gtttgtcggt gaacgctctc
7741 tactagagtc acactggctc accttcgggt gggcctttct gcgtttatat gttgaagcta
7801 ttttatggct agctcagtcc taggtacaat gctagcgtac tttaacttta agaaggagat
7861 atacatatga agattctggt cattggtgct ggtccagctg gtctggtttt cgcatcccaa
7921 ctgaagcagg cacgcccttt gtgggccatt gacatcgtgg agaagaatga cgagcaagaa
7981 gtgctgggct ggggtgtcgt gctgcctggc cgtccgggtc agcacccggc gaacccgctg
8041 tcctatctgg atgcaccgga gcgtctgaat ccgcaatttc tggaggactt caaactggtg
8101 catcataatg agccgtcctt gatgtccacg ggcgttttgt tgtgcggcgt ggagcgtcgc
8161 ggtctggttc acgcgctgcg cgataagtgc cgcagccaag gcattgctat tcgtttcgaa
8221 agcccgttgc tggaacacgg tgagctgccg ctggcggact atgatctggt ggtcctggct
8281 aatggtgtta atcacaaaac cgcgcatttc accgaggctc tggtcccgca ggtggactac
8341 ggccgcaata agtacatttg gtatggcact agccagctgt tcgatcagat gaatctggtt
8401 tttcgtaccc atggtaaaga tatctttatc gcgcatgcct ataagtatag cgataccatg
8461 agcacgttca ttgtcgaatg tagcgaagag acttacgcac gcgcacgcct gggcgaaatg
8521 tccgaagagg cgagcgcaga atacgttgct aaggtgttcc aggccgagct gggtggtcac
8581 ggcctggtga gccagccggg tctgggttgg cgtaacttca tgacgttgtc tcatgaccgt
8641 tgtcatgatg gtaagttggt tctgctgggt gacgcgctgc aaagcggtca ctttagcatc
8701 ggccacggca ccacgatggc cgtggtggtg gcgcagctgc tggttaaagc gctgtgtacc
8761 gaagatggtg tgcctgccgc gctgaaacgt ttcgaagagc gtgccctgcc gctggtgcag
8821 ttgttccgtg gccacgcaga caacagccgc gtttggttcg aaaccgtcga agagcgcatg
8881 cacctgtcct cggcggaatt tgtgcaaagc ttcgacgcac gccgcaaaag cctgccgccg
8941 atgccggaag cactggcgca gaatctgcgt tatgctttgc agcgctgagg atcctcgacc
9001 aggcatcaaa taaaacgaaa ggctcagtcg aaagactggg cctttcgttt tatctgttgt
9061 ttgtcggtga acgctctcta ctagagtcac actggctcac cttcgggtgg gcctttctgc
9121 gtttatatgt tcttcctatt ttatggctag ctcagtccta ggtacaatgc tagcgtactt
9181 taactttaag aaggagatat acatatgaaa cgtgcgatta tcgttggtgg cggcctggcg
9241 ggtggcctga ccgcgatcta cctggcgaag cgtggctacg aagtgcacgt cgtggagaag
9301 cgtggtgatc ctctgcgcga tctgagctct tacgtggacg ttgttagcag ccgtgcgatc
9361 ggcgtgagca tgaccgttcg tggtatcaag agcgttttgg ctgcgggcat tccgcgtgca
9421 gagctggatg cgtgtggcga accgatcgtg gcaatggctt tctccgtggg tggtcagtat
9481 cgcatgcgcg aactgaagcc gttggaggat ttccgtccgc tgagcttgaa ccgtgcggcg
9541 tttcaaaagc tgctgaacaa atacgcgaac ctggcaggcg ttcgttacta ctttgagcat
9601 aagtgcctgg atgttgacct ggatggtaag agcgtgttga ttcagggcaa agatggtcag
9661 ccgcagcgtc tgcaaggtga catgattatc ggtgcggatg gcgcccacag cgccgtccgt
9721 caggcgatgc agagcggcct gcgtcgtttc gagttccagc aaacgttctt ccgccatggc
9781 tacaaaaccc tggttttgcc ggacgcgcaa gcactgggtt accgtaaaga cacgctgtac
9841 tttttcggca tggattccgg tggcctgttc gcgggtcgtg cggctacgat cccagatggt
9901 agcgtcagca tcgccgtttg cctgccgtac tcgggtagcc cttccctgac gaccaccgac
9961 gaaccgacga tgcgtgcgtt cttcgatcgt tacttcggtg gcctgccgcg tgacgcgcgt
10021 gacgaaatgc tgcgtcagtt tctggcgaag ccgagcaacg acctgattaa cgtgcgctct
10081 agcacctttc actataaggg taatgtgctg ttgctgggtg atgctgcgca tgcgactgcg
10141 ccgttcctgg gtcagggtat gaacatggcg ctggaggacg cccgcacgtt tgtcgagctg
10201 ctggaccgcc accagggcga ccaagacaaa gcctttccgg agttcacgga gctgcgcaaa
10261 gtccaggcag acgcaatgca agacatggct cgcgccaact atgacgtttt gagctgctcg
10321 aacccgatct tttcatgcg tgcgcgttac acgcgttaca tgcattccaa gtttccgggc
10381 ctgtatccgc cggatatggc cgagaaactg tactttacga gcgagccgta cgatcgtctg
10441 caacaaatcc agcgtaaaca gaatgtttgg tacaagattg gtcgcgtgaa ttgaggatcc
10501 tcgaccaggc atcaaataaa acgaaaggct cagtcgaaag actgggcctt tcgttttatc
10561 tgttgtttgt cggtgaacgc tctctactag agtcacactg gctcaccttc gggtgggcct

```
   10621 ttctgcgttt atatgtttta ggtacagaga cc
//
```

#### pTU2S-a-RebFH-VioABCDE

```
LOCUS       Exported File         13308 bp ds-DNA    circular SYN 22-SEP-2017
DEFINITION  .
ACCESSION   .
VERSION     .
KEYWORDS    pTU2S-a-RebFH-VioABCDE
SOURCE      synthetic DNA construct
  ORGANISM  synthetic DNA construct
REFERENCE   1  (bases 1 to 13308)
  AUTHORS   .
  TITLE     Direct Submission
  JOURNAL   Exported 22 Sep 2017 from SnapGene 1.1.3
            http://www.snapgene.com
COMMENT
FEATURES             Location/Qualifiers
     source          1..13308
                     /organism="synthetic DNA construct"
                     /mol_type="other DNA"
     misc_feature    1..6
                     /label="BBa Suffix"
                     /note="BBa Suffix"
                     /note="color: #ff00ff"
     misc_feature    146..165
                     /label="VR Primer"
                     /note="VR Primer"
                     /note="color: #00ffff"
     misc_feature    1922..1923
                     /label="VF2 Primer"
                     /note="VF2 Primer"
                     /note="color: #00ffff"
     misc_feature    1930..1933
                     /label="Scar(1)"
                     /note="Scar"
                     /note="color: #80ff00"
     primer_bind     1952..1971
                     /label="T7"
                     /note="T7"
                     /note="color: #00ffff; direction: RIGHT"
     misc_feature    1952..1968
                     /label="T7 Promoter"
                     /note="T7 Promoter"
                     /note="color: #00ffff"
     misc_binding    1971..1993
                     /label="LacO"
                     /note="LacO"
                     /note="color: #6495ed"
     misc_feature    1998..2003
                     /label="BBa Prefix(1)"
                     /note="BBa Prefix"
                     /note="color: #ff00ff"
     misc_feature    2051..2068
                     /label="His-Tag(4)"
                     /label="His-Tag(4)"
                     /note="His-Tag"
                     /note="color: #00ffff"
```

```
misc_feature    2078..2095
                /label="Thrombin cleavage(4)"
                /note="Thrombin cleavage"
                /note="color: #ff8080"
misc_feature    2102..3355
                /label="VioA"
                /note="VioA"
                /note="color: #00ffff"
primer_bind     3517..3536
                /label="T7(1)"
                /note="T7"
                /note="color: #00ffff; direction: RIGHT"
misc_feature    3517..3533
                /label="T7 Promoter(1)"
                /note="T7 Promoter"
                /note="color: #00ffff"
misc_binding    3536..3558
                /label="LacO(1)"
                /note="LacO"
                /note="color: #6495ed"
misc_feature    3563..3568
                /label="BBa Prefix(2)"
                /note="BBa Prefix"
                /note="color: #ff00ff"
misc_feature    3616..3633
                /label="His-Tag(1)"
                /note="His-Tag"
                /note="color: #00ffff"
misc_feature    3643..3660
                /label="Thrombin cleavage(1)"
                /note="Thrombin cleavage"
                /note="color: #ff8080"
misc_feature    3667..6660
                /label="VioB"
                /note="VioB"
                /note="color: #00ff80"
misc_feature    4317..4336
                /label="VioBSeqF"
                /note="VioBSeqF"
                /note="color: #00ffff"
misc_feature    5994..6015
                /label="VioBSeqR"
                /note="VioBSeqR"
                /note="color: #00ffff"
primer_bind     6822..6841
                /label="T7(2)"
                /note="T7"
                /note="color: #00ffff; direction: RIGHT"
misc_feature    6822..6838
                /label="T7 Promoter(4)"
                /note="T7 Promoter"
                /note="color: #00ffff"
misc_binding    6841..6863
                /label="LacO(2)"
                /note="LacO"
                /note="color: #6495ed"
misc_feature    6868..6873
                /label="BBa Prefix(3)"
                /note="BBa Prefix"
                /note="color: #ff00ff"
```

```
misc_feature    6921..6938
                /label="His-Tag"
                /note="His-Tag"
                /note="color: #00ffff"
misc_feature    6948..6965
                /label="Thrombin cleavage"
                /note="Thrombin cleavage"
                /note="color: #ff8080"
misc_feature    6972..7544
                /label="VioE"
                /note="VioE"
                /note="color: #8080ff"
primer_bind     7706..7725
                /label="T7(3)"
                /note="T7"
                /note="color: #00ffff; direction: RIGHT"
misc_feature    7706..7722
                /label="T7 Promoter(3)"
                /note="T7 Promoter"
                /note="color: #00ffff"
misc_binding    7725..7747
                /label="LacO(3)"
                /note="LacO"
                /note="color: #6495ed"
misc_feature    7752..7757
                /label="BBa Prefix(4)"
                /note="BBa Prefix"
                /note="color: #ff00ff"
misc_feature    7805..7822
                /label="His-Tag(2)"
                /note="His-Tag"
                /note="color: #00ffff"
misc_feature    7832..7849
                /label="Thrombin cleavage(3)"
                /note="Thrombin cleavage"
                /note="color: #ff8080"
misc_feature    7856..8974
                /label="VioD"
                /note="VioD"
                /note="color: #ffff80"
primer_bind     9136..9155
                /label="T7(4)"
                /note="T7"
                /note="color: #00ffff; direction: RIGHT"
misc_feature    9136..9152
                /label="T7 Promoter(2)"
                /note="T7 Promoter"
                /note="color: #00ffff"
misc_binding    9155..9177
                /label="LacO(4)"
                /note="LacO"
                /note="color: #6495ed"
misc_feature    9182..9187
                /label="BBa Prefix(5)"
                /note="BBa Prefix"
                /note="color: #ff00ff"
misc_feature    9235..9252
                /label="His-Tag(3)"
                /note="His-Tag"
                /note="color: #00ffff"
```

```
     misc_feature    9262..9279
                     /label="Thrombin cleavage(2)"
                     /note="Thrombin cleavage"
                     /note="color: #ff8080"
     misc_feature    9286..10572
                     /label="VioC"
                     /note="VioC"
                     /note="color: #ff80c0"
     misc_feature    10739..10744
                     /label="BBa Prefix"
                     /note="BBa Prefix"
                     /note="color: #ff00ff"
     misc_feature    10745..10750
                     /label="BsaI"
                     /note="BsaI"
                     /note="color: #ff0000"
     misc_feature    10752..10755
                     /label="Scar"
                     /note="Scar"
                     /note="color: #80ff00"
     CDS             10832..11341
                     /label="RebF_reductase"
                     /note="RebF_reductase"
                     /note="color: #ffc0cb; direction: RIGHT"
     CDS             11561..13150
                     /label="halogenase_RebH"
                     /note="halogenase_RebH"
                     /note="color: #ffc0cb; direction: RIGHT"
     misc_feature    11639..11644
                     /label="BBa Suffix(1)"
                     /note="BBa Suffix"
                     /note="color: #ff00ff"
     misc_feature    11666..11671
                     /label="BBa Suffix(2)"
                     /note="BBa Suffix"
                     /note="color: #ff00ff"
     misc_feature    11720..11725
                     /label="BBa Suffix(3)"
                     /note="BBa Suffix"
                     /note="color: #ff00ff"
     misc_feature    complement(13303..13308)
                     /label="BsaI(1)"
                     /note="BsaI"
                     /note="color: #00ff00; direction: LEFT"
ORIGIN
        1 ctgcagtccg gcaaaaaagg gcaaggtgtc accaccctgc ccttttttctt taaaaccgaa
       61 aagattactt cgcgttatgc aggcttcctc gctcactgac tcgctgcgct cggtcgttcg
      121 gctgcggcga gcggtatcag ctcactcaaa ggcggtaata cggttatcca cagaatcagg
      181 ggataacgca ggaaagaaca tgtgagcaaa aggccagcaa aaggccagga accgtaaaaa
      241 ggccgcgttg ctggcgtttt tccacaggct ccgcccccct gacgagcatc acaaaaatcg
      301 acgctcaagt cagaggtggc gaaacccgac aggactataa agataccagg cgtttccccc
      361 tggaagctcc ctcgtgcgct ctcctgttcc gaccctgccg cttaccggat acctgtccgc
      421 ctttctccct tcgggaagcg tggcgctttc tcatagctca cgctgtaggt atctcagttc
      481 ggtgtaggtc gttcgctcca agctgggctg tgtgcacgaa ccccccgttc agcccgaccg
      541 ctgcgcctta tccggtaact atcgtcttga gtccaacccg gtaagacacg acttatcgcc
      601 actggcagca gccactggta acaggattag cagagcgagg tatgtaggcg gtgctacaga
      661 gttcttgaag tggtggccta actacggcta cactagaaga acagtatttg gtatctgcgc
      721 tctgctgaag ccagttacct tcggaaaaag agttggtagc tcttgatccg gcaaacaaac
      781 caccgctggt agcggtggtt tttttgtttg caagcagcag attacgcgca gaaaaaaagg
      841 atctcaagaa gatcctttga tcttttctac ggggtctgac gctcagtgga acgaaaactc
```

901 acgttaaggg attttggtca tgagattatc aaaaaggatc ttcacctaga tccttttaaa
961 ttaaaaatga agttttaaat caatctaaag tatatatgag taaacttggt ctgacagctc
1021 gaggcttgga ttctcaccaa taaaaaacgc ccggcggcaa ccgagcgttc tgaacaaatc
1081 cagatggagt tctgaggtca ttactggatc tatcaacagg agtccaagcg agctcgatat
1141 caaattacgc cccgccctgc cactcatcgc agtactgttg taattcatta agcattctgc
1201 cgacatggaa gccatcacaa acggcatgat gaacctgaat cgccagcggc atcagcacct
1261 tgtcgccttg cgtataatat ttgcccatgg tgaaaacggg ggcgaagaag ttgtccatat
1321 tggccacgtt taaatcaaaa ctggtgaaac tcacccaggg attggctgac acgaaaaaca
1381 tattctcaat aaaccettta gggaaatagg ccaggttttc accgtaacac gccacatctt
1441 gcgaatatat gtgtagaaac tgccggaaat cgtcgtggta ttcactccag agcgatgaaa
1501 acgtttcagt ttgctcatgg aaaacggtgt aacaagggtg aacactatcc catatcacca
1561 gctcaccgtc tttcattgcc atacgaaatt ccggatgagc attcatcagg cgggcaagaa
1621 tgtgaataaa ggccggataa aacttgtgct tatttttctt tacggtcttt aaaaaggccg
1681 taatatccag ctgaacggtc tggttatagg tacattgagc aactgactga aatgcctcaa
1741 aatgttcttt acgatgccat tgggatatat caacggtggt atatccagtg attttttct
1801 ccattttagc ttccttagct cctgaaaatc tcgataactc aaaaaatacg cccggtagtg
1861 atcttatttc attatggtga aagttggaac ctcttacgtg cccgatcaac tcgagtgcca
1921 cctgacgtcc tatatctcta tcccgcgaaa ttaatacgac tcactatagg ggaattgtga
1981 gcggataaca attcccctct agaaataatt ttgtttaact ttaagaagga gatataccat
2041 gggcagcagc catcatcatc atcatcacag cagcggcctg gtgccgcgcg gcagccatat
2101 gaaacattct tccgatatct gcattgttgg tgctggtatt tctggtttga cgtgcgcaag
2161 ccatctgctg gacagcccgg catgccgtgg tctgagcctg cgtatctttg acatgcagca
2221 agaagccggt ggccgtatcc gcagcaaaat gctggatggt aaggcaagca ttgaactggg
2281 cgcaggtcgc tactcccctc agttgcaccc gcatttccaa agcgcaatgc agcactatag
2341 ccaaaagagc gaagtctatc cgttcaccca gttgaagttc aaatctcacg tgcagcaaaa
2401 gctgaagcgc gccatgaatg aactgtcccc gcgtctgaaa gagcatggta aagagagctt
2461 tttgcagttt gtcagccgtt atcaaggtca cgatagcgcg gttggtatga tccgctctat
2521 gggttacgac gcactgttcc tgccggatat cagcgcagaa atggcctacg acattgtggg
2581 taagcacccg gagatccaga gcgtgacgga caacgacgcg aaccaatggt ttgcagcgga
2641 aacgggcttt gctggtctga ttcagggcat caaggctaag gttaaggcgg caggtgcgcg
2701 ttttagcctg ggttatcgtc tgctgagcgt ccgtaccgac ggtgacggct acctgctgca
2761 actggcaggt gacgacggct ggaaactgga gcaccgtacc cgccatctga ttctggcgat
2821 tccgccgagc gcgatggcgg gtttgaatgt tgattttcca gaagcctggt ccggtgcgcg
2881 ctatggcagc ctgccgctgt ttaagggctt tctgacgtac ggtgagccgt ggtggttgga
2941 ctacaaactg gacgatcagg tgctgattgt tgacaacccg ctgcgcaaaa tctatttcaa
3001 aggcgataag tacctgttct tctataccga tagcgagatg gcgaattact ggcgcggttg
3061 tgtcgcggag ggcgaggacg gttacctgga gcaaattcgc acccatttgg ctagcgcact
3121 gggtatcgtc cgtgaacgta tcccgcaacc gctggcacac gttcacaagt attgggcgca
3181 cggcgttgag ttttgccgtg attctgatat tgaccacccg agcgcactgt ctcatcgcga
3241 cagcggtatc atcgcgtgct ccgatgcgta cacggagcat tgtggttgga tggagggcgg
3301 tctgctgagc gcccgtgagg caagccgtct gctgttgcag cgtatcgccg cgtgaggatc
3361 ctcgaccagg catcaaataa aacgaaaggc tcagtcgaaa gactgggcct ttcgttttat
3421 ctgttgtttg tcggtgaacg ctctctacta gagtcacact ggctcacctt cgggtgggcc
3481 tttctgcgtt tatatgtttg ccctatcccg cgaaattaat acgactcact ataggggaat
3541 tgtgagcgga taacaattcc cctctagaaa taattttgtt taactttaag aaggagatat
3601 accatgggca gcagccatca tcatcatcat cacagcagcg gcctggtgcc gcgcggcagc
3661 catatgagca ttctggattt cccgcgtatc cacttccgtg gctgggcccg tgtcaatgcg
3721 ccgaccgcga accgcgatcc gcacggccac atcgatatgg ccagcaatac cgtggcgatg
3781 gcgggtgagc cgttcgacct ggcacgccat cctacggagt tccaccgtca cctgcgctcc
3841 ctgggtccgc gcttcggctt ggatggtcgt gctgacccgg aaggcccgtt cagcctggcc
3901 gagggctaca acgctgccgg taacaaccac ttttcgtggg agagcgcaac cgttagccac
3961 gtgcaatggg atggcggtga ggcggatcgt ggtgacggtc tggtcggtgc tcgtttggca
4021 ctgtggggtc actacaatga ttatctgcgt accaccttca atcgtgctcg ttgggtcgac
4081 agcgacccga cgcgccgtga cgctgcacaa atctatgcgg gccaattcac cattagcccg
4141 gctggtgccg gtccgggtac gccgtggctg tttacggcag acattgatga tagccatggt
4201 gcacgttgga cgcgtggcgg ccacattgca gagcgtggcg gccacttctt ggatgaagag
4261 tttggtctgg cacgcctgtt tcagttctct gtgccgaaag atcacccaca ttttctgttt
4321 cacccgggtc cgtttgattc cgaggcctgg cgtcgtctgc aattggctct ggaggatgac
4381 gacgttctgg gtctgaccgt gcaaatatgcg ttgttcaata tgagcacccc gcctcagccg
4441 aacagcccgg tttttcacga tatggtcggt gttgtcggtc tgtggcgtcg tggtgaactg

4501 gcgagctacc cggctggtcg tctgctgcgt ccgcgtcaac cgggtctggg tgacctgacc
4561 ctgcgcgtca acggtggtcg cgttgcgctg aatttggcgt gtgccattcc gttcagcact
4621 cgtgccgcgc agccaagcgc accggaccgc ctgaccccgg acctgggtgc caaactgccg
4681 ctgggcgatc tgctgctgcg tgatgaggac ggcgcactgt tggcacgtgt gccgcaggct
4741 ctgtaccaag actattggac gaatcacggt attgtggacc tgccgctgct gcgcgaaccg
4801 cgtggtagct tgaccctgag cagcgaactg gcggagtggc gtgagcaaga ctgggtcacc
4861 caaagcgacg cgtctaacct gtacctggag gcaccggatc gccgtcacgg tcgctttttc
4921 cctgagagca tcgcgctgcg cagctacttt cgcggtgaag cgcgtgcgcg tccggatatc
4981 ccgcatcgta tcgagggcat gggcctggtc ggcgtcgaat ctcgtcagga tggcgacgct
5041 gcggaatggc gtctgacggg tctgcgtccg ggtccggcac gcattgttct ggacgatggt
5101 gccgaggcga tccctctgcg tgttctgcct gacgattggg cgctggatga cgcgaccgtc
5161 gaagaagtgg attacgcctt tttgtaccgc cacgttatgg cgtattacga gctggtgtat
5221 ccattcatga gcgacaaggt gttttccctg gctgatcgtt gcaaatgtga aacgtacgca
5281 cgtctgatgt ggcagatgtg tgatccgcag aaccgcaaca agtcctatta catgccgagc
5341 acccgcgaac tgtcggcacc gaaagctcgt ttgttcttga agtatctggc ccacgtggaa
5401 ggccaggcac gcctgcaagc acctccgcca gcgggtccgg cacgcattga atctaaagcc
5461 cagttggcgg cagagctgcg taaagccgtc gacctggagc tgtctgtgat gctgcaatac
5521 ctgtacgcgg cgtatagcat tccgaactat gcacagggcc aacaacgtgt tcgtgacggt
5581 gcgtggaccg ccgagcagct gcaactggcg tgcggtagcg gtgaccgtcg ccgtgatggc
5641 ggtattcgtg cagcactgct ggaaattgct catgaagaaa tgattcatta cctggtcgtt
5701 aacaacctgc tgatggccct gggcgagccg ttctacgcgg gtgtcccgct gatgggcgaa
5761 gcggcacgtc aggcgtttgg cctggacacc gagttcgctc tggaaccgtt tagcgaaagc
5821 acgctggcac gttttgttcg tctggaatgg ccgcacttta tcccagcacc gggcaaatcc
5881 atcgcggact gctatgccgc cattcgtcag gcgtttttgg atctgccgga cttgtttggt
5941 ggcgaggcag gtaagcgtgg cggtgaacac cacctgttcc tgaatgagct gaccaaccgt
6001 gcgcatccgg gttatcaact ggaagttttc gatcgcgact cggcgctgtt tggtattgca
6061 tttgtgaccg atcagggcga aggtggcgct ctggacagcc cgcactacga acatagccat
6121 tttcaacgtc tgcgtgaaat gagcgcgcgt atcatggctc aaagcgcacc gttcgaaccg
6181 gcgctgccgg cgttgcgtaa tccggttctg gatgagagcc cggttgccaa acgtgtcgca
6241 gacggtcgtg cgcgtgcgct gatggcattg taccaaggcg tttatgagct gatgtttgcg
6301 atgatggcgc agcacttcgc cgtgaaaccg ctgggtagct tgcgtcgcag ccgcctgatg
6361 aacgcagcaa tcgatctgat gaccggtctg ttgcgtccgc tgagctgcgc gctgatgaac
6421 ctgccaagcg gcatcgccgg tcgcacggcc ggtccgccgc tgccgggtcc ggttgacacc
6481 cgtagctatg acgactacgc gctgggctgt cgcatgctgg cacgccgttg cgagcgtctg
6541 ctggagcagg cgagcatgct ggaaccgggt tggctgccgg atgcgcagat ggagctgctg
6601 gatttctatc gtcgccaaat gctggacttg gcgtgcggca aactgagccg cgaggcctaa
6661 ggatcctcga ccaggcatca aataaaacga aaggctcagt cgaaagactg ggcctttcgt
6721 tttatctgtt gtttgtcggt gaacgctctc tactagagtc acactggctc accttcgggt
6781 gggcctttct gcgtttatat gttccggcta tcccgcgaaa ttaatacgac tcactatagg
6841 ggaattgtga gcggataaca attcccctct agaaataatt ttgtttaact ttaagaagga
6901 gatataccat gggcagcagc catcatcatc atcatcacag cagcggcctg gtgccgcgcg
6961 gcagccatat ggagaaccgt gagccaccac tgttgccagc ccgttggagc agcgcctatg
7021 tctcttattg gagcccgatg ctgccggatg accagctgac cagcggctat tgctggttcg
7081 actatgaacg tgacatctgt cgtattgacg gcctgttcaa tccgtggagc gagcgtgata
7141 ctggttatcg cctgtggatg tcggaggttg gtaatgcggc cagcggccgt acctggaaac
7201 aaaaagtcgc ctatggtcgt gagcgtaccg ccctgggtga acagctgtgt gagcgtccgc
7261 tggatgatga gactggccct tttgccgaat tgttcctgcc acgcgatgtc ctgcgccgtc
7321 tgggtgcccg tcacattggc cgtcgcgtgg ttctgggtcg cgaagcggac ggttggcgtt
7381 accagcgccc aggtaaaggt ccgagcaccc tgtacctgga tgcggcgagc ggcactccac
7441 tgcgcatggt caccggcgat gaagcgtcgc gtgcaagcct gcgtgatttt ccgaatgtga
7501 gcgaggcgga gatcccggac gcggttttcg cggccaagcg ctaaggatcc tcgaccaggc
7561 atcaaataaa acgaaaggct cagtcgaaag actgggcctt tcgttttatc tgttgtttgt
7621 cggtgaacgc tctctactag agtcacactg gctcaccttc gggtgggcct ttctgcgttt
7681 atatgttgaa gctatcccgc gaaattaata cgactcacta taggggaatt gtgagcggat
7741 aacaattccc ctctagaaat aattttgttt aactttaaga aggagatata ccatgggcag
7801 cagccatcat catcatcatc acagcagcgg cctggtgccg cgcggcagcc atatgaagat
7861 tctggtcatt ggtgctggtc cagctggtct ggttttcgca tcccaactga agcaggcacg
7921 ccctttgtgg gccattgaca tcgtggagaa gaatgacgag caagaagtgc tgggctgggg
7981 tgtcgtgctg cctggccgtc cgggtcagca cccggcgaac ccgctgtcct atctggatgc
8041 accggagcgt ctgaatccgc aatttctgga ggacttcaaa ctggtgcatc ataatgagcc

8101 gtccttgatg tccacgggcg ttttgttgtg cggcgtggag cgtcgcggtc tggttcacgc
8161 gctgcgcgat aagtgccgca gccaaggcat tgctattcgt ttcgaaagcc cgttgctgga
8221 acacggtgag ctgccgctgg cggactatga tctggtggtc ctggctaatg gtgttaatca
8281 caaaaccgcg catttcaccg aggctctggt cccgcaggtg gactacggcc gcaataagta
8341 catttggtat ggcactagcc agctgttcga tcagatgaat ctggtttttc gtacccatgg
8401 taaagatatc tttatcgcgc atgcctataa gtatagcgat accatgagca cgttcattgt
8461 cgaatgtagc gaagagactt acgcacgcgc acgcctgggc gaaatgtccg aagaggcgag
8521 cgcagaatac gttgctaagg tgttccaggc cgagctgggt ggtcacggcc tggtgagcca
8581 gccgggtctg ggttggcgta acttcatgac gttgtctcat gaccgttgtc atgatggtaa
8641 gttggttctg ctgggtgacg cgctgcaaag cggtcacttt agcatcggcc acggcaccac
8701 gatggccgtg gtggtggcgc agctgctggt taaagcgctg tgtaccgaag atggtgtgcc
8761 tgccgcgctg aaacgtttcg aagagcgtgc cctgccgctg gtgcagttgt tccgtggcca
8821 cgcagacaac agccgcgttt ggttcgaaac cgtcgaagag cgcatgcacc tgtcctcggc
8881 ggaatttgtg caaagcttcg acgcacgccg caaaagcctg ccgccgatgc cggaagcact
8941 ggcgcagaat ctgcgttatg ctttgcagcg ctgaggatcc tcgaccaggc atcaaataaa
9001 acgaaaggct cagtcgaaag actgggcctt tcgttttatc tgttgtttgt cggtgaacgc
9061 tctctactag agtcacactg gctcaccttc gggtgggcct ttctgcgttt atatgttctt
9121 cctatcccgc gaaattaata cgactcacta taggggaatt gtgagcggat aacaattccc
9181 ctctagaaat aattttgttt aactttaaga aggagatata ccatgggcag cagccatcat
9241 catcatcatc acagcagcgg cctggtgccg cgcggcagcc atatgaaacg tgcgattatc
9301 gttggtggcg gcctggcggg tggcctgacc gcgatctacc tggcgaagcg tggctacgaa
9361 gtgcacgtcg tggagaagcg tggtgatcct ctgcgcgatc tgagctctta cgtggacgtt
9421 gttagcagcc gtgcgatcgg cgtgagcatg accgttcgtg gtatcaagag cgttttggct
9481 gcgggcattc cgcgtgcaga gctggatgcg tgtggcgaac cgatcgtggc aatggctttc
9541 tccgtgggtg gtcagtatcg catgcgcgaa ctgaagccgt tggaggattt ccgtccgctg
9601 agcttgaacc gtgcggcgtt tcaaaagctg ctgaacaaat acgcgaacct ggcaggcgtt
9661 cgttactact ttgagcataa gtgcctggat gttgacctgg atggtaagag cgtgttgatt
9721 cagggcaaag atggtcagcc gcagcgtctg caaggtgaca tgattatcgg tgcggatggc
9781 gcccacagcg ccgtccgtca ggcgatgcag agcggcctgc gtcgtttcga gttccagcaa
9841 acgttcttcc gccatggcta caaaaccctg gttttgccgg acgcgcaagc actgggttac
9901 cgtaaagaca cgctgtactt tttcggcatg gattccggtg gcctgttcgc gggtcgtgcg
9961 gctacgatcc cagatggtag cgtcagcatc gccgtttgcc tgccgtactc gggtagccct
10021 tccctgacga ccaccgacga accgacgatg cgtgcgttct tcgatcgtta cttcggtggc
10081 ctgccgcgtg acgcgcgtga cgaaatgctg cgtcagtttc tggcgaagcc gagcaacgac
10141 ctgattaacg tgcgctctag cacctttcac tataagggta atgtgctgtt gctgggtgat
10201 gctgcgcatg cgactgcgcc gttcctgggt cagggtatga acatggcgct ggaggacgcc
10261 cgcacgtttg tcgagctgct ggaccgccac cagggcgacc aagacaaagc ctttccggag
10321 ttcacggagc tgcgcaaagt ccaggcagac gcaatgcaag acatggctcg cgccaactat
10381 gacgttttga gctgctcgaa cccgatcttt ttcatgcgtg cgcgttacac gcgttacatg
10441 cattccaagt ttccgggcct gtatccgccg gatatggccg agaaactgta ctttacgagc
10501 gagccgtacg atcgtctgca acaaatccag cgtaaacaga atgtttggta caagattggt
10561 cgcgtgaatt gaggatcctc gaccaggcat caaataaaac gaaaggctca gtcgaaagac
10621 tgggcctttc gttttatctg ttgtttgtcg gtgaacgctc tctactagag tcacactggc
10681 tcaccttcgg gtgggccttt ctgcgtttat atgttttagg tacgaattcg cggccgcttc
10741 tagaggtctc actatatctc tattttatgg ctagctcagt cctaggtaca atgctagcgt
10801 actttaactt taagaaggag atatacatat gaccatcgaa tttgatcgtc cgggtgcaca
10861 tgttaccgca gcagatcatc gtgccctgat gagcctgttt ccgaccggtg ttgcagttat
10921 taccgcaatt gatgaagcag gtacaccgca tggtatgacc tgtaccagcc tgaccagcgt
10981 taccctggac cctccgaccc tgctggtttg tctgaatcgt gcaagcggca ccctgcatgc
11041 cgttcgtggt ggtcgttttg gtgttaatct gctgcatgca cgtggtcgtc gtgcagcaga
11101 agtttttagc accgcagttc aggatcgctt tggtgaagtt cgttgggaac atagtgatgt
11161 taccggtatg ccgtggctgg ccgaagatgc acatgcattt gcaggttgtg ttgttcgtaa
11221 aagcaccgtt gttggtgatc atgaaattgt tctgggtgaa gtgcatgaag ttgttcgtga
11281 acatgatctg ccgctgctgt atggtatgcg tgaatttgca gtttggacac cggaaggtta
11341 aggatcctcg accaggcatc aaataaaacg aaaggctcag tcgaaagact gggcctttcg
11401 ttttatctgt tgtttgtcgg tgaacgctct ctactagagt cacactggct caccttcggg
11461 tgggcctttc tgcgtttata tgtttgccct attttatggc tagctcagtc ctaggtacaa
11521 tgctagcgta ctttaacttt aagaaggaga tatacatatg agcggcaaaa tcgacaaaat
11581 tctgattgtt ggtggtggca ccgcaggttg gatggcagca agctatctgg gtaaagcact
11641 gcagggtaca gcagatatta ccctgctgca ggcaccggat attccgaccc tgggtgttgg

11701 tgaagcaacc attccgaatc tgcagaccgc attttttgat tttctgggta ttccggaaga
11761 tgaatggatg cgtgaatgta atgcaagcta taaagtggcc atcaaattca ttaattggcg
11821 taccgcaggc gaaggcacca gcgaagcacg tgaactggat ggtggtccgg atcattttta
11881 tcatagcttt ggtctgctga aataccatga gcagattccg ctgagccatt attggtttga
11941 tcgtagctat cgtggtaaaa ccgttgaacc gtttgattac gcctgttata aagaaccggt
12001 tattctggat gcaaatcgta gtccgcgtcg tctggatggt agcaaagtta ccaattatgc
12061 atggcatttt gatgcacatc tggttgcaga ttttctgcgt cgttttgcaa ccgaaaaact
12121 gggtgttcgt catgttgaag atcgtgttga acatgtgcag cgtgatgcaa atggtaatat
12181 tgaaagcgtt cgtaccgcaa ccggtcgtgt tttgatgcc gacctgtttg ttgattgtag
12241 cggttttcgt ggcctgctga ttaacaaagc aatggaagaa ccgtttctgg atatgagcga
12301 tcatctgctg aatgatagcg cagttgcaac ccaggttccg catgatgatg atgccaatgg
12361 tgtggaaccg tttaccagcg caattgcaat gaaaagcggt tggacctgga aaattccgat
12421 gctgggtcgt tttggcaccg gttatgttta tagcagccgc tttgccaccg aagatgaagc
12481 agttcgtgaa ttttgtgaaa tgtggcatct ggacccggaa acccagccgc tgaatcgtat
12541 tcgttttcgt gttggtcgta atcgtcgtgc atgggttggt aattgtgtta gcattggcac
12601 cagcagctgt tttgttgaac cgctggaaag caccggtatc tattttgttt atgcagcact
12661 gtatcagctg gtgaaacatt ttccggataa aagcctgaat ccggttctga ccgcacgttt
12721 taatcgtgaa attgaaacca tgttcgatga cacccgtgat tttattcagg cccactttta
12781 tttcagtccg cgtaccgata ccccgttttg gcgtgcaaat aaagaactgc gtctggcaga
12841 tggtatgcaa gaaaaaattg atatgtatcg tgccggtatg gcaattaatg caccggcaag
12901 tgatgatgca cagctgtatt atggcaactt tgaagaagaa tttcgcaact tctggaacaa
12961 cagcaactat tattgtgttc tggcaggtct gggtctggtt ccggatgcac cgtcaccgcg
13021 tctggcccac atgccgcagg caaccgaatc agttgatgaa gtttttggtg cagttaaaga
13081 tcgtcagcgt aacctgctgg aaaccctgcc gagcctgcat gaatttctgc gccagcagca
13141 tggtcgttaa ggatcctcga ccaggcatca aataaaacga aaggctcagt cgaaagactg
13201 ggcctttcgt tttatctgtt gtttgtcggt gaacgctctc tactagagtc acactggctc
13261 accttcgggt gggcctttct gcgtttatat gttccgggta cagagacc
//

#### pTU2S-b(VioAE)-VioBCD

LOCUS Exported File 10343 bp ds-DNA circular SYN 02-APR-2016

DEFINITION .

ACCESSION .

VERSION .

KEYWORDS pTU2S-b(VioAE)-VioBCD

SOURCE synthetic DNA construct

ORGANISM synthetic DNA construct

REFERENCE 1 (bases 1 to 10343)

AUTHORS .

TITLE Direct Submission

JOURNAL Exported 19 Jan 2019 from SnapGene 1.1.3

http://www.snapgene.com

FEATURES Location/Qualifiers

source 1..10343

/organism="synthetic DNA construct"

/mol_type="other DNA"

misc_feature 94..682

/note="pMB1 Origin"

/note="color: #993300"

CDS complement(980..1639)

/codon_start=1

/note="CmR"

/note="color: #ccffcc"

/translation="MEKKITGYTTVDISQWHRKEHFEAFQSVAQCTYNQTVQLDITAFL

KTVKKNKHKFYPAFIHILARLMNAHPEFRMAMKDGELVIWDSVHPCYTVFHEQTETFSS

LWSEYHDDFRQFLHIYSQDVACYGENLAYFPKGFIENMFFVSANPWVSFTSFDLNVANM

DNFFAPVFTMGKYYTQGDKVLMPLAIQVHHAVCDGFHVGRMLNELQQYCDEWQGGA"

misc_feature 1765..1768

/note="Scar"

/note="color: #80ff00"

misc_feature 1777..1811

/note="BBa_J23114 promoter"

/note="color: #00ff00"

misc_feature 1816..1838

/note="pET_RBS"

/note="color: #0000ff"

misc_feature 1845..3098

/note="VioA"

/note="color: #00ffff"

misc_feature 3109..3237

/note="BBa_B0015"

/note="color: #ff0000"

misc_feature 3250..3284

/note="BBa_J23114 promoter"

/note="color: #00ff00"

misc_feature 3289..3311

/note="pET_RBS"

/note="color: #0000ff"

```
misc_feature    3318..3890
                /note="VioE"
                /note="color: #8080ff"
misc_feature    3901..4029
                /note="BBa_B0015"
                /note="color: #ff0000"
misc_feature    4057..4062
                /note="BBa Prefix"
                /note="color: #ff00ff"
misc_feature    4063..4068
                /note="BsaI"
                /note="color: #ff0000"
misc_feature    4070..4073
                /note="Scar"
                /note="color: #80ff00"
misc_feature    4082..4116
                /note="Promoter Library"
                /note="color: #00ff00"
misc_feature    4121..4154
                /note="RBS Library"
                /note="color: #0000ff"
misc_feature    4161..7154
                /note="VioB"
                /note="color: #00ff80"
misc_feature    7165..7293
                /note="Terminator Library"
                /note="color: #ff0000"
misc_feature    7306..7340
                /note="Promoter Library"
                /note="color: #00ff00"
misc_feature    7345..7378
                /note="RBS Library"
                /note="color: #0000ff"
misc_feature    7385..8671
                /note="VioC"
                /note="color: #ff80c0"
```

misc_feature 8682..8810

/note="Terminator Library"

/note="color: #ff0000"

misc_feature 8823..8857

/note="Promoter Library"

/note="color: #00ff00"

misc_feature 8862..8895

/note="RBS Library"

/note="color: #0000ff"

misc_feature 8902..10020

/note="VioD"

/note="color: #ffff80"

misc_feature 10031..10159

/note="Terminator Library"

/note="color: #ff0000"

misc_feature complement(10173..10178)

/note="BsaI"

/note="color: #00ff00; direction: LEFT"

misc_feature 10179..10184

/note="BBa Suffix"

/note="color: #ff00ff"

misc_feature 10324..10343

/note="VR Primer"

/note="color: #00ffff"

ORIGIN

```
   1 atccacagaa tcaggggata acgcaggaaa gaacatgtga gcaaaaggcc agcaaaaggc
  61 caggaaccgt aaaaaggccg cgttgctggc gtttttccac aggctccgcc cccctgacga
 121 gcatcacaaa aatcgacgct caagtcagag gtggcgaaac ccgacaggac tataaagata
 181 ccaggcgttt ccccctggaa gctccctcgt gcgctctcct gttccgaccc tgccgcttac
 241 cggatacctg tccgcctttc tcccttcggg aagcgtggcg ctttctcata gctcacgctg
 301 taggtatctc agttcggtgt aggtcgttcg ctccaagctg ggctgtgtgc acgaaccccc
 361 cgttcagccc gaccgctgcg ccttatccgg taactatcgt cttgagtcca acccggtaag
 421 acacgactta tcgccactgg cagcagccac tggtaacagg attagcagag cgaggtatgt
 481 aggcggtgct acagagttct tgaagtggtg gcctaactac ggctacacta gaagaacagt
 541 atttggtatc tgcgctctgc tgaagccagt taccttcgga aaaagagttg gtagctcttg
 601 atccggcaaa caaaccaccg ctggtagcgg tggttttttt gtttgcaagc agcagattac
 661 gcgcagaaaa aaaggatctc aagaagatcc tttgatcttt tctacggggt ctgacgctca
 721 gtggaacgaa aactcacgtt aagggatttt ggtcatgaga ttatcaaaaa ggatcttcac
 781 ctagatcctt ttaaattaaa aatgaagttt taaatcaatc taaagtatat atgagtaaac
 841 ttggtctgac agctcgaggc ttggattctc accaataaaa aacgcccggc ggcaaccgag
 901 cgttctgaac aaatccagat ggagttctga ggtcattact ggatctatca acaggagtcc
 961 aagcgagctc gatatcaaat tacgccccgc cctgccactc atcgcagtac tgttgtaatt
1021 cattaagcat tctgccgaca tggaagccat cacaaacggc atgatgaacc tgaatcgcca
```

1081 gcggcatcag caccttgtcg ccttgcgtat aatatttgcc catggtgaaa acgggggcga
1141 agaagttgtc catattggcc acgtttaaat caaaactggt gaaactcacc cagggattgg
1201 ctgacacgaa aaacatattc tcaataaacc ctttagggaa ataggccagg ttttcaccgt
1261 aacacgccac atcttgcgaa tatatgtgta gaaactgccg gaaatcgtcg tggtattcac
1321 tccagagcga tgaaaacgtt tcagtttgct catggaaaac ggtgtaacaa gggtgaacac
1381 tatcccatat caccagctca ccgtctttca ttgccatacg aaattccgga tgagcattca
1441 tcaggcgggc aagaatgtga ataaaggccg gataaaactt gtgcttattt ttctttacgg
1501 tctttaaaaa ggccgtaata tccagctgaa cggtctggtt ataggtacat tgagcaactg
1561 actgaaatgc ctcaaaatgt tctttacgat gccattggga tatatcaacg gtggtatatc
1621 cagtgatttt tttctccatt ttagcttcct tagctcctga aaatctcgat aactcaaaaa
1681 atacgcccgg tagtgatctt atttcattat ggtgaaagtt ggaacctctt acgtgcccga
1741 tcaactcgag tgccacctga cgtcctatat ctctatttta tggctagctc agtcctaggt
1801 acaatgctag cgtactttaa ctttaagaag gagatataca tatgaaacat tcttccgata
1861 tctgcattgt tggtgctggt atttctggtt tgacgtgcgc aagccatctg ctggacagcc
1921 cggcatgccg tggtctgagc ctgcgtatct ttgacatgca gcaagaagcc ggtggccgta
1981 tccgcagcaa aatgctggat ggtaaggcaa gcattgaact gggcgcaggt cgctactccc
2041 ctcagttgca cccgcatttc caaagcgcaa tgcagcacta tagccaaaag agcgaagtct
2101 atccgttcac ccagttgaag ttcaaatctc acgtgcagca aaagctgaag cgcgccatga
2161 atgaactgtc cccgcgtctg aaagagcatg gtaaagagag ctttttgcag tttgtcagcc
2221 gttatcaagg tcacgatagc gcggttggta tgatccgctc tatgggttac gacgcactgt
2281 tcctgccgga tatcagcgca gaaatggcct acgacattgt gggtaagcac ccggagatcc
2341 agagcgtgac ggacaacgac gcgaaccaat ggtttgcagc ggaaacgggc tttgctggtc
2401 tgattcaggg catcaaggct aaggttaagg cggcaggtgc gcgttttagc ctgggttatc
2461 gtctgctgag cgtccgtacc gacggtgacg gctacctgct gcaactggca ggtgacgacg
2521 gctggaaact ggagcaccgt acccgccatc tgattctggc gattccgccg agcgcgatgg
2581 cgggtttgaa tgttgatttt ccagaagcct ggtccggtgc gcgctatggc agcctgccgc
2641 tgtttaaggg ctttctgacg tacggtgagc cgtggtggtt ggactacaaa ctggacgatc
2701 aggtgctgat tgttgacaac ccgctgcgca aaatctattt caaaggcgat aagtacctgt
2761 tcttctatac cgatagcgag atggcgaatt actggcgcgg ttgtgtcgcg gagggcgagg
2821 acggttacct ggagcaaatt cgcacccatt tggctagcgc actgggtatc gtccgtgaac
2881 gtatcccgca accgctggca cacgttcaca agtattgggc gcacggcgtt gagttttgcc
2941 gtgattctga tattgaccac ccgagcgcac tgtctcatcg cgacagcggt atcatcgcgt
3001 gctccgatgc gtacacggag cattgtggtt ggatggaggg cggtctgctg agcgcccgtg
3061 aggcaagccg tctgctgttg cagcgtatcg ccgcgtgagg atcctcgacc aggcatcaaa
3121 taaaacgaaa ggctcagtcg aaagactggg cctttcgttt tatctgttgt ttgtcggtga
3181 acgctctcta ctagagtcac actggctcac cttcgggtgg gcctttctgc gtttatatgt
3241 ttgccctatt ttatggctag ctcagtccta ggtacaatgc tagcgtactt taactttaag
3301 aaggagatat acatatggag aaccgtgagc caccactgtt gccagcccgt tggagcagcg
3361 cctatgtctc ttattggagc ccgatgctgc cggatgacca gctgaccagc ggctattgct
3421 ggttcgacta tgaacgtgac atctgtcgta ttgacggcct gttcaatccg tggagcgagc
3481 gtgatactgg ttatcgcctg tggatgtcgg aggttggtaa tgcggccagc ggccgtacct
3541 ggaaacaaaa agtcgcctat ggtcgtgagc gtaccgccct gggtgaacag ctgtgtgagc
3601 gtccgctgga tgatgagact ggccctttg ccgaattgtt cctgccacgc gatgtcctgc
3661 gccgtctggg tgcccgtcac attggccgtc gcgtggttct gggtcgcgaa gcggacggtt
3721 ggcgttacca gcgcccaggt aaaggtccga gcaccctgta cctggatgcg gcgagcggca
3781 ctccactgcg catggtcacc ggcgatgaag cgtcgcgtgc aagcctgcgt gattttccga
3841 atgtgagcga ggcggagatc ccggacgcgg ttttcgcggc caagcgctaa ggatcctcga
3901 ccaggcatca aataaaacga aaggctcagt cgaaagactg ggcctttcgt tttatctgtt
3961 gtttgtcggt gaacgctctc tactagagtc acactggctc accttcgggt gggcctttct
4021 gcgtttatat gttccgggta cgaattcgcg gccgcttcta gaggtctcac tatatctcta
4081 tytkayrgct agctcagycc twggkaywrt gctagcgtac agatctaata attttgttta
4141 actttrrrrr ratacatatg agcattctgg atttcccgcg tatccacttc cgtggctggg
4201 cccgtgtcaa tgcgccgacc gcgaaccgcg atccgcacgg ccacatcgat atggccagca
4261 ataccgtggc gatggcgggt gagccgttcg acctggcacg ccatcctacg gagttccacc
4321 gtcacctgcg ctccctgggt ccgcgcttcg gcttggatgg tcgtgctgac ccggaaggcc
4381 cgttcagcct ggccgagggc tacaacgctg ccggtaacaa ccacttttcg tgggagagcg
4441 caaccgttag ccacgtgcaa tgggatggcg gtgaggcgga tcgtggtgac ggtctggtcg
4501 gtgctcgttt ggcactgtgg ggtcactaca atgattatct gcgtaccacc ttcaatcgtg
4561 ctcgttgggt cgacagcgac ccgacgcgcc gtgacgctgc acaaatctat gcgggccaat
4621 tcaccattag cccggctggt gccggtccgg gtacgccgtg gctgtttacg gcagacattg

4681 atgatagcca tggtgcacgt tggacgcgtg gcggccacat tgcagagcgt ggcggccact
4741 tcttggatga agagtttggt ctggcacgcc tgtttcagtt ctctgtgccg aaagatcacc
4801 cacattttct gtttcacccg ggtccgtttg attccgaggc ctggcgtcgt ctgcaattgg
4861 ctctggagga tgacgacgtt ctgggtctga ccgtgcaata tgcgttgttc aatatgagca
4921 ccccgcctca gccgaacagc ccggtttttc acgatatggt cggtgttgtc ggtctgtggc
4981 gtcgtggtga actggcgagc tacccggctg gtcgtctgct gcgtccgcgt caaccgggtc
5041 tgggtgacct gaccctgcgc gtcaacggtg gtcgcgttgc gctgaatttg gcgtgtgcca
5101 ttccgttcag cactcgtgcc gcgcagccaa gcgcaccgga ccgcctgacc ccggacctgg
5161 gtgccaaact gccgctgggc gatctgctgc tgcgtgatga ggacggcgca ctgttggcac
5221 gtgtgccgca ggctctgtac caagactatt ggacgaatca cggtattgtg gacctgccgc
5281 tgctgcgcga accgcgtggt agcttgaccc tgagcagcga actggcggag tggcgtgagc
5341 aagactgggt cacccaaagc gacgcgtcta acctgtacct ggaggcaccg gatcgccgtc
5401 acggtcgctt tttccctgag agcatcgcgc tgcgcagcta ctttcgcggt gaagcgcgtg
5461 cgcgtccgga tatcccgcat cgtatcgagg gcatgggcct ggtcggcgtc gaatctcgtc
5521 aggatggcga cgctgcggaa tggcgtctga cgggtctgcg tccgggtccg gcacgcattg
5581 ttctggacga tggtgccgag gcgatccctc tgcgtgttct gcctgacgat tggcgctgg
5641 atgacgcgac cgtcgaagaa gtggattacg ccttttgta ccgccacgtt atggcgtatt
5701 acgagctggt gtatccattc atgagcgaca aggtgttttc cctggctgat cgttgcaaat
5761 gtgaaacgta cgcacgtctg atgtggcaga tgtgtgatcc gcagaaccgc aacaagtcct
5821 attacatgcc gagcacccgc gaactgtcgg caccgaaagc tcgtttgttc ttgaagtatc
5881 tggcccacgt ggaaggccag gcacgcctgc aagcacctcc gccagcgggt ccggcacgca
5941 ttgaatctaa agcccagttg gcggcagagc tgcgtaaagc cgtcgacctg gagctgtctg
6001 tgatgctgca atacctgtac gcggcgtata gcattccgaa ctatgcacag ggccaacaac
6061 gtgttcgtga cggtgcgtgg accgccgagc agctgcaact ggcgtgcggt agcggtgacc
6121 gtcgccgtga tggcggtatt cgtgcagcac tgctggaaat tgctcatgaa gaaatgattc
6181 attacctggt cgttaacaac ctgctgatgg ccctgggcga gccgttctac gcgggtgtcc
6241 cgctgatggg cgaagcggca cgtcaggcgt ttggcctgga caccgagttc gctctggaac
6301 cgtttagcga aagcacgctg gcacgttttg ttcgtctgga atggccgcac tttatcccag
6361 caccgggcaa atccatcgcg gactgctatg ccgccattcg tcaggcgttt ttggatctgc
6421 cggacttgtt tggtggcgag gcaggtaagc gtggcggtga acaccacctg ttcctgaatg
6481 agctgaccaa ccgtgcgcat ccgggttatc aactggaagt tttcgatcgc gactcggcgc
6541 tgtttggtat tgcatttgtg accgatcagg gcgaaggtgg cgctctggac agcccgcact
6601 acgaacatag ccattttcaa cgtctgcgtg aaatgagcgc gcgtatcatg gctcaaagcg
6661 caccgttcga accggcgctg ccggcgttgc gtaatccggt tctggatgag agcccgggtt
6721 gccaacgtgt cgcagacggt cgtgcgcgtg cgctgatggc attgtaccaa ggcgtttatg
6781 agctgatgtt tgcgatgatg gcgcagcact tcgccgtgaa accgctgggt agcttgcgtc
6841 gcagccgcct gatgaacgca gcaatcgatc tgatgaccgg tctgttgcgt ccgctgagct
6901 gcgcgctgat gaacctgcca agcggcatcg ccggtcgcac ggccggtccg ccgctgccgg
6961 gtccggttga cacccgtagc tatgacgact acgcgctggg ctgtcgcatg ctggcacgcc
7021 gttgcgagcg tctgctggag caggcgagca tgctggaacc gggttggctg ccggatgcgc
7081 agatggagct gctggatttc tatcgtcgcc aaatgctgga cttggcgtgc ggcaaactga
7141 gccgcgaggc ctaaggatcc tcgaccaggc atcaaataaa acgaaaggct cagtcgaaag
7201 actgggcctt tcgttttatc tgttgtttgt cggtgaacgc tctctactag agtcacactg
7261 gctcaccttc gggtgggcct ttctgcgttt atatgtttgc cctatytkay rgctagctca
7321 gycctwggka ywrtgctagc gtacagatct aataattttg tttaacttr rrrrrataca
7381 tatgaaacgt gcgattatcg ttggtggcgg cctggcgggt ggcctgaccg cgatctacct
7441 ggcgaagcgt ggctacgaag tgcacgtcgt ggagaagcgt ggtgatcctc tgcgcgatct
7501 gagctcttac gtggacgttg ttagcagccg tgcgatcggc gtgagcatga ccgttcgtgg
7561 tatcaagagc gttttggctg cgggcattcc gcgtgcagag ctggatgcgt gtggcgaacc
7621 gatcgtggca atggctttct ccgtgggtgg tcagtatcgc atgcgcgaac tgaagccgtt
7681 ggaggatttc cgtccgctga gcttgaaccg tgcggcgttt caaaagctgc tgaacaaata
7741 cgcgaacctg gcaggcgttc gttactactt tgagcataag tgcctggatg ttgacctgga
7801 tggtaagagc gtgttgattc agggcaaaga tggtcagccg cagcgtctgc aaggtgacat
7861 gattatcggt gcggatggcg cccacagcgc cgtccgtcag gcgatgcaga gcggcctgcg
7921 tcgtttcgag ttccagcaaa cgttcttccg ccatggctac aaaaccctgg ttttgccgga
7981 cgcgcaagca ctgggttacc gtaaagacac gctgtacttt ttcggcatgg attccggtgg
8041 cctgttcgcg ggtcgtgcgg ctacgatccc agatggtagc gtcagcatcg ccgtttgcct
8101 gccgtactcg ggtagccctt ccctgacgac caccgacgaa ccgacgatgc gtgcgttctt
8161 cgatcgttac ttcggtggcc tgccgcgtga cgcgcgtgac gaaatgctgc gtcagtttct
8221 ggcgaagccg agcaacgacc tgattaacgt gcgctctagc acctttcact ataagggtaa

8281 tgtgctgttg ctgggtgatg ctgcgcatgc gactgcgccg ttcctgggtc agggtatgaa
8341 catggcgctg gaggacgccc gcacgtttgt cgagctgctg gaccgccacc agggcgacca
8401 agacaaagcc tttccggagt tcacggagct gcgcaaagtc caggcagacg caatgcaaga
8461 catggctcgc gccaactatg acgttttgag ctgctcgaac ccgatctttt tcatgcgtgc
8521 gcgttacacg cgttacatgc attccaagtt tccgggcctg tatccgccgg atatggccga
8581 gaaactgtac tttacgagcg agccgtacga tcgtctgcaa caaatccagc gtaaacagaa
8641 tgtttggtac aagattggtc gcgtgaattg aggatcctcg accaggcatc aaataaaacg
8701 aaaggctcag tcgaaagact gggcctttcg ttttatctgt tgtttgtcgg tgaacgctct
8761 ctactagagt cacactggct caccttcggg tgggcctttc tgcgtttata tgttccggct
8821 atytkayrgc tagctcagyc ctwggkaywr tgctagcgta cagatctaat aattttgttt
8881 aactttrrrr rratacatat gaagattctg gtcattggtg ctggtccagc tggtctggtt
8941 ttcgcatccc aactgaagca ggcacgccct ttgtgggcca ttgacatcgt ggagaagaat
9001 gacgagcaag aagtgctggg ctggggtgtc gtgctgcctg gccgtccggg tcagcacccg
9061 gcgaacccgc tgtcctatct ggatgcaccg gagcgtctga atccgcaatt tctggaggac
9121 ttcaaactgg tgcatcataa tgagccgtcc ttgatgtcca cgggcgtttt gttgtgcggc
9181 gtggagcgtc gcggtctggt tcacgcgctg cgcgataagt gccgcagcca aggcattgct
9241 attcgtttcg aaagcccgtt gctggaacac ggtgagctgc cgctggcgga ctatgatctg
9301 gtggtcctgg ctaatggtgt taatcacaaa accgcgcatt tcaccgaggc tctggtcccg
9361 caggtggact acggccgcaa taagtacatt tggtatggca ctagccagct gttcgatcag
9421 atgaatctgg ttttcgtac ccatggtaaa gatatcttta tcgcgcatgc ctataagtat
9481 agcgatacca tgagcacgtt cattgtcgaa tgtagcgaag agacttacgc acgcgcacgc
9541 ctgggcgaaa tgtccgaaga ggcgagcgca gaatacgttg ctaaggtgtt ccaggccgag
9601 ctgggtggtc acggcctggt gagccagccg ggtctgggtt ggcgtaactt catgacgttg
9661 tctcatgacc gttgtcatga tggtaagttg gttctgctgg gtgacgcgct gcaaagcggt
9721 cactttagca tcggccacgg caccacgatg gccgtggtgg tggcgcagct gctggttaaa
9781 gcgctgtgta ccgaagatgg tgtgcctgcc gcgctgaaac gtttcgaaga gcgtgccctg
9841 ccgctggtgc agttgttccg tggccacgca gacaacagcc gcgtttggtt cgaaaccgtc
9901 gaagagcgca tgcacctgtc ctcggcggaa tttgtgcaaa gcttcgacgc acgccgcaaa
9961 agcctgccgc cgatgccgga agcactggcg cagaatctgc gttatgcttt gcagcgctga
10021 ggatcctcga ccaggcatca aataaaacga aaggctcagt cgaaagactg ggcctttcgt
10081 tttatctgtt gtttgtcggt gaacgctctc tactagagtc acactggctc accttcgggt
10141 gggcctttct gcgtttatat gttgaaggta cagagaccct gcagtccggc aaaaaagggc
10201 aaggtgtcac caccctgccc tttttcttta aaaccgaaaa gattacttcg cgttatgcag
10261 gcttcctcgc tcactgactc gctgcgctcg gtcgttcggc tgcggcgagc ggtatcagct
10321 cactcaaagg cggtaatacg gtt
//
